# The glycine betaine-cobalamin feedback loop drives cross-feeding between marine bacteria and algae

**DOI:** 10.64898/2025.12.19.695462

**Authors:** Jonathan Hammer, Myriel Staack, Martin Sperfeld, Tom Haufschild, Nicolai Kallscheuer, Delia A. Narváez-Barragán, Carl-Eric Wegner, Kirsten Küsel, Shinichi Sunagawa, Georg Pohnert, Torsten Schubert, Einat Segev, Christian Jogler

## Abstract

Heterotrophic bacteria supply cobalt-containing cobalamin (Cbl) to marine microalgae, which in return provide organic substrates. Such metabolic cross-feeding can regulate the species composition of prolific marine plankton and biofilms. Algal-produced glycine betaine (GB) can be catabolized by bacteria using a Cbl-dependent demethylase (MtgBCD). Yet, GB’s impact on bacterial Cbl production during cross-feeding, and its control by cobalt scarcity remains elusive. Here, we demonstrate that *Phaeobacter inhibens* bacteria boost their Cbl production 25-fold when grown in monocultures with GB compared to glucose. During co-cultivation, *P. inhibens* satisfied the Cbl requirements of *Gephyrocapsa huxleyi* algae. Transcriptomic analysis of mono- and co-cultures revealed co-expression of *mtgBCD* and *gbcAB* genes encoding Cbl-dependent and -independent GB demethylases, respectively. Distinct growth defects of deletion mutants indicate that *P. inhibens* switches from Cbl-dependent to -independent GB demethylation under cobalt limitation, involving a Cbl riboswitch. Our findings suggest a positive feedback loop in which algal GB release stimulates the bacterial supply of Cbl. We predict the breakdown of this interaction under naturally occurring cobalt limitation which potentially contributes to the transient nature of algal blooms. Comparative genomics indicate that this mechanism is widespread in *Rhodobacterales* and other abundant marine *Alphaproteobacteria*, underscoring its pivotal role in global ocean productivity.

## Introduction

The ocean annually absorbs about a quarter of anthropogenic CO_2_ emissions^1,2^. Marine microalgae and cyanobacteria contribute to this CO_2_ sink by using sunlight to build up biomass from dissolved CO_2_. They account for approximately half of the annual net primary production on earth^3^. Under favorable environmental conditions, microalgae like *Gephyrocapsa huxleyi* (formerly *Emiliania huxleyi*) can form large transient blooms. These blooms are strongly shaped by interactions between the algae and associated heterotrophic bacteria which exchange metabolites that influence bloom dynamics and biogeochemical processes^4–6^.

*Phaeobacter inhibens* and other members of the order *Rhodobacterales* can dominate the algal microbiota and naturally co-occur with *G. huxleyi* blooms^7–12^. *G. huxleyi* is frequently co-cultivated with *P. inhibens* to study interaction dynamics^7,13–17^. In these co-cultures, *P. inhibens* shows a “Jekyll and Hyde” lifestyle characterized by an initial mutualistic phase, followed by an antagonistic phase leading to algal demise^7^. Some other *Rhodobacterales* show similar behaviors in co-cultures with algae^18–20^. During mutualism, *Rhodobacterales* grow on organic compounds released by the algae and in return promote algal growth by the production of phytohormones and vitamins^7,21–24^. Especially important is the supply with the cobalt-containing co-factor cobalamin (vitamin B_12_, Cbl), as roughly half of all algae depend on it, yet cannot synthesize it *de novo*^25^. Auxotrophic algae only possess the Cbl-dependent methionine synthase MetH while the Cbl-independent methionine synthase MetE is absent^26^. In the ocean, presence or absence of Cbl can alter microbial community compositions and the primary productivity of microalgae^27–30^, thus establishing Cbl as a marine keystone metabolite^6^. Cbl dependency of *G. huxleyi* is indicated by the lack of *metE* but controversially debated^26,31,32^.

Glycine betaine (trimethylglycine, GB) is a compatible solute produced by many microalgae and bacteria^33–37^ and a dominant marine metabolite^6^. It can serve as a carbon-, nitrogen- and energy source for bacteria that live associated with phototrophs^14,15,38,39^. Recently, it has been shown that *G. huxleyi* shares GB with *P. inhibens* during mutualistic growth of the co-culture, but increasing GB release induces H_2_O_2_ production of *P. inhibens*, which accelerates the population collapse of *G. huxleyi*^14,15^. Algal methylated compounds like GB and dimethylsulfoniopropionate (DMSP) act as chemoattractants of some bacteria including *Ruegeria* sp. TM1040 (*Rhodobacterales*)^40–43^ and shorten the lag-phase of *P. inhibens* due to methyl group assimilation through the methionine cycle^14^. Recently, methionine biosynthesis and GB catabolism were shown to be connected in *Ruegeria pomeroyi* (*Rhodobacterales*) by a modular enzyme that requires Cbl as a co-factor and dually functions as GB demethylase and methionine synthase (MtgBCDE)^44^. The *mtgBCDE* genes were found in cosmopolitan marine pelagic bacteria of the *Alpha*-, *Delta*- and *Gammaproteobacteria.* Furthermore, the *mtgBCD* genes are highly expressed in *P. inhibens* during the mutualistic phase of the co-culture with *G. huxleyi*^14^.

A stimulatory effect of GB on the Cbl biosynthesis of some bacteria is known since the second half of the 20^th^ century and GB is used as an additive during the biotechnological Cbl production^45,46^. However, GB-stimulated Cbl biosynthesis has never been addressed in an ecological context and its effect on important marine Cbl producers like members of the *Rhodobacterales* is unknown^47^. Bacterial Cbl biosynthesis is tightly regulated, mainly on the post-transcriptional level, by binding of Cbl to a cobalamin riboswitch in the 5’-untranslated region (UTR) of the Cbl biosynthesis operon transcripts^46,48–50^. The resulting conformational changes of the mRNA prevent ribosome binding and thus translation. This negative feedback usually prevents the energetically costly overproduction of Cbl, which competes with the reported effect of GB-stimulated Cbl biosynthesis.

In this study, we investigate how GB demethylation affects the Cbl production in *P. inhibens* (strain DSM 17395) and thus the cross-feeding with *G. huxleyi* (strain CCMP3266). By limiting the availability of cobalt, the central ion of the Cbl corrin-ring, we control the level of Cbl produced by *P. inhibens* and show that adaptations to this condition allow growth of *P. inhibens* but transiently alter its cell morphology. Using mutant strains, we elucidate that *P. inhibens* switches from Cbl-dependent to -independent GB demethylation when cobalt availability is limited. Comparative transcriptomics suggest that GB-induced changes in *P. inhibens’* metabolism also affect the redox status of the cell. We present a mechanistic model for how growth on GB enables Cbl overproduction leading to a positive feedback loop of algal GB biosynthesis and bacterial Cbl production. Based on (pan-)genomic analyses we predict that this mechanism is widespread in members of the *Rhodobacterales* and other abundant marine alphaproteobacterial orders.

## Results

### Co-expression of cobalamin-dependent and -independent glycine betaine demethylases

*P. inhibens* can grow on GB as sole carbon and energy source like other marine heterotrophic bacteria that use Cbl-dependent or -independent enzymes for GB demethylation^15,38,44,51^. In *P. inhibens*, we found three different enzymes that putatively demethylate GB: the GB--homocysteine *S*-methyltransferase MtgE, also referred to as Bmt or Bhmt^39,52^, the heterodimeric GB monooxygenase GbcAB encoded as a two-gene operon^53–55^ and the modular Cbl-dependent GB demethylase^44,56^. The latter consists of a GB--Cbl methyltransferase (MtgB), a Cbl-binding protein (MtgC) and a Cbl--tetrahydrofolate methyltransferase (MtgD). The previously reported split methionine synthase^57^ and modular GB demethylase^44^ share MtgC and MtgD (Fig. 1). MtgE acts as GB--homocysteine methyltransferase or as the Cbl--homocysteine methyltransferase module of the split methionine synthase, depending on the binding to MtgCD^52,57^. We performed a differential gene expression analysis using RNA sequencing (RNA-Seq) data from *P. inhibens* cultures grown on GB or glucose as sole carbon and energy source to investigate how the expression of the different demethylation-related genes is affected (Fig. 1). Of the genes involved in GB and one-carbon metabolism, 21 ranked among the top 100 upregulated and highly expressed genes (Supplementary Data S1). Of the four trimethylamine methyltransferase family protein-encoding genes (*mtgB1-4*), three were upregulated in the presence of GB, but only the *mtgB1* transcript was highly abundant alongside *mtgC* and *mtgD* (Fig. 1 and Supplementary Data S1). In contrast, *mtgE* was downregulated. By comparing the transcriptional ratios based on mean expression levels (normalized as transcripts per million, TPM) of *mtgE*, *mtgB1-4*, and *mtgC* across RNA-Seq data of *P. inhibens* (including co-cultures with *G. huxleyi*^14^), we noticed that the presence of methyl group donors such as GB or DMSP induced a shift in the ratios, displacing the methionine synthesis-associated *mtgE* transcripts (Fig. 1). In contrast, during growth of *P. inhibens* in monocultures without methyl group donors, the demethylation associated *mtgB1-4* transcripts were displaced. This suggests substrate-induced changes in gene expression levels that dictate the formation of the competing Cbl-dependent GB demethylase (MtgBCD) and methionine synthase (MtgECD) complexes. However, not only *mtgBCD* were highly expressed during growth on GB but also *gbcAB*, suggesting co-expression of genes encoding Cbl-dependent and -independent GB demethylases (Fig. 1 and Supplementary Data S1).

**Figure 1:**
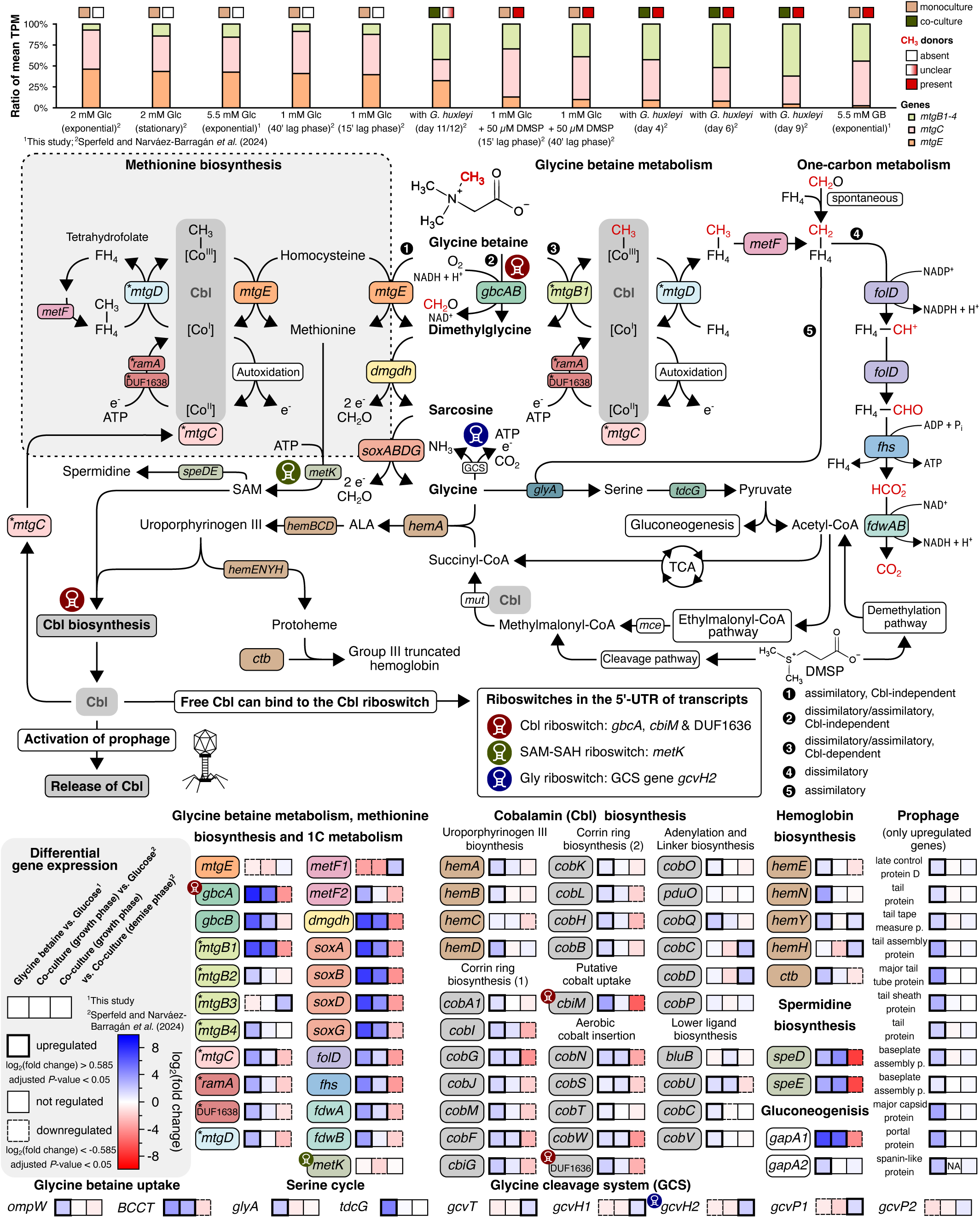
Gene expression patterns of cobalamin-dependent and -independent glycine betaine metabolism as well as methionine, cobalamin, hemoglobin and spermidine biosynthesis in *P. inhibens* during growth with glycine betaine and during co-cultivation with *G. huxleyi*. GB can be demethylated via a modular cobalamin-dependent methyltransferase system (MtgBCD) that shares modules with the split methionine synthase (MtgECD). **Upper panel**: Transcriptional ratios of *mtgB1-4* (green), *mtgC* (light red) and *mtgE* (orange) differ between conditions in which methyl group donors are absent (white boxes) or present (red boxes) for both, *P. inhibens* monocultures (brown boxes) or co-cultures with *G. huxleyi* (green boxes). **Metabolic map:** The anabolic split methionine synthase (grey box with dashed frame) and the catabolic Cbl-dependent GB demethylase differ by one module encoded by *mtgE* or *mtgB1*, respectively. Oxidation states of the cobalt ion during the Cbl-dependent methyl transfer reactions are displayed (dark grey box). Alternatively, GB can be demethylated by a heterodimeric monooxygenase GbcAB. The pathways that link GB demethylation to the dissimilatory and assimilatory one-carbon metabolism and to the biosynthesis pathways of Cbl, heme/hemoglobin and spermidine are displayed. Furthermore, pathways that link DMSP demethylation to Cbl biosynthesis are shown. Riboswitches in the 5’-UTR of transcripts involved in the displayed pathways are highlighted. Asterisks highlight genes only present in *mtgC*-containing *Rhodobacteriales* genomes (36 of 40). **Lower panel:** In three boxes, gene expression changes are displayed as log_2_(fold change): box 1) *P. inhibens* cultures with GB compared to glucose, box 2) *P. inhibens-G. huxleyi* co-cultures during the growth phase (days 4, 6 and 9) compared to *P. inhibens* monocultures with glucose^14^, box 3) co-cultures in the demise phase (day 11/12) compared to the growth phase^14^. Colors reflect upregulation (blue), no regulation (white) and downregulation (red) and the frame (bold, normal or dashed line) reflects significance as indicated in the legend (grey box in the bottom left corner). Gene accession numbers, functional annotations and log_2_(fold change) values are listed in Supplementary Tables S1 and S2. Glc, Clucose; FH_4_, tetrahydrofolate; GCS, glycine cleavage system; TCA, tricarboxylic acid cycle; SAM, *S*-adenosylmethionine; SAH, *S*-adenosylhomocysteine; NA, not applicable

### Glycine betaine induces gene expression changes affecting cobalamin biosynthesis and low-oxygen adaption in *P. inhibens*

In addition to the potential involvement of Cbl as a co-factor, GB catabolism is linked to Cbl biosynthesis as sequential demethylation yields glycine, which — together with succinyl-CoA — serves as substrate for the aminolevulinate synthase HemA (Fig. 1). Formation of aminolevulinic acid by HemA represents the first committed step towards the biosynthesis of uroporphyrinogen III, the shared precursor of Cbl and heme biosynthesis. Thus, we analyzed whether GB affects the expression of Cbl biosynthesis genes in *P. inhibens* (Fig. 1). During growth on GB, *hemA* ranked among the top 100 upregulated and highly expressed genes (Supplementary Data S1). In contrast genes involved in glycine oxidation via the glycine cleavage system (GCS) were downregulated (Fig. 1). Together with the upregulation of the serine hydroxymethyltransferase gene (*glyA*), this suggests that glycine utilization for aminolevulinic acid biosynthesis and for one-carbon assimilation via the serine cycle is prevailing. Two of three genes related to uroporphyrinogen III biosynthesis were also upregulated. Furthermore, many Cbl biosynthesis genes, including all genes involved in corrin ring biosynthesis and cobalt insertion, two of six genes encoding enzymes involved in adenylation and linker biosynthesis and three of four genes encoding enzymes involved in lower ligand biosynthesis were upregulated. A gene annotated as energy-coupling factor ABC transporter permease had the highest expression level of all upregulated genes during cultivation with GB (Supplementary Data S1). It encodes a protein of the CbiM/NikMN family that is putatively involved in cobalt uptake^58^ and is located in a large Cbl biosynthesis gene cluster (Supplementary Fig. S1). Of the two Cbl riboswitches in this cluster, one is situated in the 5’-UTR of the *cbiM* transcript. This suggests post-transcriptional regulation of cobalt uptake and Cbl biosynthesis depending on the intracellular Cbl concentration. In addition to the Cbl biosynthesis branch, genes encoding enzymes for heme biosynthesis from uroporphyrinogen III were upregulated (Fig. 1). Our transcriptomic data suggests that GB catabolism is metabolically linked to Cbl and heme biosynthesis.

Prior to the discovery that MtgCD is essential for aerobic growth of *R. pomeroyi* on GB, Cbl-dependent catabolic methyltransferases were considered anaerobe enzymes^44,56,59–62^. For Cbl-dependent methyl transfer the cobalt(I) reduced state is required (Fig. 1). It has been reported that even under anaerobic assay conditions, cobalt(I) is oxidized to the inactive cobalt(II) state in one of 100 turnover cycles^63^. Cobalt autoxidation presumably increases with rising oxygen concentration. In *P. inhibens*, the enzymatic reactivation of Cbl likely requires ATP^57,64^. Thus, Cbl-dependent GB demethylation under aerobic conditions could be energetically inefficient. We searched the transcriptomic data for evidence of a more reducing intracellular environment, notably lower oxygen concentrations, that would limit cobalt oxidation during growth on GB. Among the top 100 upregulated and highly expressed genes in the presence of GB, we found *ccoNOQP* (*fixNOPQ*), encoding the cytochrome c oxidase *cbb_3_* (Supplementary Data S1). This terminal oxidase has a high oxygen affinity and is commonly associated with microaerophilic growth conditions^65–67^. Genes encoding the oxygen-dependent and -independent coproporphyrinogen III oxidases, *hemF* and *hemN*, were also upregulated with log_2_-(fold change) values of 0.713 and 3.605, respectively. This suggests that presence of GB especially effects expression of *hemN* which has been shown to be repressed by oxygen in various bacteria^66,68–71^. In addition to changes in the expression of heme biosynthesis genes, we observed upregulation of a gene encoding a group III truncated hemoglobin (*ctb*). The only characterized homologue is involved in moderating the oxygen fluxes in the microaerophilic *Campylobacter jejuni*^72,73^. Furthermore, the *fixAB* genes encoding a heterodimeric electron transfer flavoprotein (ETF), were among the top 10 upregulated and highly expressed genes (Supplementary Data S1). In mitochondria, oxidized ETFs instead of O_2_ serve as acceptors for electrons from dimethylglycine and sarcosine demethylation by dehydrogases^74,75^. The bacterial ETFs have been frequently reported to be involved in anaerobic or microaerophilic^76–80^ as well as methylotrophic metabolism^81,82^. From reduced ETFs the electrons can enter the electron transport chain by the ETF-ubiquinone oxidoreductase reaction. The gene putatively encoding this enzyme (*fixC*) is also among the top 100 upregulated and highly expressed genes. In summary, GB triggers expression of genes typically associated with microaerophilic or anaerobic lifestyles in bacteria, despite aerobic cultivation conditions of *P. inhibens*.

### The gene expression profile of glycine betaine-fed *P. inhibens* monocultures is mirrored in co-cultures with *G. huxleyi*

To verify the ecological relevance of our RNA-Seq results, we compared them with dual RNA-Seq data previously generated for *P. inhibens* and *G. huxleyi* co-cultures^14^. This includes data for samples taken at days 4, 6 and 9 during the mutualistic growth phase and at day 11/12 during the algal demise phase. GB catabolic genes, as well as one-carbon dissimilation genes were upregulated in *P. inhibens* monocultures with GB and in algal-bacterial co-cultures during the growth phase, compared to bacterial monocultures on glucose (Fig. 1). Co-expression of Cbl-dependent (*mtgBCD*) and -independent GB demethylases (*gbcAB*) was observed in all cultures that contained GB (either added, or produced by algae), while *mtgE* (required for methionine synthesis) was always downregulated under those conditions, as mentioned earlier. Most of the Cbl biosynthesis genes were upregulated during growth on GB compared to glucose in monocultures. During the growth phase of the co-cultures, some of the Cbl biosynthesis-related genes involved in corrin ring biosynthesis, cobalt insertion and lower ligand biosynthesis were upregulated, too. Since prophage activation is a proposed release mechanism for Cbl during cross-feeding^83^ and prophage genes were upregulated in mono-cultures with GB (Fig. 1), we checked expression of related genes in the RNA-Seq data generated for the co-cultures. However, prophage gene transcripts were barely detected.

To identify whether GB-related gene expression changes over time, we analyzed expression levels at different timepoints of the co-cultures. The ratio between the mean expression levels (normalized as TPM) of *mtgB1*, *mtgC*, *mtgD*, *gbcA* and *gbcB* indicates a dominance of *mtgBCD* at day 4 (early growth phase), but the relative share of *gbcAB* increased until day 9 (late growth phase) (Fig. 2a). In the monoculture of *P. inhibens* with 5.5 mM GB, the ratio between Cbl-dependent and -independent GB demethylase was almost equal. Differential gene expression analysis revealed that only 65 genes were upregulated from day 4 to day 9 of the co-culture (Supplementary Data S2), 40 of which were also upregulated in the monoculture with GB (Supplementary Data S1). This included genes related to GB and one-carbon metabolism as well as Cbl biosynthesis (Fig. 2b). While the *mtgBD* genes were upregulated, *mtgC* was highly expressed at a constant level throughout the mutualistic growth phase (day 4, 6 and 9). Mean TPM of *mtgBD* and *gbcAB* constantly increased from day 4 to day 9. The putative cobalt uptake-related gene *cbiM* was highly upregulated from day 4 to day 9. This is the expected response to increasing cobalt demands while its availability decreases. Most of the selected Cbl biosynthesis genes were not upregulated, except for the cobyrinate a,c-diamide synthase gene *cobB*. While the ETF-encoding genes (*fixAB*) were constantly high expressed, the genes related to low intracellular oxygen concentrations (*ccoNOQP* and *hemN*) were upregulated from day 4 to day 9. Interestingly, denitrification genes were highly upregulated at the same time (Supplementary Data S2). Taken together, this suggests modifications of the electron transport chain during co-cultivation. Mean TPM of all displayed genes declined sharply on day 11/12 of the co-culture after algal cell death (Fig. 2). This fits to the downregulation of GB catabolism-, one-carbon dissimilation- and Cbl biosynthesis-related genes during the demise phase of the co-culture in comparison to the growth phase (Fig. 1 and Supplementary Data S3).

**Figure 2:**
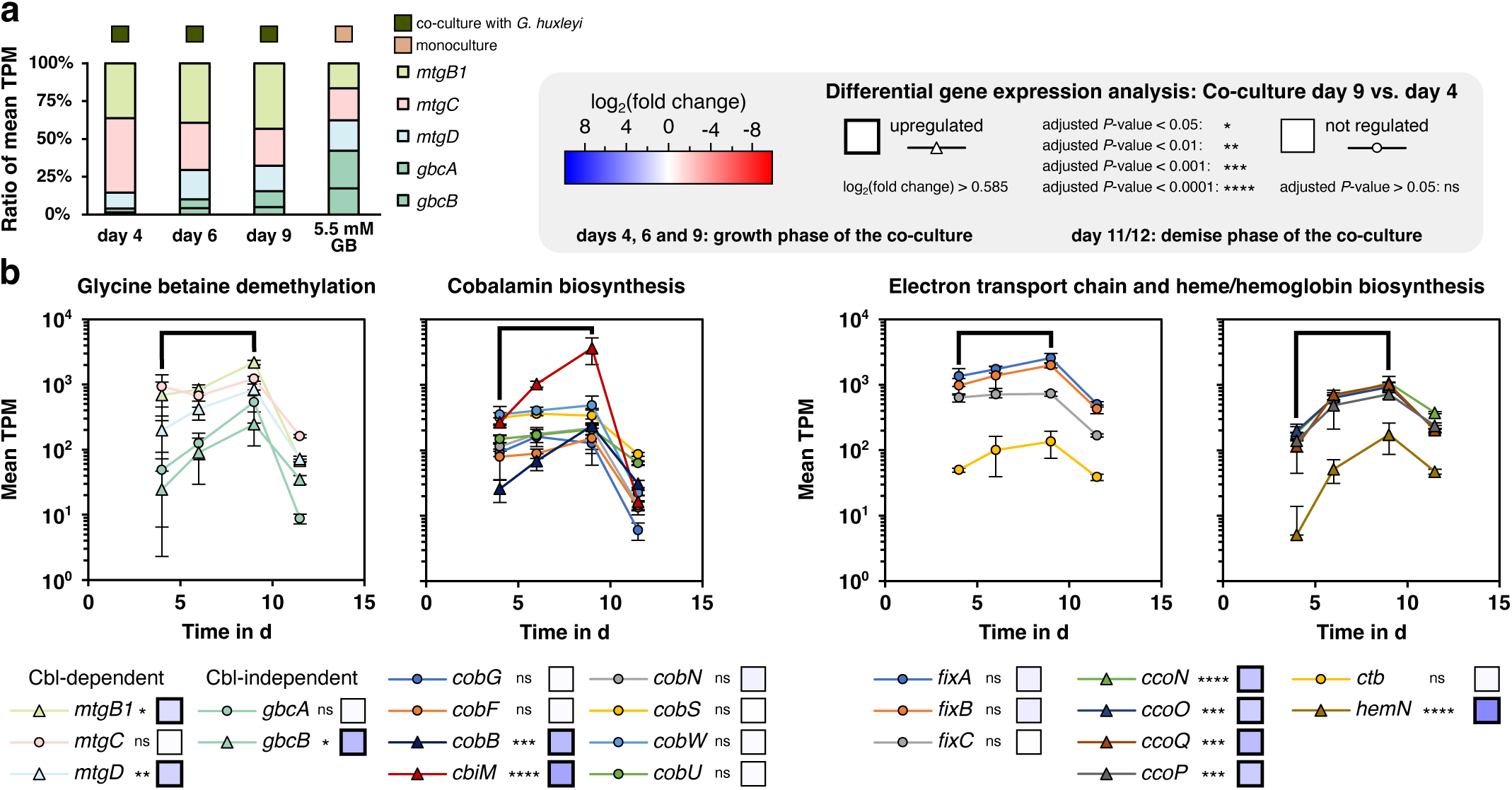
Genes involved in glycine betaine demethylation, cobalamin biosynthesis, cobalt uptake, electron transport, and heme biosynthesis are upregulated from day 4 to day 9 of the *P. inhibens*-*G. huxleyi* co-culture. **a**, Transcriptional ratios of the Cbl-dependent (*mtgBCD*) and -independent GB demethylases (*gbcAB*) are compared between day 4, 6 and 9 of the *P. inhibens-G. huxley* co-culture (green box) as well as the *P. inhibens* monoculture with 5.5 mM GB (brown box). **b,** Expression levels of selected genes during co-cultivation with *G. huxleyi* are plotted over time. Days 11/12 of the co-culture – after algal cell death – are plotted as one timepoint at 11.5 days. Co-cultivation was done in biological triplicates and error bars represent the standard deviation (n = 3). Gene expression changes as log_2_(fold change) are displayed in colored boxes (blue, upregulation; white, no regulation; red, downregulation) and refer to the *P. inhibens-G. huxleyi* co-culture on day 9 (late growth phase, immediately before induced algal cell death) compared to day 4 (early growth phase). Box frames (bold and normal) and asterisks denote significance as indicated in the legend (grey box). Genes were selected for display only when upregulated in *P. inhibens* monocultures with 5.5 mM GB in comparison to the glucose control. Additionally, Cbl biosynthesis genes were selected for display when upregulated during the growth phase of the co-culture (day 4 to 9) compared to glucose or on day 9 compared to day 4. Expression of *fixABC* indicates electron transport via ETF. The high-affinity cytochrome c oxygenase *cbb_3_* (*ccoNOQP*) and the oxygen-independent heme biosynthesis protein (*hemN*), are commonly expressed under low oxygen conditions. A group III truncated hemoglobin is encoded by *ctb*. Gene accession numbers, functional annotations and log_2_(fold change) values are listed in Supplementary Tables S1 and S2.

In summary, the gene expression profile of *P. inhibens* in GB monocultures is mirrored in the co-cultures with *G. huxleyi* – the natural GB source – while genes directly or indirectly linked to GB metabolism are progressively expressed with increasing algal cell density.

### Cobalt availability determines the enzyme used for glycine betaine demethylation

To dissect the interplay between Cbl-dependent and -independent GB demethylation in *P. inhibens*, we restricted the bacterial Cbl production by limiting the cobalt availability. In our setting, cobalt-limitation does not refer to a limiting effect on the growth of *P. inhibens*, but to a limiting effect on the Cbl production. We analyzed the cobalt concentration in our media with and without addition of CoCl_2_ via inductively coupled plasma optical emission spectroscopy (ICP-OES) and confirmed that it is below the detection limit of this method (lower than 0.02 mg l^-1^ or 0.34 µM) (See Supplementary Text on “Cobalt limitation in the context of this study” and Supplementary Fig. S2). We performed growth experiments with the *P. inhibens* wild type in comparison to single-gene deletion mutants of *mtgC*, *mtgB1*, *gbcAB*, and a double-gene deletion mutant of *mtgB1* + *gbcAB* under cobalt-replete and - limited conditions (Fig. 3a). Growth of the *P. inhibens* wild type with GB as sole carbon and energy source was not affected by cobalt limitation. Growth on glucose under cobalt-replete and -limited conditions served as controls and confirmed that the gene deletions did not lead to non-specific growth defects. The Δ*gbcAB* mutant revealed that these genes are essential for growth under cobalt-limited conditions but not for growth under cobalt-replete conditions. As expected, the Δ*mtgB1* mutant grew under cobalt-limited conditions but exhibited surprising growth defects under cobalt-replete conditions, despite the presence of a functional *gbcAB* that could rescue the lack of *mtgB1*. In line, the Δ*mtgB1* Δ*gbcAB* double mutant was unable to grow on GB under both, cobalt-replete and -limited conditions. Further, the Δ*mtgC* mutant, which lacks the Cbl-binding protein, grew only if methionine was added and showed growth dynamics similar to the Δ*mtgB1* mutant. This confirms the double role of MtgC in GB demethylation and methionine biosynthesis^44^.

**Figure 3:**
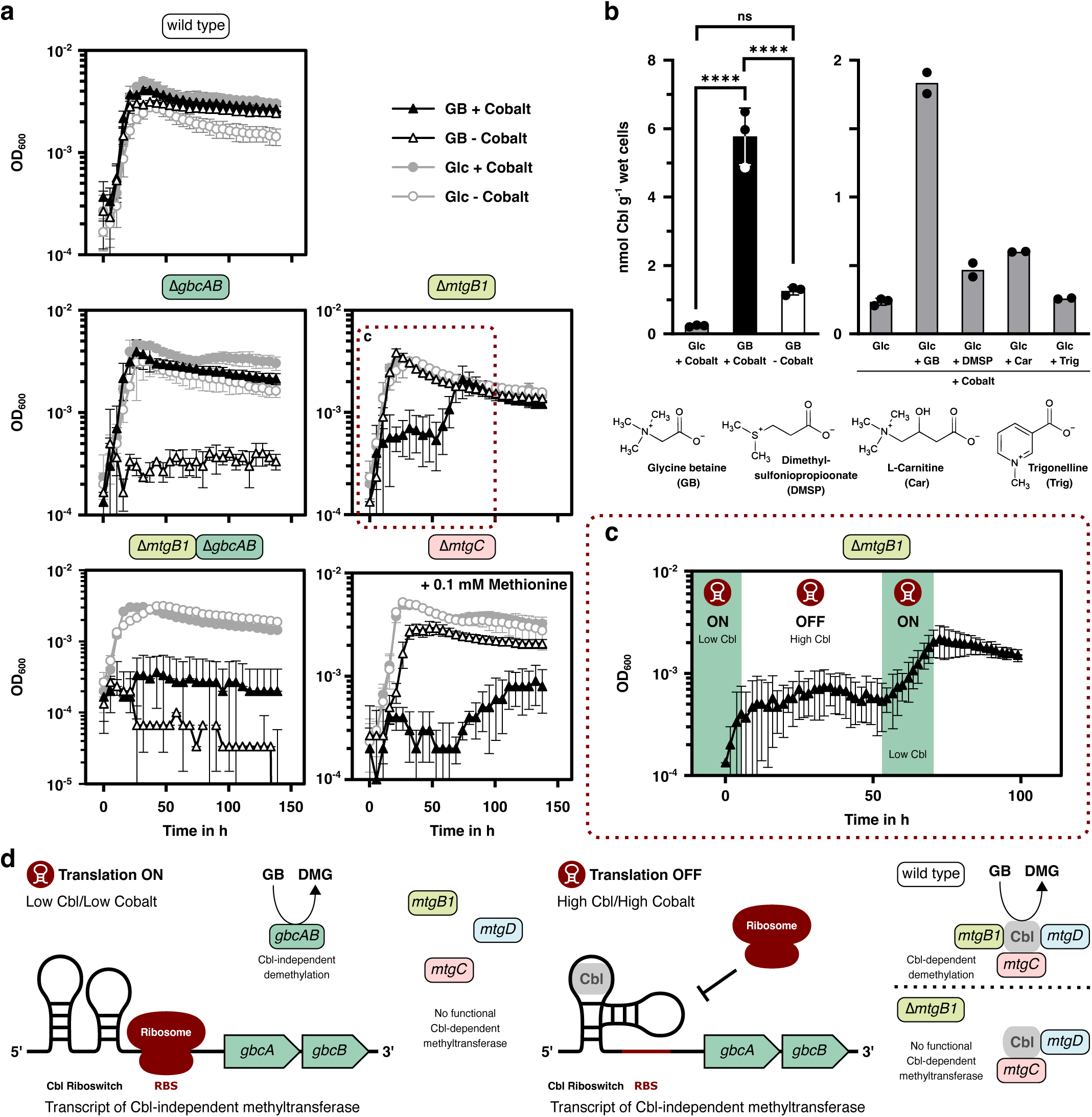
Intracellular cobalamin concentrations regulate cobalamin-dependent and - independent glycine betaine demethylation in *P. inhibens*. **a**, Growth of *P. inhibens* wild type and deletion mutants on 1 mM GB or 1 mM glucose (Glc) in medium containing (+ Cobalt) or lacking cobalt (- Cobalt). Mutants lack either the Cbl-independent monooxygenase (Δ*gbcAB*), components of the modular Cbl-dependent methyltransferase (GB--Cbl methyltransferase, Δ*mtgB1*; Cbl-binding protein, Δ*mtgC*) or both (Δ*mtgB1* Δ*gbcAB*). Since the *mtgC* deletion mutant is methionine auxotrophic, 0.1 mM methionine was added. Growth curves represent mean values of biological triplicates and error bars represent the standard deviation (n = 3). OD_600_ values are displayed without pathlength (λ) correction. In the graphs only every tenth timepoint is displayed. **b**, Quantification of the intracellular Cbl concentration of *P. inhibens* under cobalt-replete (+ Cobalt) or -limited (- Cobalt) conditions. *P. inhibens* was grown on 5.5 mM glucose or 5.5 mM GB, (left) or on 5.5 mM Glucose to which 1 mM of the *N*- or *S*-methylated compounds were added (right); their chemical structures are displayed below. Bars represent the mean value; dots represent individual measured values. The error bars represent the standard deviation (only displayed for n ≥ 3). To determine the significance of differences, ANOVA followed by Tukey’s post-hoc test was used (adjusted *P*-value > 0.05: ns, not significant; adjusted *P*-value < 0.0001: ****, significant). **c**, Deficiency phenotype of the Δ*mtgB1* mutant during growth with GB under cobalt-replete conditions. Growth phases are shaded in green. Predicted intracellular Cbl levels (high/low) and the resulting translational state of the Cbl riboswitch in the 5’-UTR of *gbcAB* (ON or OFF) are indicated. Switch between both states is a consequence of Cbl binding to the riboswitch. **d**, Proposed model of the Cbl-regulated GB demethylation involving a Cbl riboswitch. RBS, ribosome binding site; DMG, dimethylglycine

Our data demonstrates that *P. inhibens* does not use GbcAB for GB demethylation under cobalt-replete conditions despite high expression levels of the encoding genes (Fig. 1 and Supplementary Data S1), pointing towards additional post-transcriptional regulation. This agrees with a Cbl riboswitch in the 5’-UTR of the *gbcAB* transcript that likely prevents translation upon Cbl-binding. The growth curve of the Δ*mtgB1* deletion mutant on GB under cobalt-replete conditions indicates that the *P. inhibens* mutant produces Cbl in the presence of GB, despite the dependency on GbcAB for GB demethylation (Fig. 3a+c). Over time, this led to growth arrest of the Δ*mtgB1* mutant because of free Cbl binding to the Cbl riboswitch on the 5’-UTR of the *gbcAB* transcript (Fig. 3c+d). Notably, the Δ*mtgB1* mutant started growing again, which is presumably due to decreasing concentrations of free Cbl.

In conclusion, our data suggest a tight regulation that links cobalt availability to Cbl biosynthesis and GB degradation: When cobalt is depleted, Cbl biosynthesis decreases which activates Cbl-independent GB demethylation, involving a Cbl riboswitch.

### Glycine betaine and cobalt availability jointly control the cobalamin production and cell morphology of *P. inhibens*

Given the GB-induced upregulation of Cbl biosynthesis genes (Fig. 1) and the switch to Cbl-independent GB demethylation under cobalt-limitation (Fig. 3a), we quantified intracellular Cbl concentrations under cobalt-replete and -limited conditions. We found that *P. inhibens* significantly increased Cbl production by a factor of 25 if grown on GB compared to glucose under cobalt-replete conditions (Fig. 3b). In contrast, cobalt limitation led to a significant decrease in Cbl production. This is in line with the observed switch from Cbl-dependent to - independent GB demethylation. The difference between the Cbl concentrations during growth on GB under cobalt limitation and during growth on glucose under cobalt-replete conditions is not statistically significant (adjusted *P*-value = 0.0915). However, the mean is still 5.4-fold as high suggesting a potential biological effect. Strikingly, neither DMSP and trigonelline – methylated compounds produced by algae^35^ – nor the structurally similar carnitine had a comparable effect (Fig. 3b). This highlights the exceptional role of GB as a stimulant of bacterial Cbl biosynthesis.

During routine microscopic controls, we observed a transient elongation of *P. inhibens* cells during cobalt depletion (Fig. 4), presumably resulting from impaired cell division. Collectively, our data show that cobalt limitation suppresses Cbl biosynthesis, triggers nutrient stress-induced filamentation, and that this morphological shift is reversed by adaptive responses.

**Figure 4:**
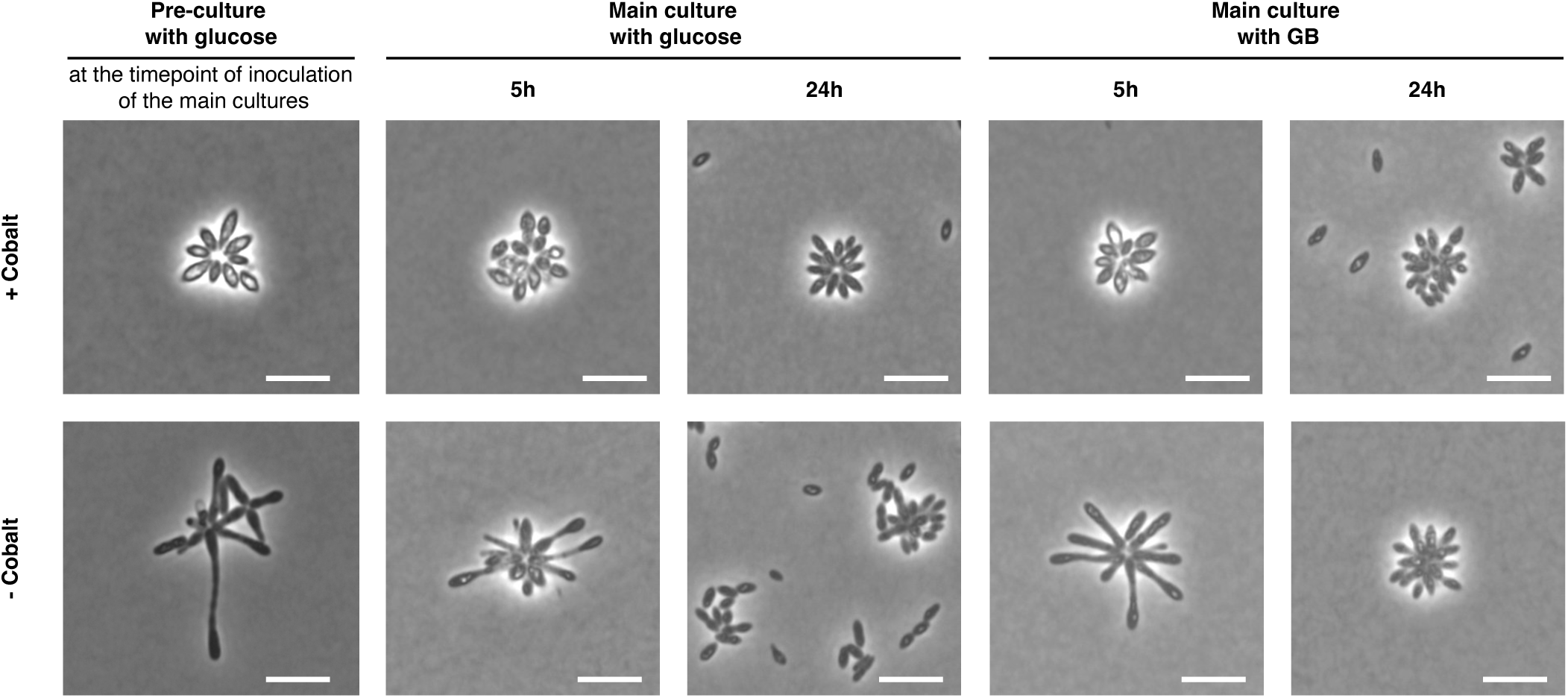
Transient elongation of *P. inhibens* cells in response to cobalt limitation. Phase contrast wide field microscopy of *P. inhibens* cells grown on 5.5 mM glucose or GB as sole carbon and energy source under cobalt-replete (+ Cobalt) and -limited conditions (- Cobalt). Pre-cultures were imaged after 48 h of cultivation at the timepoint of inoculation of the main cultures. Main cultures were imaged after 5 and 24 h of cultivation.

### *P. inhibens* supplies *G. huxleyi* with cobalamin during co-cultivation

During co-cultivation, *G. huxleyi* produces GB, which *P. inhibens* catabolizes^14,15^, thereby increasing its Cbl biosynthesis (Fig. 3b). However, the level of Cbl dependency of *G. huxleyi* is debated^26,31,32^. Thus, we tested whether *G. huxleyi* (strain CCMP3266) requires Cbl for its growth by passaging it in medium without Cbl supplementation. This led to significantly reduced growth, but not to a complete growth arrest (Fig. 5). However, co-cultivation of *G. huxleyi* with *P. inhibens* entirely reverted the growth deficiency of *G. huxleyi* under Cbl-limited conditions. This indicates that *P. inhibens* supplies *G. huxleyi* with Cbl, which increases algal cell numbers during co-cultivation.

**Figure 5:**
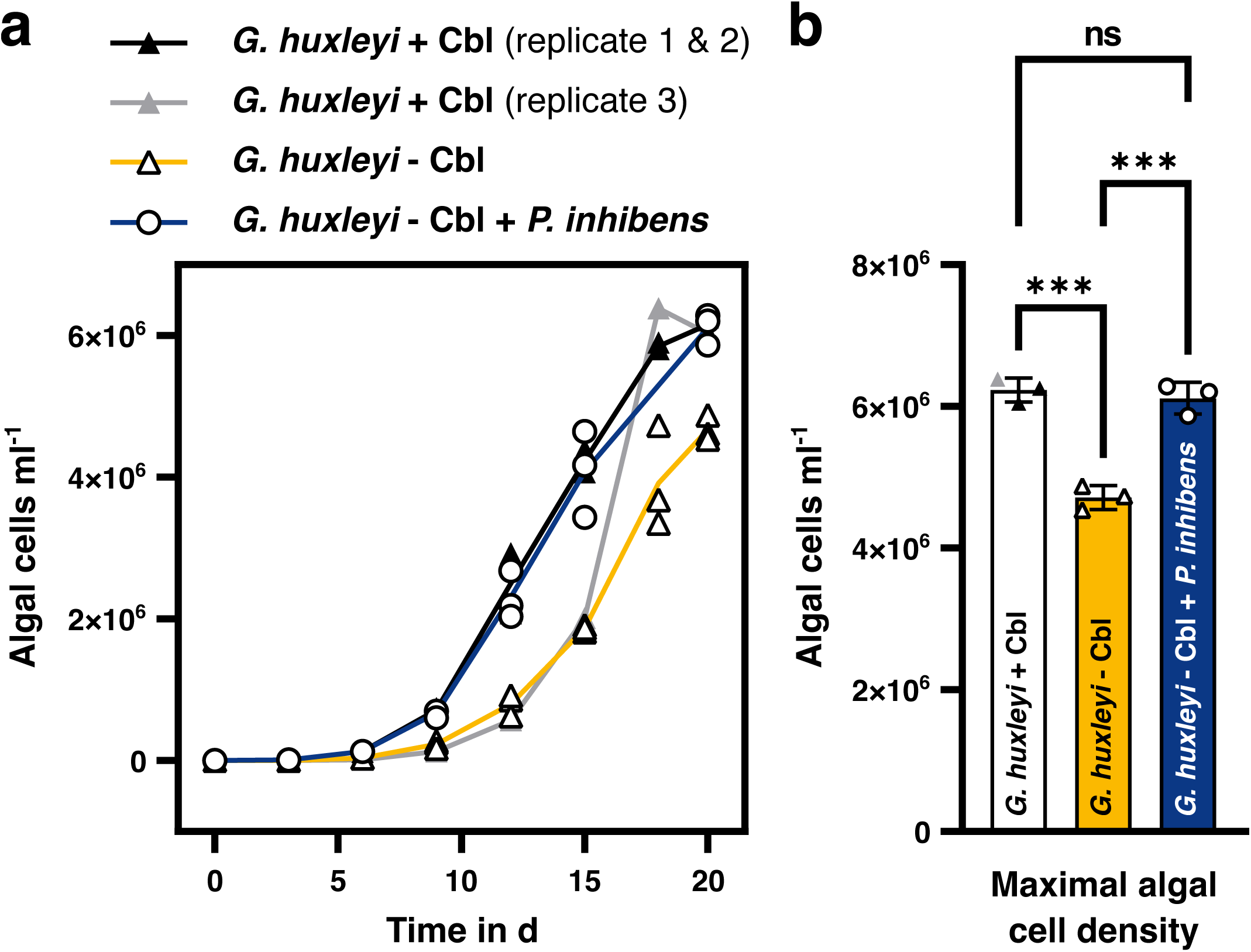
*P. inhibens* restores growth of cobalamin-limited *G. huxleyi*. *G. huxleyi* showed growth deficiency in axenic cultures without Cbl supplementation (- Cbl) in contrast to axenic cultures supplemented with Cbl (+ Cbl). Growth of *G. huxleyi* was restored in medium without Cbl supplementation by co-cultivation with *P. inhibens* (- Cbl + *P. inhibens*). **a,** Lines represent the mean algal cell numbers; triangles or dots represent the individual measured values. Cultivations were done in biological triplicates (n = 3). Replicate 3 of *G. huxleyi* + Cbl was plotted individually because of distinct growth but eventually reached a similar maximal cell density as replicates 1 and 2. **b,** The maximal algal cell density measured within an interval of 20 days of cultivation in the monocultures of *G. huxleyi* under Cbl-limited conditions is lower than under Cbl-replete conditions or in the co-culture with *P. inhibens* under Cbl-limited conditions. Bars represent the mean value; the individual measured values are represented by triangles or dots. The error bars represent the standard deviation (n = 3). To determine the significance of differences, ANOVA followed by Tukey’s post-hoc test was used (adjusted *P*-value > 0.05: ns, not significant; adjusted *P*-value ≤ 0.001: ***, significant).

### Cbl-dependent and -independent GB demethylation is widespread in Cbl-producing *Alphaproteobacteria*

To determine whether GB-stimulated Cbl production is a common mechanism underlying algal-bacterial cross-feeding, we evaluated presence-absence variations of *mtgBCDE* and *gbcAB* genes, the *gbcAB* 5’-UTR riboswitch and Cbl biosynthesis genes in 40 genomes from the order *Rhodobacterales* (Supplementary Tables S3, S4 and S5 as well as Supplementary Data S4). Importantly, only the *mtgE* gene, which links GB demethylation with methionine synthesis, was part of the *Rhodobacterales* core genome. Of the 40 analyzed *Rhodobacterales* members, 36 possess *mtgBCD*, 22 of which additionally possess *gbcAB*. The four genomes that lack *mtgBCD* contain *gbcAB* but lack the Cbl riboswitch. Since *mtgCD* encode proteins that are also part of a split methionine synthase, all 4 genomes lacking *mtgBCD* encode either the canonical methionine synthase MetH or a distinct split methionine synthase. The latter consists of a multi-domain protein (MetH2, pterin-binding, cobalamin-binding and reactivation domains), while the homocysteine *S*-methyltransferase is encoded by a separate gene (*metH1*). In 27 genomes – including the confirmed Cbl producer *P. inhibens* DSM 17395 – we found 28 genes encoding enzymes required for aerobic *de novo* Cbl biosynthesis (Supplementary Table S4). All 40 genomes contained at least 25 of the Cbl biosynthesis genes. This indicates that despite the common potential to biosynthesize Cbl, in many *Rhodobacterales* genomes, the genes encoding the Cbl-independent GB demethylase GbcAB are maintained in addition to *mtgBCD* encoding the Cbl-dependent GB demethylase.

To assess the broad applicability of our findings across phylogenetically diverse marine bacteria, we analyzed 15,525 dereplicated, high-quality genomes from the Ocean Microbiomics Database (OMDB)^84,85^. Using these species-level representative genomes, we investigated the prevalence of protein families (Pfams) associated with Cbl-dependent (MtgBCD) and -independent (GbcAB) GB demethylation, as well as Cbl biosynthesis (Supplementary Data S5). This analysis revealed that only about 18% of marine bacteria have the genetic potential to synthesize Cbl *de novo* (Fig. 6 and Supplementary Fig. S3). Among the *Alphaproteobacteria*, this trait is more widespread: 45% of *Alphaproteobacteria* possess the required genes. Other important Cbl producers are members of the *Desulfobacterota*, *Bacillota*, *Actinomycetota* and *Cyanobacteriota* (Fig. 6). Of the predicted Cbl producers, 58% possess at least one of the GB demethylases, while 26% possess both (Supplementary Fig. S3). Importantly, Cbl-dependent and -independent GB demethylation mainly co-occurs in alphaproteobacterial orders that represent 93.3% of the Cbl producers possessing both demethylases. Besides the *Rhodobacterales*, to which *P. inhibens* belongs, this also includes the globally abundant *Puniceispirillales* (SAR116) (Fig. 6)^86–89^. In contrast, only 31% of the non-producer possess at least one of the GB demethylases with prevalence of GbcAB (Supplementary Fig. S3). Our data shows that GB demethylation is a prevalent trait of Cbl-producing *Alphaproteobacteria*, half of which maintain both Cbl-dependent and - independent GB demethylases (Supplementary Fig. S3).

**Figure 6:**
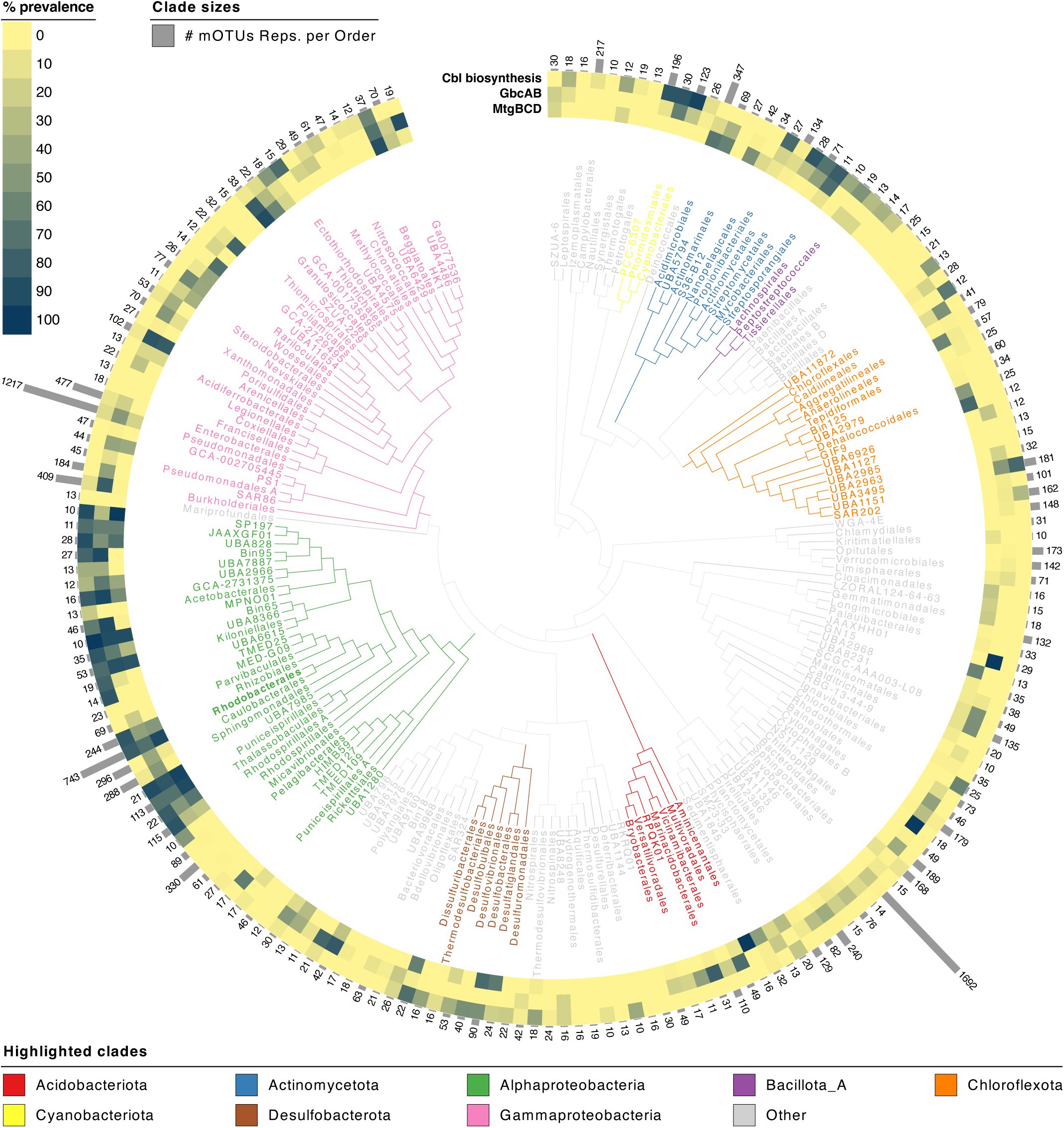
Phylogenetic distribution of genes encoding cobalamin-dependent and -independent GB demethylation together with cobalamin biosynthesis in marine bacteria. A database containing 15,525 dereplicated, high-quality, species-level genomes of marine bacteria (isolate genomes, metagenome assembled genomes and single-cell amplified genomes) was searched for the presence of protein families (Pfams) associated with Cbl-dependent and - independent GB demethylases. The two functions were considered as present if all Pfams of MtgBCD or GbcAB, respectively, were detected. Following Sañudo-Wilhelmy *et al.* (2014), we further queried 16 marker genes involved in Cbl biosynthesis and considered the pathway as present in a genome when > 75% of the corresponding Pfams were detected^47^. Pfam prevalences were mapped onto the GTDB taxonomic tree (R220), which was collapsed at the order level. Only orders represented by ≥ 10 genomes are displayed, with the outer bars indicating the number of genomes per order. The analyzed genomes are a subset of the OMDB and include only representatives of species-level operational taxonomic units (mOTUs).

## Discussion

Cross-feeding of fixed carbon and Cbl between microalgae and heterotrophic bacteria is a key microbial interaction in the ocean, where Cbl scarcity can limit algal growth and carbon fixation^25,27–30^. GB is a dominant marine metabolite and a chemical mediator of algal-bacterial interactions^6,14,15,90^. Our results highlight a stimulatory effect of GB on the Cbl production in *P. inhibens* (Fig. 3 b). Of the *N*- and *S*-methylated compounds tested in this study, none had a comparably stimulating effect on the Cbl production. This is in line with the reported specificity of the stimulatory effect of GB in *Pseudomonas denitrificans*^45^. A recently reported modular Cbl-dependent GB demethylase (MtgBCD) links GB catabolism and Cbl co-factor requirements in abundant marine bacteria^44^. While Mausz *et al.* could not identify genes encoding the Cbl-independent heterodimeric GB monooxygenase GbcAB in *P. inhibens*, we found that *P. inhibens* and many other abundant marine bacteria possess *gbcAB* in addition to *mtgBCD* (Fig. 6). In *P. inhibens*, the respective genes encoding both enzymes are co-expressed in the presence of GB (Fig. 1), and the availability of cobalt determines the enzyme used for GB demethylation (Fig. 3a). The switch between both enzymes is regulated by the intracellular concentration of Cbl that in turn is influenced by cobalt availability (Fig. 3b). We conclude that free Cbl binds to MtgC serving as a co-factor for enzymatic Cbl-dependent GB demethylation and to the Cbl riboswitch located in the 5’-UTR of the *gbcAB* mRNA, preventing its translation (Fig. 3c+d).

In the pelagic ocean, cobalt can be scarce with concentrations in the low picomolar range^27,91,92^ Cobalt acquisition is crucial for *P. inhibens* when growing on GB. This is indicated by the upregulation of the cobalt uptake-related gene *cbiM* which shows the highest expression of all differentially expressed genes. Post-transcriptional regulation of *gbcAB* and *cbiM* via Cbl riboswitches allows *P. inhibens* to rapidly respond to dynamic nutrient fluxes and to oppose cobalt depletion. Furthermore, a Cbl riboswitch sequence was found in the 5’-UTR of a gene encoding a DUF1636 domain-containing protein that is potentially involved in cobalt insertion into the corrin-ring^93^. Considering its position as first gene of a large Cbl biosynthesis operon (Supplementary Fig. S1), the riboswitch likely affects the translation of the entire transcript^50^. This is a common mechanism to regulate Cbl uptake or biosynthesis and usually prevents overproduction of Cbl^46,49,50^.

We propose that GB stimulates the Cbl production in *P. inhibens* not only by inducing upregulation of Cbl biosynthesis genes, but also by strongly inducing the expression of the Cbl-binding protein MtgC. If MtgC is not saturated, it can capture *de novo* biosynthesized Cbl, preventing binding of Cbl to the target riboswitches and thus negative feedback on the Cbl biosynthesis. In contrast, when *mtgC* is expressed at a moderate level, such as during growth on glucose as sole carbon source, *de novo* biosynthesized Cbl quickly exceeds the binding capacity of MtgC and free Cbl can bind to the riboswitch to prevent further biosynthesis. Similarly, MtgC could influence translation of the *gbcAB* mRNA. This is supported by the characteristic growth phases of the Δ*mtgB1 and* Δ*mtgC* mutants (Fig. 2 a). The Δ*mtgC* mutant shows a more severe growth defect under cobalt-replete conditions despite methionine supplementation and fails to establish stable growth. This might be due to a “stop-and-go” translation of *gbcAB* as decreasing Cbl concentrations recurrently allow translation and initiate Cbl biosynthesis which immediately backfires due to the lack of the buffering action of MtgC. In addition to its previously reported and here confirmed role in GB demethylation and methionine biosynthesis^44,57^, we suggest a potential regulatory function of MtgC by controlling the levels of free Cbl. This is further supported by the initially high expression of *mtgC* that is decoupled from *mtgBD* expression during co-cultivation of *P. inhibens* and *G. huxleyi* (Fig. 2).

The intertwining between methionine biosynthesis and GB demethylation^44^ is another factor that likely supports the GB-stimulated Cbl biosynthesis. Eight molecules of *S*-adenosylmethionine (SAM) are required per molecule of Cbl formed. Methionine biosynthesis via MtgCDE ensures SAM recycling via the methionine cycle while GB catabolism via MtgBCD and subsequent enzymes adds three methyl groups to the one-carbon pool eventually yielding glycine (Fig. 1a). The methyl groups can be used by MtgCDE or enter the one-carbon oxidation pathway. This way, activity of both enzyme complexes contributes directly to the supply of important resources (SAM and glycine) and energy for Cbl biosynthesis. The functional significance of *mtgE* downregulation in the presence of methylated substrates awaits elucidation. Increased spermidine biosynthesis during growth on GB is suggested by upregulation of *speDE* (Fig. 1) and may additionally enhance the activity of MtgBCD/MetgCDE as spermidine had a stimulatory effect on the activity of rat liver MetH (up to 400-fold increase) likely because it prevented oxidation of the catalytically active cobalt(I)^94^. Growth on GB strongly induced expression of *fixABC* genes, suggesting that electrons generated during dimethylglycine demethylation are transferred to ETFs and funneled into the electron transport chain^74,75^. Without this route, these electrons would reduce OLJ to toxic HLJOLJ. A functional link between GB catabolism and ETFs was recently proposed in *Vibrio natriegens* using guilt-by-association analysis^51^. Notably, in *P. inhibens* the *fixAB* genes lie immediately adjacent to *pduO*, which encodes the cob(I)alamin adenosyltransferase that accepts electrons from reduced ETFs^95^. This genomic arrangement raises the intriguing possibility that electrons liberated during sequential GB demethylation to glycine are directly channeled into late-stage Cbl biosynthesis. In summary, GB catabolism and Cbl biosynthesis are tightly intertwined in *P. inhibens*, which presumably drives Cbl overproduction in the presence of GB.

Our data suggest that allelopathic interactions between *G. huxleyi* and *P. inhibens* involve a positive feedback loop of GB-induced bacterial Cbl production and algal GB biosynthesis (Fig. 7). This model is strongly supported by the coordinated upregulation of GB demethylation, cobalt uptake and Cbl biosynthesis genes during the mutualistic growth phase of the co-culture (Fig. 1 and 2). *G. huxleyi* can produce high amounts of GB with intracellular concentrations reaching up to 30 to 35 mM^33–35^. SAM is the donor of three methyl groups for algal GB biosynthesis and recycled via the methionine cycle (Fig. 7). In algae like *G. huxleyi* that lack *metE*, this obligatorily involves the Cbl-dependent MetH. The required Cbl can be provided by *P. inhibens* that boosts its Cbl production in response to GB (Fig. 3b). The bacterial Cbl increases algal cell numbers (Fig. 5), which increases GB production and completes the feedback loop^15^. When cobalt is depleted, bacteria decrease their Cbl production (Fig. 3b). This could cause the breakdown of the positive feedback loop. Considering cobalt scarcity in ocean water, the here described cross-feeding dynamics have the potential to influence marine phytoplankton species composition and primary productivity^27,30,91,92^.

**Figure 7:**
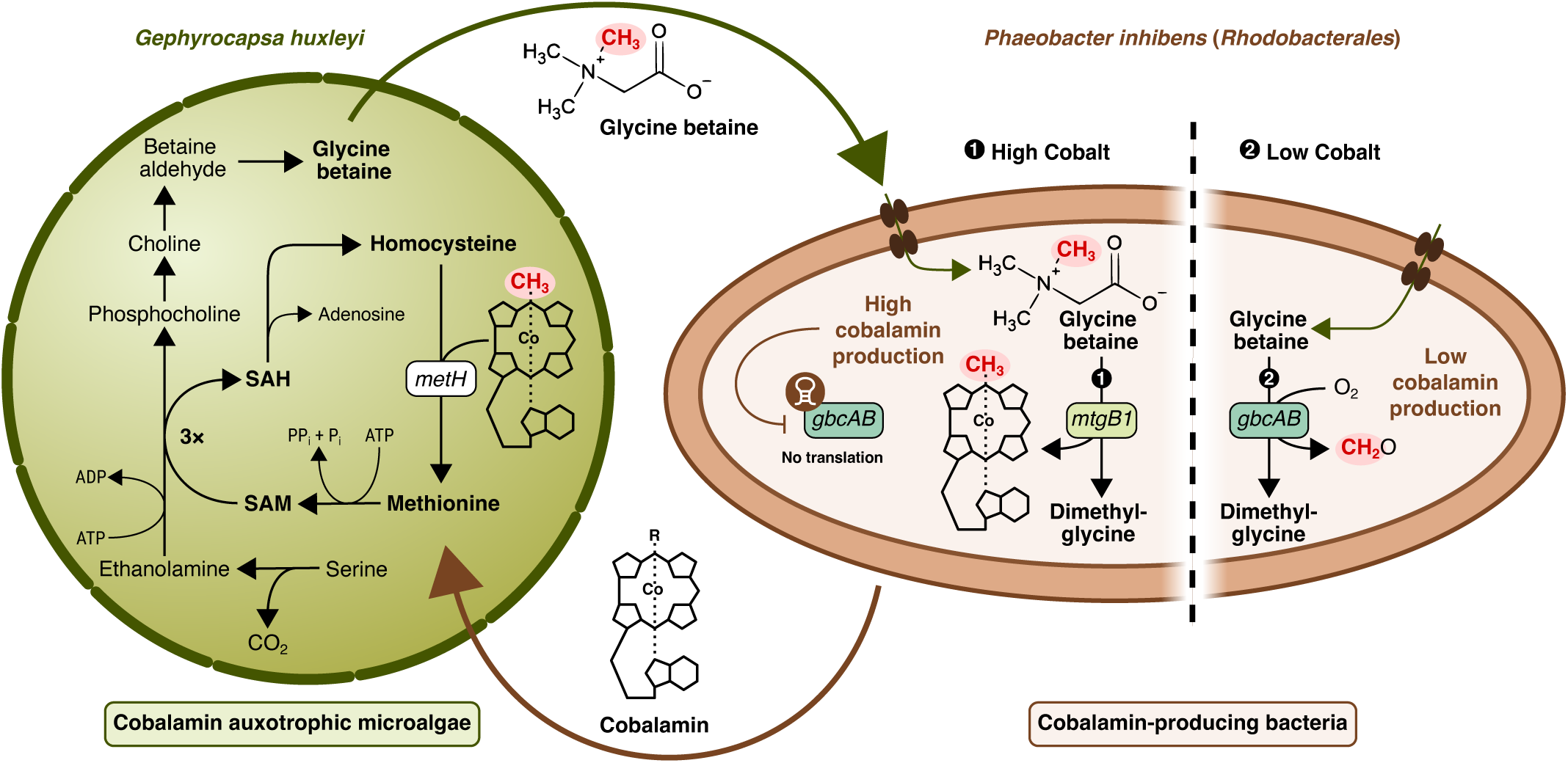
Proposed model of the positive feedback between algal GB biosynthesis in *G. huxleyi* and GB-stimulated Cbl production in *P. inhibens* under cobalt-replete and -limited conditions. During algal GB biosynthesis, recycling of the methyl group donor SAM involves the Cbl-dependent methionine synthase encoded by *metH*. GB is shared with *P. inhibens* and stimulates the bacterial Cbl-production and thus the amount of Cbl shared with the alga. In return this stimulates algal GB biosynthesis. *P. inhibens* demethylates algal GB using a Cbl-dependent or -independent demethylase, encoded by *mtgB1* or *gbcAB*, respectively, depending on the available amounts of cobalt that modulate Cbl biosynthesis.

The proposed GB-Cbl feedback loop might be a universal mechanism, as *P. inhibens* also provided Cbl in co-cultures with the diatom *Thalassiosira pseudonana*^24^ that is also known to produce GB^34^. However, sharing of Cbl does not occur systematically and phylogenetically close relatives show differences in their sharing behavior independent of their general capability of Cbl biosynthesis^24,96^. Strain-specific differences in the bacterial response to chemical mediators like GB as well as the varying amounts of GB shared by different algal species may contribute to this phenomenon. How Cbl is released by bacteria remains mostly elusive. In co-cultivation experiments of *Roseovarius* sp. (*Rhodobacterales*) and *Colwellia* sp. the activation of a prophage was required for successful Cbl cross-feeding, indicating a merely passive Cbl release^83^. We observed an upregulation of prophage genes during growth with GB though at low expression levels (TPM < 50). One possible explanation is that only a small subpopulation of *P. inhibens* cells is affected by prophage activation. This could be the consequence of GB-induced redox stress^15^ and a heterogenic stress response^97^. Lysis of a few bacterial cells with high intracellular concentrations of Cbl might be sufficient to cover algal demands. Considering the low number of bacterial reads in the RNA-Seq data for the co-cultures (between 0.2 to 0.5 Million reads)^14^ in comparison to the RNA-Seq data of this study (7.1 to 10.3 Million reads, Supplementary Table S6), pro-phage activation in a small subpopulation during algal-bacterial co-cultivation would likely not be detectable.

Remarkably, *P. inhibens* activates an expression profile typically associated with microaerophilic/anaerobic growth while growing exponentially on GB under aerobic shaking conditions or in co-cultures with photosynthetic, oxygen-producing *G. huxleyi* (Fig. 2 and Supplementary Table S2). In the facultatively anaerobic *Dinoroseobacter shibae* (*Rhodobacterales*), these gene expression changes are induced by the oxygen-sensitive transcriptional regulator FnrL during the transition from aerobic to anaerobic growth^66^. However, a disturbed redox balance can also lead to activation of Fnr transcriptional regulators as in a Δ*phaC* mutant of *Herbaspirillum seropedicae*, that is unable to synthesize polyhydroxybutyrate^98,99^. Thus, we suggest that increased electron fluxes into the electron transport chain may lead to oxygen depletion and induce changes in the redox status of *P. inhibens* cells during growth on GB. This may support the enzymatic Cbl-dependent demethylation reaction as it potentially lowers the frequency of cobalt autoxidation. However, under hypoxic conditions reductive stress can also lead to the formation of reactive oxygen species which are then detoxified to H_2_O_2_^100^. This fits to the recently reported increased H_2_O_2_ production of *P. inhibens* in response to GB^15^. Thus, increased Cbl production may also mitigate redox stress as in the acidophilic iron-oxidizing bacterium *Leptospirillum* Group II CF-1^101^. In eukaryotes it has been frequently reported that Cbl also plays a role in redox balancing and Cbl itself has superoxide dismutase activity^102–106^. Since the Cbl-indendendent monooxygenase GbcAB strictly requires O_2_ for GB demethylation, further research is necessary to investigate how exactly GB affects the redox status and electron transfer in *P. inhibens* and how this may change upon cobalt limitation. The observed transient cell elongation during adaptation to cobalt limitation could be a symptom of such changes in the redox balance and connected to lower Cbl levels (Fig. 4). Under conditions of increased redox stress, GbcAB might be more prone to inactivation than MtgBCD because of its redox-sensitive Rieske [2Fe-2S] iron-sulphur cluster. Consistent with this, Cbl-independent enzymes for deoxyribonucleotide and methionine biosynthesis are redox-sensitive, explaining why genes encoding their Cbl-dependent counterparts are upregulated under oxidative stress conditions in association with eukaryotic hosts^107–109^. As an example, the Cbl-dependent ribonucleotide reductase NrdJ is essential in *Sinorhizobium meliloti* to establish the root nodule symbiosis with its plant host^109^. Since *S. meliloti* also possess *mtgBCD* and *gbcAB* including the riboswitch (Supplementary Table S3), regulation of Cbl-related genes during bacteria-phototroph interactions might be a universal trait that initially evolved during marine algal-bacterial interactions.

The distribution of the genes *mtgBCD*, *mtgE* and *gbcAB* as well as the riboswitch in the 5’-UTR of *gbcAB* in the genomes of 40 marine and terrestrial members of the *Rhodobacterales* sheds light on the evolution and diversification of GB demethylation and methionine biosynthesis genes (Supplementary Table S3). In constantly cobalt-rich environments, the Cbl-dependent regulation of *gbcAB* can lead to its loss since the organism can always use MtgBCD. This has been shown for the MetH/MetE couple in the microalgae *Chlamydomonas reinhardtii* that loses the functionality of MetE when Cbl is always present due to constant downregulation of *metE*^110–112^. Similarly, cobalt-rich environments can lead to the loss of the riboswitch of *gbcAB* for the same reason. If an organism additionally acquired an alternative methionine synthase this can lead to the loss of *mtgBCD*. In environments with low oxidative stress or less GB this might not be a disadvantage and may explain why *mtgBCD* is less abundant in terrestrial members of the *Rhodobacterales* while it is dominant in phototroph-associated marine members of this order^113^. Beyond the *Rhodobacterales*, Cbl-dependent and -independent GB-demethylases co-occur in many Cbl-producing orders within the *Alphaproteobacteria* and emphasize the role of this class as main Cbl producers in the ocean (Fig. 5). Considering the previously reported stimulatory effect of GB on the Cbl biosynthesis in bacteria like *P. denitrificans* and *Priestia megaterium*^45^ – that lack the Cbl-dependent GB demethylase – a GB-stimulated Cbl production is likely more widespread than the Cbl-dependent GB demethylation in various phylogenetically distinct bacterial lineages. This emphasizes GB’s critical role as a currency in cross-feeding between Cbl producers and auxotrophs.

## Materials and Methods

### *Phaeobacter inhibens*, media composition and general cultivation conditions

The bacterial strain *P. inhibens* DSM 17395 was purchased from the German Collection of Microorganisms and Cell cultures (DSMZ, Braunschweig, Germany). *P. inhibens* was cultivated in complex medium, either in liquid ½ YTSS (yeast extract, 2□g □L^−1^; tryptone, 1.25□g□ L^−1^; Sigma sea salts, 20□g □L^−1^), or on solid ½ YTSS agar plates (agar, 16□g□ L^−1^), or in liquid marine M1H NAG ASW_v.1_ culture broth as described by Jeske. *et al.* (2016) but without vitamin solution and with reduced amounts of glucose^114^. Marine M1H NAG ASW_v.1_ culture broth contained basal artificial seawater version 1 (ASW_v.1_, 11.74 g L^-1^ NaCl, 1.96 g L^-1^ Na_2_SO_4_, 5.32 g L^-1^ MgCl_2_ · 6H_2_O, 715 mg L^-1^ CaCl_2_ · 2H_2_O, 96 mg L^-1^ NaHCO_3_, 346 mg L^-1^ KCl, 48 mg L^-1^ KBr, 13 mg L^-1^ H_3_BO_3_, 20 mg L^-1^ SrCl_2_ · 6H_2_O, and 1.5 mg L^-1^ NaF), 0.25 g L^-1^ peptone, 0.25 g L^-1^ yeast extract, Hutner’s mineral salts (250 µg L^-1^ Na-EDTA, 1.095 mg L^-1^ ZnSO_4_ 7 H_2_O, 500 µg L^-1^ FeSO4 · 7 H_2_O, 154 µg L^-1^ MnSO_4_ · H_2_O, 39.5 µg L^-1^ CuSO_4_ · 7H_2_O, 20.3 µg L^-1^ CoCl_2_ · 6H_2_O, 17.7 µg L^-1^ Na_2_B_4_O_7_ · 10H_2_O), 0.1% (w/v) N-acetylglucosamine, 0.0025% (w/v) glucose and trace elements (1.5 mg L^-1^ N(CH_2_COONa)_3_ · H_2_O, 500 µg L^-1^ MnSO_4_ · H_2_O, 100 µg L^-1^ FeSO_4_ · 7H_2_O, 100 µg L^-1^ Co(NO_3_)_2_ · 6H_2_O, 100 µg L^-1^ ZnCl_2_, 50 µg L^-1^ NiCl_2_ · 6H_2_O, 50 µg L^-1^ H_2_SeO_3_, 10 µg L^-1^ CuSO_4_ · 5H_2_O, 10 µg L^-1^ KAl(SO_4_)_2_ · 12H_2_O, 10 µg L^-1^ H_3_BO_3_, 10 µg L^-1^ NaMoO_4_ · 2H_2_O, 10 µg L^-1^ Na_2_WO_4_ · 2H_2_O) and was buffered with 10 mM HEPES at pH 8.0. For solid marine M1H NAG ASW_v.1_ culture broth agar plates 15 g L^-1^ agar was added.

*P. inhibens* was cultivated in a defined mineral medium based on a previously published protocol^115^, either in ASW_v.2a_ medium that contained basal artificial seawater version 2 (ASW_v.2_, 23.926 g L^-1^ NaCl, 4.008 g L^-1^ Na_2_SO_4_, 677 mg L^-1^ KCl, 98 mg L^-1^ KBr, 3 mg L^-1^ NaF, 21 mg L^-1^ Na_2_CO_3_, 168 mg L^-1^ NaHCO_3_, 10.3 g L^-1^ MgCl · 6H_2_O, 1.5 g L^-1^ CaCl_2_ · 2H_2_O, 25 mg L^-1^ SrCl_2_ · 6H_2_O), vitamins (thiamine HCl, 100 □μg □L^−1^; biotin, 0.5□ μg□ L^−1^), L1 trace element solution (4.36 □mg L^−1^ Na_2_EDTA · 2H_2_O, 3.15□ mg□ L^−1^ FeCl_3_ · 6H_2_O, 178.1□ μg L^−1^ MnCl_2_ · 4H_2_O; 23 □μg □L^−1^ ZnSO_4_ · 7H_2_O, 11.9□μg□ L^−1^ CoCl_2_ · 6H_2_O, 2.5 □μg □L^−1^ CuSO_4_ · 5H_2_O, 19.9□ μg □L^−1^ Na_2_MoO_4_ · 2H_2_O, 2.7□ μg □L^−1^ H_2_SeO_3_, 2.6□ μg□ L^−1^ NiSO_4_·6H_2_O, 1.8 □μg□ L^−1^ Na_3_VO_4_, 1.9□ μg □L^−1^ K_2_CrO_4_), L1 nutrients (75 mg L^-1^ NaNO_3_, 5 mg L^-1^ NaH_2_PO_4_ · H_2_O), and NSP solutions (5 mM NH_4_Cl, 33 mM Na_2_SO_4_, 2 mM KH_2_PO_4_) or in ASW_v.2b_ medium that is identical to ASW_v.2a_ except containing only 1.29 µg L^-1^ H_2_SeO_3_ and lacking KH_2_PO_4_, or in ASW_v.2c_ that is identical to ASW_v.2a_ except lacking Na_2_SO_4_, KH_2_PO_4_ and vitamins. The pH was adjusted to 8.0 with HCl. Defined mineral or complex media were used as indicated. In defined mineral medium carbon sources were added as indicated. Where indicated, the medium was prepared with a trace element solution without CoCl_2_ · 6H_2_O (- Cobalt). If not indicated otherwise, the trace element solution with cobalt (+ Cobalt) was used. For long-term storage ½ YTSS with 15% glycerol or marine M1H NAG ASW_v.1_ culture broth with 50% glycerol stock medium (12.6 g L^-1^ K_2_HPO_4_, 900 mg L^-1^ Na-citrate, 100 mg L^-1^ MgSO_4_ · 7H_2_O, 1.8 g L^-1^ (NH_4_)_2_SO_4_, 3.6 g L^-1^ KH_2_PO_4_, 88 g L^-1^ glycerol) were used and cryo stocks were stored at -80°C.

### Cultivation, RNA extraction, sequencing and analysis

Bacterial pre-cultures were prepared in 30 mL ASW_v.2b_ medium with 5.5 mM glucose or GB. For inoculation, a single *P. inhibens* colony was picked from a ½ YTSS agar plate cultivated for 48 h at 30°C, resuspended in ASW_v.2b_ and used to inoculate two Erlenmeyer flasks containing either glucose or GB. Pre-cultures were incubated for 24 h at 30°C. Main cultures were prepared in 100 mL ASW_v.2b_ medium with 5.5 mM glucose or GB, using 4 replicate flasks per carbon source, and inoculated with the respective pre-culture to an OD_600_ of 0.01 (Ultrospec 2100 Pro, Biochrom). Growth of the eight main culture flasks was monitored hourly by measurement of OD_600_. When the cells reached an OD_600_ of 0.10 ± 0.02, while still being within the exponential growth phase (Supplementary Fig. S4), 50 mL culture were harvested by 5 min centrifugation at 4,000 rpm and room temperature. The supernatants were removed by vacuum. Cells were immediately resuspended in 450 µL RLY lysis buffer (from ISOLATE II RNA Mini Kit, Meridian Bioscience) + 1% β-mercaptoethanol by vortexing to quench RNase activity. The suspensions were transferred into 2 mL screw cap tubes with 300 mg beads (SPEX Low Binding Silica Beads, 100 µm). The tubes were put into liquid nitrogen for snap freezing and stored overnight at -80°C until RNA extraction. The OD_600_ of the main cultures was checked again approximately 1 h after cell harvest to ensure cell harvest during exponential growth.

RNA extraction and library preparation was performed as previously reported for *P. inhibens*^14^. After disrupting cells by 5□min bead beating at 30□s^−1^ in a mixer mill MM 400 (Retsch), RNA was extracted using the ISOLATE II RNA Mini Kit (Meridian Bioscience), including the optional on-column DNase treatment. A second DNase treatment using the TURBO DNase kit (Thermo Fisher Scientific) was conducted after RNA elution. Then, RNA was purified with 2x concentrated RNAClean XP magnetic beads (Beckman Coulter), and RNA quantity and integrity were evaluated with the Qubit RNA HS Assay (Invitrogen, Thermo Fisher Scientific) and Tapestation RNA ScreenTape analysis (Agilent Technologies), respectively (Supplementary Fig. S5). Three of the four replicate RNA extracts per carbon source were selected for library preparation. For each of the extracts, 100 ng RNA were fragmented by heat (3 min, 94°C), followed by a DNase and FastAP treatment and another clean-up with 2x concentrated RNAClean XP magnetic beads. The RNA was 3’-tagged by ligating barcoded adapters 13 to 18 (RNase-free high-performance liquid chromatography purified; ordered from Integrated DNA Technologies)^116^. Barcoded RNA of all samples was pooled and purified with RNA Clean & Concentrator-5 columns (Zymo Research). The mRNA was enriched using the riboPOOL rRNA depletion kit in combination with Pan-Bacteria probes (siTOOLs Biotech). Afterwards, the enriched mRNA was purified with 2x concentrated RNAClean XP magnetic beads. Reverse transcription was conducted with the SuperScript IV First-Strand Synthesis System (Invitrogen, Thermo Fisher Scientific), using the AR2 primers^116^. The resulting cDNA was purified with 2x concentrated RNAClean XP magnetic beads and 3’-ligated to the 3Tr3 adaptor^116^ (HPLC purified; Integrated DNA Technologies). The resulting cDNA was purified twice with 2x or 1.5x concentrated RNAClean XP magnetic beads, respectively. After PCR cycle optimization, the 3’-ligated cDNA was amplified with 9 PCR cycles, which added Illumina sequencing adapters. The amplified cDNA was purified with a double-sided clean-up step (right: 0.5x, left: 1.5x; RNAClean XP beads), followed by a left-sided clean-up step (0.7x). Size distribution of the final library was evaluated via TapeStation ScreenTape analysis. The final library was sequenced on a NovaSeq 6000 System with a S1 100-cycle cassette (Illumina) in paired-end mode (read 1: 64□bp, read 2: 54□bp, index 1: 0□bp, index 2: 0□bp), aiming for 10 million reads per sample. RNA-seq data have been deposited in the NCBI Sequence Read Archive (SRA) and are publicly available under accession PRJNA1372585.

Data analysis was performed as previously reported for *P. inhibens*^14^. Briefly, Illumina raw reads were demultiplexed with fastq-multx (v1.3.1)^117^ (allowing one mismatch per barcode), quality filtered and trimmed with cutadapt (v1.18)^118^, mapped to the reference genome of *P. inhibens* DSM 17395 (accession: <u>GCF_000154765.2</u>) with STAR (v2.7.5c^119^, and reads per genes counted with featureCounts (v2.0.0). For each sample, sequencing depth, total gene feature counts and non-rRNA gene feature counts are summarized in Supplementary Table S6. After analyzing sample-to-sample correlations, we excluded the third replicate of the glucose condition, as sequencing failed because of wrong barcode assignment (Supplementary Fig. S6). Using DESeq2 (v1.36.0), differentially expressed genes were identified comparing the GB condition with glucose; the corresponding adjusted *P*-value and log2(fold changes) are provided in Supplementary Data S1. To allow comparison of expression levels between samples, gene feature counts were normalized to transcripts per million (TPM; Supplementary Data S1). Automatically generated NCBI RefSeq annotations^120^, partially curated GenBank submitter annotations^121^ and KEGG annotations^122^, as well as links to the unifying UniProt Archive (UniParc^123^ links redirecting to additional databases) were added to the feature table (Supplementary Data S1). Features of *P. inhibens* were filtered for relevance, based on differential gene expression (adjusted *P*-value < 0.05; |og_2_(fold change) > 0.585, upregulated; |og_2_(fold change) < 0.585, downregulated). The upregulated genes were then ranked either by the mean TPM (upregulated and highly expressed genes) or by the log_2_-(fold change) (highly upregulated genes). The top 100 genes of both ranked lists were analyzed to determine relevant pathways during growth on GB (Supplementary Data S1). For some genes that were noticeable e.g., due to a high expression levels or genetic context, domain annotations of the InterPro database^124^ were additionally used to assign putative functions. This information was used to create a metabolic map of GB metabolism and connected assimilatory and dissimilatory pathways in *P. inhibens* based on KEGG metabolic pathway annotation map^122^. Former gaps in the pathways were closed by consulting literature on GB demethylation^39,53–56^, methionine biosynthesis^57^ and one-carbon metabolism^125^ and searching for homologous genes in *P. inhibens* using nucleotide and protein BLAST^126^.

To investigate how the modularity of MtgBCD and MtgCDE is reflected by the expression levels of the coding genes in *P. inhibens* under various conditions, the transcriptional ratios were calculated based on mean TPM of *mtgE*, the sum of the mean TPM of *mtgB1-4* (encoding the GB--Cbl methyltransferase MtgB1 and related uncharacterized methyltransferases) and the mean TPM of *mtgC* and compared between the RNA-Seq data of this study and a previous publication, generated and processed with the same protocol and bioinformatics pipeline^14^. This included data derived from *P. inhibens* monocultures during lag-phase (either 20 or 40 min after inoculation) with 1 mM glucose with or without addition of 50 µM DMSP, during exponential growth with 2 mM or 5.5 mM glucose or 5.5 mM or during stationary phase with 2 mM glucose as well as from different time points of co-cultures with *G. huxleyi* (days 4, 6, 9 and 11/12). In the same way transcriptional ratios of *mtgB1*, *mtgC* and *mtgD*, *gbcA* and *gbcB* were calculated and compared between *P. inhibens* monocultures with GB (5.5 mM) and the days 4, 6 and 9 of the co-culture with *G. huxleyi*.

Identified differential gene expression patterns of *P. inhibens* during growth on GB were compared with the previously published dual RNA sequencing data for co-cultures of *P. inhibens* and *G. huxleyi*^14^, which were systemically processed with the same bioinformatics pipeline. In this and the previous study^14^, *P. inhibens* cultures grown with glucose as the sole carbon and energy source served as control condition for differential gene expression analysis. Additionally, the dual RNA-seq data of the previous study^14^ was reanalyzed to identify temporal changes in the bacterial expression levels during co-cultivation (before algal death) using DESeq2 to identify differentially expressed genes by comparing co-culture samples from day 4 and day 9 (Supplementary Data S2). To investigate whether bacterial expression patterns change after bacteria induced algal cell death of the algae, we used DESeq2 to identify differentially expressed genes by comparing co-culture samples before (day 4, 6 and 9) and after algal population collapse (day 11/12) (Supplementary Data S3).

### Mutant construction and molecular work

The bacterial strains and plasmids used in this study as well as their relevant characteristics are listed in Supplementary Tables S7 and S8. Oligonucleotides used in this study are listed in Supplementary Table S9. *Escherichia coli* K-12 strains were used for cloning and routinely cultivated in LB medium (5 g L^-1^ yeast extract, 10 g L^-1^ tryptone, 5 g L^-1^ NaCl) at 37 °C. If required, 50 mg L^-1^ kanamycin or 20 mg L^-1^ gentamycin were added for selection of recombinant clones and maintenance of plasmids. For the selection of recombinant *P. inhibens* cells, 30 µg mL^-1^ gentamycin or 50 µg mL^-1^ kanamycin was used. To generate the *mtgB1* knockout mutant (Δ*mtgB1*), ∼1,000□bp regions upstream and downstream of the *mtgB1* gene (accession: PGA1_RS07270) were PCR-amplified. The gentamycin resistance marker was amplified from the plasmid pBBR1MCS-5^127^. The PCR products (upstream region + gentamycin resistance + downstream region) were assembled and cloned into the pCII-TOPO vector (Invitrogen, Thermo Fisher Scientific) using restriction-free cloning^128^, generating the plasmid pDN5. DNA manipulation and cloning PCR were performed with Phusion High Fidelity DNA polymerase (Thermo Fisher Scientific), according to the manufacturer recommended PCR conditions. PCR-amplified DNA was purified using the NucleoSpin Gel and PCR Clean-up kit (Macherey-Nagel). For transformation, 10 □µg of the constructed plasmid were added to 300 □µL of competent *P. inhibens* DSM 17395 cells (prepared as in Dower *et al*., 1988)^129^ and electroporated with a 2.5□kV pulse (MicroPulser, Bio-Rad Laboratories). Electroporated cells were recovered in 2□mL of ½ YTSS medium for 12□h at 30□°C and then transferred to selective ½ YTSS agar plates containing gentamycin. The other knockout mutants were constructed with a related protocol with the following modifications: Q5 polymerase Master mix (New England Biolabs) was used for the amplification of ca. 1500 bp flanking homology arms of the inactivation target, cloning was performed by restriction-ligation, competent cells were prepared from 50 mL of an overnight culture in modified ½ YTSS with ASW_v.1_ instead of sea salts and a kanamycin resistance gene was used as additional selection marker to allow the construction of the double deletion mutant. Obtained deletion mutants were checked by PCR using primers that bind in the genome outside of the previously amplified homology regions. An additional PCR with primers binding within the open reading frame of the deleted gene was included to check for contamination with residual wild type cells and only PCR-negative strains were used for subsequent experiments. Successful null mutants were further verified by Sanger sequencing.

### Growth assays of *P. inhibens*

For the first pre-culture, ASW_v.2c_ medium with 5.5 mM glucose was inoculated with 6-9 colonies from Marine M1H NAG ASW_v.1_ culture broth agar plates that were incubated for 7 days. For each biological replicate, a separate pre-culture was prepared. The appropriate antibiotics (50 µg mL^-1^ kanamycin or 20 µg mL^-1^ gentamycin or both) were added for the *P inhibens* mutants (Supplementary Table S7). For the Δ*mtgC* mutant, 1 mM methionine was added to all pre-cultures. The pre-cultures were incubated for 72 h at 28°C with shaking (100 rpm). For the second pre-cultures ASW_v.2c_ medium was prepared with a trace element solution that did not contain CoCl_2_ · 6H_2_O. 5.5 mM glucose were added, and the second pre-culture was inoculated from the first pre-culture to an OD_600_ of 0.01 and incubated for 48 h under the same conditions as the first pre-culture. For the microplate reader assay, ASW_v.2c_ medium was prepared in two separate batches, with a trace elements solution that contained CoCl_2_ · 6H_2_O (ASW_v.2c_ medium with cobalt) and with a trace elements solution that did not contain CoCl_2_ · 6H_2_O (ASW_v.2c_ medium without cobalt). ASW_v.2c_ medium with cobalt and ASW_v.2c_ medium without cobalt was split in three batches each. To one batch no carbon source was added, to the second batch 1 mM glucose was added and to the third batch 1.2 mM GB was added (equimolar carbon). For the Δ*mtgC* mutant and a wild type control, 0.1 mM methionine were additional added to all three batches. The batches were distributed to 1.5 ml reaction vials for blanks and inoculation. Before inoculation, cells from the second pre-culture were harvested by centrifugation with 16,100 × g for 5 min at RT and washed in basal ASW_v.2_ three times before inoculation to a start OD_600_ of 0.01 (cuvette, 1 cm pathlength). Blanks and cultures were transferred to a 96 well plate (BRANDplates pureGrade S, Brand). The plate was sealed with parafilm, incubated at 28°C and the OD_600_ was monitored using a BioTek Epoch2 microplate spectrophotometer (Agilent). Each measurement cycle included two shaking phases while the shaking regime changed between linear to orbital to double orbital every 5 min. The shaking phases were interrupted by the OD_600_ measurements. To prevent condensation on the plate lid, a 2°C temperature gradient was established between the bottom and the lid of the plate. All cultivations were performed in biological triplicates. The mean OD_600_ values of each timepoint resulting from plate reader measurements for each biological triplicate and condition were blank corrected by subtracting the mean of the media blanks at each timepoint.

### Analysis of trace element concentrations via inductively coupled plasma - optical emission spectrometry

To evaluate concentrations of all trace elements (Co, Cr, Cu, Fe, Mn, Mo, Ni, Se, V and Zn) in the trace elements solutions with or without CoCl_2_ · 6H_2_O, samples of both solutions were acidified with 1.5% HNO_3_ and incubated for five days at 5°C to check for precipitation. Afterwards the samples were filtered with sterile Minisart syringe filters with a polyether sulfone membrane with a pore size of 0.22 µm (Sartorius). Inductively coupled plasma - optical emission spectrometry (ICP-OES) was performed with the simultaneous radial ICP-OES spectrometer 725ES (Agilent) with a charge-coupled device as detector. For calibration multi-element standard solutions were used. All samples were measured in triplicates. To calculate detection limits the three sigma criterium under consideration of the dilution factor was applied.

### Cobalamin extraction and analysis

To obtain cell mass for Cbl extraction, 1 L ASW_v.2a_ medium with 5.5 mM glucose or 5.5 mM betaine was inoculated from pre-cultures in ASW_v.2a_ medium with 5.5 mM glucose to an OD_600_ of 0.01 and incubated at 28°C with shaking (85 rpm). To screen the influence of various *N*- and *S*-methylated compounds on the Cbl production 1 mM of either DMSP, GB, carnitine or trigonelline were additionally added to a culture containing 5.5 mM glucose (Supplementary Table S10). DMSP, carnitine and trigonelline led to a decrease in the pH of the medium that was corrected by the addition of equimolar amounts of NaOH. All cultures were harvested at 1/3 to 2/3 of the maximal OD_600_ (OD_max_), except cultures with 5.5 mM GB that were harvested at OD_max_ to obtain sufficient biomass. For cell harvest the main culture underwent centrifugation at 11,900 × g for 10 min at 10°C. Because of their loose pellets, GB-grown cultures were centrifuged at 18,590 × g for 20 min. Supernatants were discarded, and cell pellets were kept on ice during further processing. The pellets were resuspended in 50 mM Tris-HCl buffer (pH 7.5) and collected in a fresh falcon tube. The weight of the falcon tube was determined in advance. The falcon tubes containing the resuspended cells were centrifuged again at 11,900 × g for 10 min or at 18,590 × g for 20 min, respectively, at 10°C. The supernatant was discarded, and the wet cell weight was determined. The cell pellet was stored at -20°C until Cbl extraction. All cultivations with GB or glucose as sole carbon and energy source were performed in biological triplicates while the cultures with glucose and an additional osmolyte were performed in biological duplicates.

For the analysis of cobalt limitation in cultures with 5.5 mM GB, cultivation and cell harvesting were conducted as described with following modifications: ASW_v.2a_ medium was prepared with trace element solution without CoCl_2_ and was used for the pre-culture as well as the main culture.

The Cbl extraction was performed according to a protocol modified after Stupperich *et al.* (1986)^130^. Cell pellets were resuspended in 50 mM Tris-HCl buffer (pH 7.5). The pH of the cell suspension was adjusted below pH 5 with glacial acetic acid. Potassium cyanide was added to each sample to a final concentration of 0.1 M. For Cbl extraction, the samples were boiled in a water bath for 15 min under stirring and then centrifuged at 17,420 × g for 10 min at 10°C. The supernatants (extracts) were collected in a fresh tube while the pellets were resuspended in 10 mL ultrapure water and underwent a second boiling step. Per mL Cbl extract 0.25 g XAD4 (Sigma-Aldrich) were resuspended in 100% HPLC-grade methanol and equilibrated with 500 mL of 0.1% acetic acid using a filter flask. The XAD4 was added to the Cbl extract, and the suspension was incubated overnight with gentle shaking (80 rpm). After XAD4 settled, the supernatant was discarded and the XAD4 was washed ten times with 10 mL ultrapure water. To elute Cbl from the XAD4-adsorber material, 10 mL 100% methanol were added, and the samples were incubated for 1 h with gentle shaking (80 rpm). The eluate was collected in a fresh tube and the elution was repeated twice as described before. The collected eluates were reduced to about 3 mL with a rotary evaporator and then completely dried overnight with a vacuum concentrator. Afterwards, the dried samples were resuspended in 2 mL ultrapure water and purified using a dry aluminum oxide column (3 g column material). The elution was performed with 40 mL of ultrapure water. The eluate was reduced to 2 mL using a rotary evaporator and completely dried overnight with a vacuum concentrator. The dried extracts were resuspended in 100 μL ultrapure water. The Cbl extracts were centrifuged at 16,100 × g for 20 min to remove insoluble particles and stored at 5°C until HPLC analysis.

To prepare Cbl standard solutions with a reliable concentration, the Cbl concentration (c) of the stock solution was determined by measuring the absorbance at 360 nm (A) of three independent dilutions (each measured as technical triplicates) and using the Lambert-Beer law 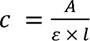 with l = 1 cm and E = 28.06 mM^−^1 cm^−1 131^. For the quantification of Cbl in the extracts via via HPLC (Knauer), Cbl standards were prepared in two independent serial dilutions (100, 50, 25, 10, 5, 2.5, 1 and 0.5 μM) to calculate Cbl concentrations from the peak areas of Cbl peaks (Supplementary Fig. S7). For HPLC analysis of the Cbl standards as well as the Cbl extracts a Kinetex 5μm EVO C18 100 Å column of the size 250 x 4.6 mm (Phenomenex) was used. For each sample the HPLC was run at a column temperature of 30°C with solvent A (14% (v/v) methanol, 0.2% (v/v) acetic acid) for 140 min with a flow of 0.5 mL min^-1^. The column was rinsed with solvent B (99.8% (v/v) methanol, 0.2% (v/v) acetic acid) and re-equilibrated with solvent A before a new sample was analyzed. The UV-Vis detector measured the absorbance at 360 nm (absorbance maximum of Cbl) and recorded the absorbance spectrum between 200 and 600 nm that is also characteristic for Cbl (Supplementary Fig. S8). The flow-through was collected in 1 ml fractions. Cbl peaks were identified by their retention time that was verified by mass spectrometry in a previous study^129^.

### Light microscopy of *P. inhibens*

To test the effect of cobalt limitation on the cell morphology of *P. inhibens*, ASW_v.2c_ medium with cobalt and supplemented with 5.5 mM glucose was inoculated with a single colony from marine M1H NAG ASW_v.1_ culture broth agar plates. The first pre-culture was incubated for 72 h at 28°C with shaking at 100 rpm. For the second passage, three cultures each were prepared in ASW_v.2c_ medium with cobalt or without cobalt containing 5.5 mM glucose. The second pre-cultures were inoculated from the first pre-culture to an initial OD_600_ of 0.01. After 48 h, main cultures were inoculated from the respective second pre-culture to an initial OD_600_ of 0.01 in ASW v2 medium with or without cobalt and supplemented with either 5.5 mM glucose or 5.5 mM GB. The cultures were incubated at 28°C with shaking at 100 rpm. Pre-cultures were sampled at the timepoint of inoculation of the main culture, and main cultures were sampled after 5 and 24 h of cultivation and prepared for microscopy. 2 µL of culture were spotted onto an 1% (w/v) agarose cushion covered with a cover slip and sealed with VLAP (33% vaseline, 33% lanoline, 33% paraffin, w/w) to limit sample movement and evaporation. Images were taken of all sample conditions and replicates.

For imaging, a microscope set-up was used as previously described^132^. Briefly, a Nikon Eclipse Ti2 inverted microscope that was used equipped with a Nikon Plan Apo λ objective (100x NA 1.45 Oil, ∞/0.17 WD 0.13) and a Hamamatsu Orca-Flash 4.0 LT Plus controlled with the Nikon© NIS Elements Software (Version 5.30.02). Native files were handled in FIJI^133^ which was also used for the adjustment of brightness and contrast.

### Co-culture of *P. inhibens* and *G. huxleyi* algae without cobalamin

Axenic cultures of *G. huxleyi* CCMP3266 algae, purchased from the National Center for Marine Algae and Microbiota (Bigelow Laboratory for Ocean Sciences), were maintained in Cbl-containing artificial seawater medium, as previously described^14^. Briefly, algae were kept at 18 °C under a 16/8h light/dark regime with 130 μmol photons m^−2^ s^−1^, and core cultures were routinely controlled for their axenic status. The algal culture medium resembles ASW_v.2a_, but without NPS solutions (named ASW _v.2d_), while Cbl was present (0.5□μg□L^−1^ cyanocobalamin). To test whether *P. inhibens* can provide Cbl to the algae, the axenic *G. huxleyi* core culture was transferred for two passages in 30 mL of Cbl-free ASW_v.2d_ medium, initiating each passage with a 1:10,000 inoculum of the preceding culture. A third passage was initiated with 600 algal cells/ml, splitting the culture into three new cultures with different conditions: 1) Cbl-free ASW_v.2d_, 2) Cbl-containing ASW_v.2d_ (0.5□μg□l^−1^ cyanocobalamin) and 3) Cbl-free ASW_v.2d_ with *P. inhibens*. For the latter condition, a 20% glycerol-containing cryostock of *P. inhibens* was revived on ½ YTSS agar plates (48 h at 30°C), pre-cultured in 30 mL of Cbl-free liquid ASW_v.2b_ medium with 2 mM glucose (48 h, 30°C), washed thrice by centrifugation, and diluted to attain 60 bacterial CFU mL^-1^ in the algal culture. Each of the three conditions was setup with three replicate flasks. Algal growth was monitored by enumerating cells using a CellStream flow cytometer with 561/702 nm excitation/emission (Merck).

### Pangenomic analysis of *Rhodobacterales*

To analyze the distribution of genes involved in Cbl-dependent and -independent GB demethylation, Cbl biosynthesis and methionine biosynthesis among members of the *Rhodobacterales*, 40 genomes representing ten different genera were selected to compute a pangenome of (Supplementary Table S11 and Supplementary Data S4). For the selection of genomes, the NCBI Reference Sequence database was filtered for complete genomes from representatives of the *Rhodobacterales* that are present in the metagenomes of two *G. huxleyi* blooms sampled in Gulf of Maine during the summer of 2015^7^. For some genera many complete genomes were available. Thus, additional criteria for genome selection were considered (listed in order of priority): the identity on species level to the reported species^7^, a marine origin of the isolate and the isolation source (e.g., algae favored over clams or environmental samples favored over industrial samples). Furthermore, well-characterized isolates were preferred to uncharacterized ones.

The genomes were processed and analyzed using the Anvi’o workflow for pangenomics as reported by Delmont *et al.* (2018) with Anvi’o version 7^134,135^. Anvi’o uses the gene prediction algorithm Prodigal to predict open-reading-frames^136^. The NCBI database of Clusters of Orthologous Groups (COG)^137^, the database of protein families (Pfam)^138^ as well as the database of KEGG Orthologs (KOfam)^122^ were used for the functional annotations of the genomes. The pangenome was computed with the program “anvi pan genome” using NCBI blast for sequence comparison between all the gene calls in all the genomes, a minimum minbit value of 0.5 to eliminate weak matches between two sequences and a Markov Cluster Algorithm inflation parameter of 8 determining the sensitivity of clustering of homologous genes. The computed pangenome was screened for genes related to GB demethylation, methionine biosynthesis, Cbl biosynthesis and cobalt uptake (Supplementary Table S5). Presence or absence of riboswitches in the 5’-UTR of *gbcAB* was analyzed by manually checking the RefSeq annotated genomes in the NCBI database^120^ that include annotations of RNA secondary structures generated using infernal (v1.1.5)^139^. To identify proteins associated with *mtgC*, the genomes were assigned to two groups based on presence or absence of *mtgC* (present: 36 genomes, absent: 4 genome). These groups were used for functional enrichment analysis based on the COG-, Pfam- or KOfam annotations by running the program “anvi-compute-functional-enrichment” which computes enrichment scores using an R script reported elsewhere^140^.

### Mapping of GB catabolism and Cbl biosynthesis across *Bacteria*

To analyze the occurrence of Cbl biosynthesis, Cbl-dependent GB demethylation, and Cbl-independent GB oxidation across the bacterial tree of life, we screened the mOTUs-db (https://motus-db.org/)^141^, which currently contains 3.75 million systematically processed prokaryotic genomes (MAGs, SAGs, and isolate genomes) from 118 thousand global samples, with genomes clustered at 124 thousand species-level operational taxonomic units (mOTUs). Each genome was taxonomically classified with GTDB R220 using GTDB-Tk (v2.4)^142^. The tool PyHMMER (-cut_ga; v0.10.15)^143^ was used to search the mOTUs-db for genomes with annotated protein families (Pfam v37.1) involved in Cbl biosynthesis (PF00590, PF01891, PF02654, PF01890, PF03186, PF02514, PF07685, PF07685, PF01888, PF01890, PF02570, PF02283, PF02571, PF01923, PF00590, PF00590), Cbl-dependent GB demethylation (MtgB: PF06253; MtgC: PF02310; MtgD: PF00809) and Cbl-independent GB demethylation (GbcA: PF00848; GbcB: PF00175) (Supplementary Table S12). The functions were flagged as present in a genome, if either 75 % of the Cbl biosynthesis Pfams, or all the Pfams associated with Cbl-dependent GB demethylation or Cbl-independent GB oxidation were present. We further filtered the mOTUs-db to retain only genomes of high-quality (≥ 90% completeness, ≤ 5% contamination). We also removed any archaeal genomes as well as genomes from other than marine sources and selected only the representative genomes per species-level mOTUs, reducing the set to 15,525 non-redundant, global genomes of marine bacteria (Data X) that are a subset of the Ocean Microbiomics Database (OMDB)^84,85^. Then, the bacterial GTBD R220 phylogenetic tree was downloaded from the GTDB webpage (https://data.gtdb.aau.ecogenomic.org/)^142^, loaded in R and pruned to branches in which GTDB identifiers matched those of the global/marine genome collection. Branches were collapsed at the order level, keeping only orders with ≥ 50 (global) or ≥ 10 (marine) genomes. Operations were done with the R packages treeio^144^ and APE^145^. The percentage prevalences of genomes per order for the three queried functions were plotted onto the pruned GTDB tree using iTOL^146^.

### Statistical analysis

Statistical analysis was performed with the GaphPad Prism 10 software package (GraphPad Software, Inc, San Diego, CA USA). A one-way ANOVA was performed to compare the means of three or more groups. If there was a significant difference between the group means (*P*-value < 0.05) a Tukey’s post-hoc test was performed to identify which pairs of group means are different from each other (adjusted *P*-value < 0.05).

## Acknowledgements

We thank I. Kamp, D. Merten and T. Schäfer (Friedrich Schiller University Jena, Germany) for performing trace-element analyses, M. C. F. van Teeseling (Friedrich Schiller University Jena, Germany) for valuable insights into morphological heterogeneity in *Alphaproteobacteria*, M. L. Hammer (Leibniz-HKI, Jena, Germany) for critical feedback on the manuscript and for support with data analysis and visualization, and V. Hotter (Friedrich Schiller University Jena, Germany) for comments on the manuscript. We are grateful for stimulating discussions on cobalamin cross-feeding with E. Petre (Leibniz-Institut für Ostseeforschung Warnemünde, Germany), for input on cobalamin-dependent methyl transfer from S. Studenik (Friedrich Schiller University Jena, Germany) and for advice on anaerobic metabolism from M. Kündgen (Friedrich Schiller University Jena, Germany). We thank H.-J. Ruscheweyh (ETH Zurich, Switzerland) for bioinformatic support with mOTUs-db and N. Wohlfarth (Friedrich Schiller University Jena, Germany) for skillful technical laboratory assistance. We thank the Honours Programme at the Friedrich Schiller University Jena for their support. This work was continuously supported by the Jena School for Microbial Communication (JSMC). We acknowledge funding of this project from the Carl Zeiss Stiftung. This study was funded by the German Research Foundation (DFG, Deutsche Forschungsgemeinschaft) under Project-ID 239748522 (ChemBioSys) and Germany’s Excellence Strategy under Project-ID 390713860 - EXC 2051. Funded by the Landesgraduiertenstipendium of the Free State of Thuringia awarded by the Friedrich Schiller University Jena. S.S. acknowledges funding from the Swiss National Science Foundation (project 205320_215395).

## Author contributions

J.H., M.Sp., T.S., E.S. and C.J. designed the study. J.H., M.St., M.Sp., T.H., N.K. and D.A.N.-B., performed and analyzed experiments. C.-E.W., K.K., G.P. and S.S. contributed to the design of the research and helped supervise the project. J.H. and C.J. wrote the manuscript. All authors discussed the results and contributed to the final manuscript.

