## Supplementary Information for "The glycine betaine-cobalamin feedback loop drives cross-feeding between marine bacteria and algae"

**Supplementary Text**

Cobalt limitation in the context of this study

Under standard laboratory conditions it is impossible to achieve cobalt concentrations comparable to those in ocean water where they can be in the low picomolar range^1,2^. Reported cobalt concentrations in ultrapure (Milli-Q) water are considerably higher (0.31 to 0.9 nM) and still sufficient for Cbl biosynthesis^3^. We analyzed the cobalt concentration in the trace elements solution that we prepared with CoCl_2_ and without CoCl_2_ via inductively coupled plasma - optical emission spectrometry (ICP-OES) and confirmed that the cobalt concentration in the trace elements solution without CoCl_2_ is below the detection limit of this method (lower than 0.02 mg l^-1^ or 0.34 µM) (Supplementary figure S2). Thus, in the context of this study, we define conditions as cobalt-limited, if *P. inhibens* was cultivated in medium supplemented with a trace elements solution that did not contain CoCl_2_ for at least one passage and then transferred into medium that was again supplemented with a trace element solution that did not contain CoCl_2_ with or without washing of the cells before inoculation of the second passage. We define conditions as cobalt-replete, if *P. inhibens* was cultivated in medium supplemented with a trace elements solution that contained CoCl_2_, independent of the presence of CoCl_2_ in the trace element solution used for the medium of the pre-cultures.

**Supplementary Figures**

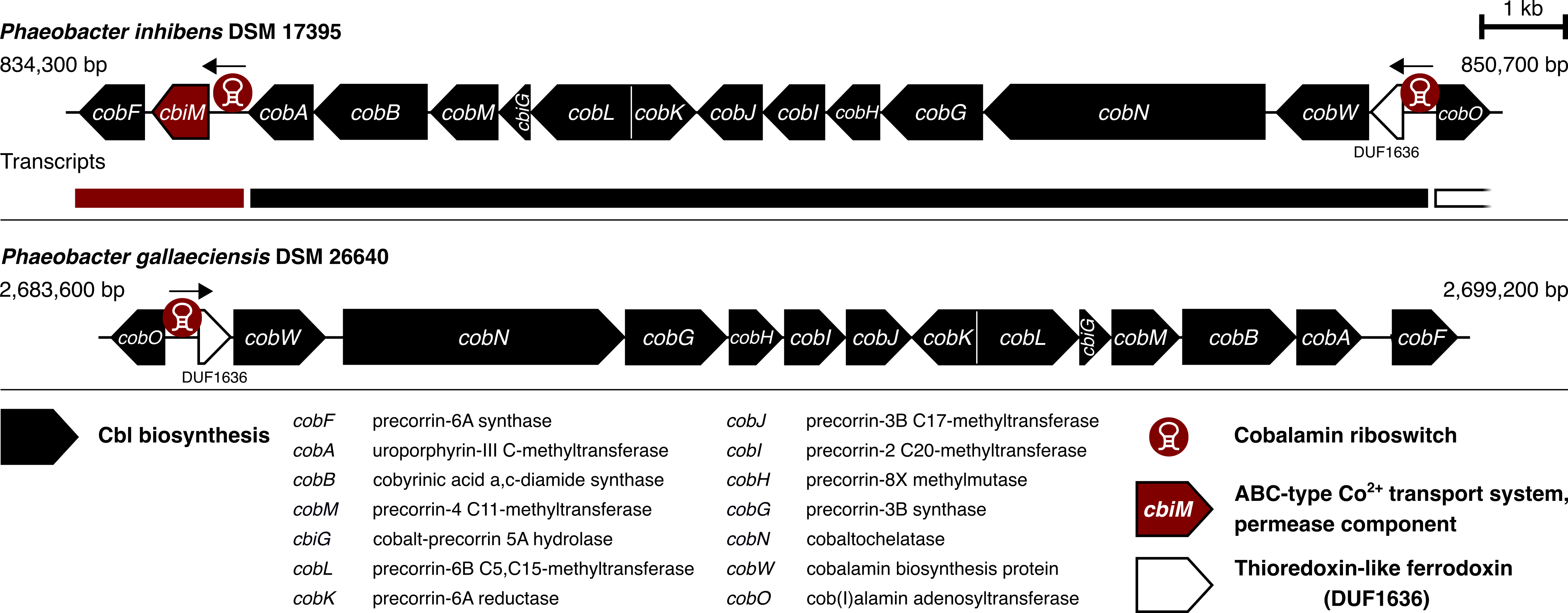

**Supplementary Fig. S1: Cbl biosynthesis gene clusters in *Phaeobacter inhibens* DSM 17395 and *Phaeobacter gallaeciensis* DSM 26640.** Cbl riboswitches post-transcriptionally regulate translation of Cbl biosynthesis operons and genes putatively involved in cobalt uptake. A putative cobalt permease is highlighted in red that is present in *P. inhibens* but missing in *P. gallaeciensis*. The latter possesses all genes encoding the consensus energy-coupling factor-type ABC cobalt transport system CbiMNQO (CbiM, GAL_RS07830) in a different locus with a Cbl riboswitch in its leader sequence and divergent to a putative TonB-dependent Cbl transporter and an adenosylcobinamide amidohydrolase encoding gene (*cbiZ*) that is involved in remodeling of Cbl. The locus is absent in *P. inhibens*.

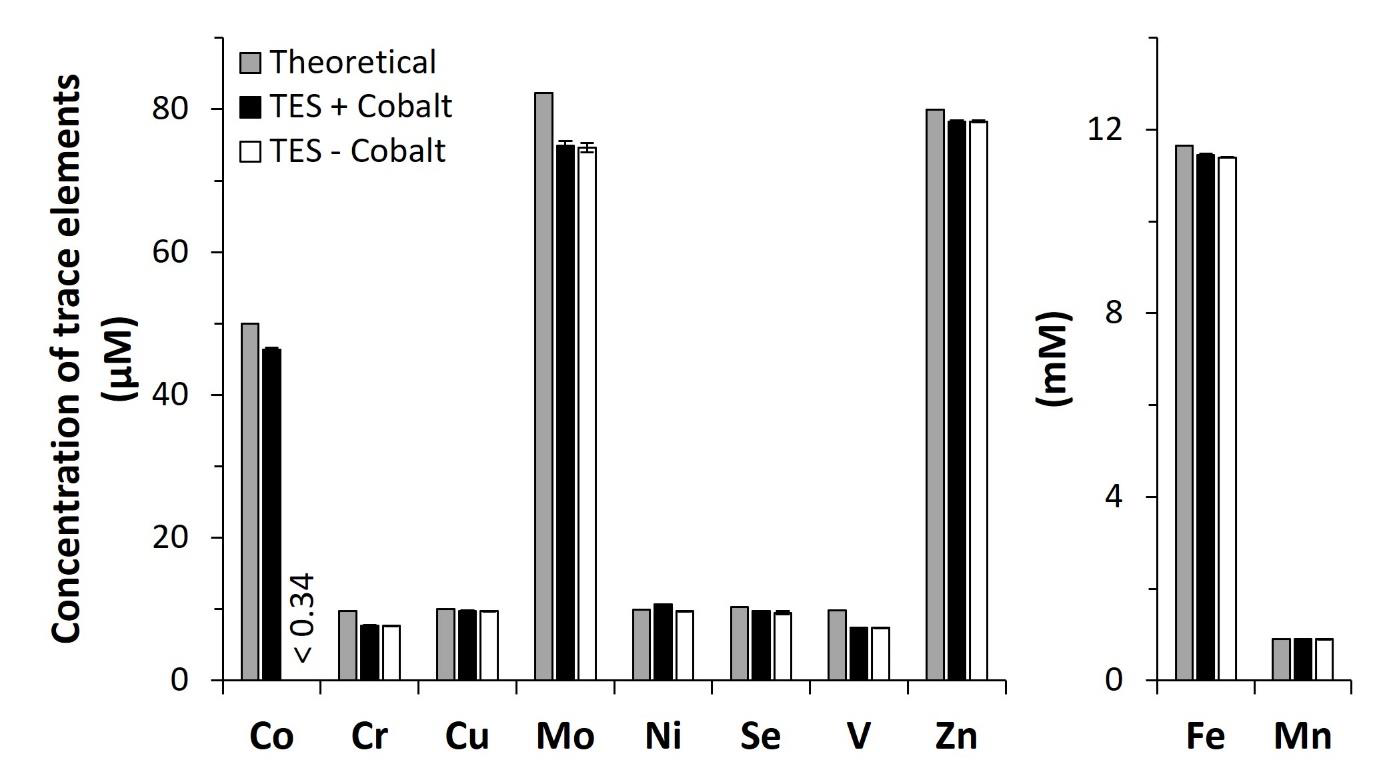

**Supplementary Fig. S2: Molar concentrations of ten trace elements in the trace elements solutions used for ASW medium preparation.** Theoretical concentrations were calculated from the recipe including Co_2_Cl. Trace elements solutions were prepared with Co_2_Cl (TES + Cobalt) and without it (TES - Cobalt). Molar concentrations were calculated from mass concentrations determined via ICP-OES. If a trace element was not detected the detection limit is displayed in the graph. ICP-OS was performed in technical triplicates. The error bars represent the standard deviation (n = 3).

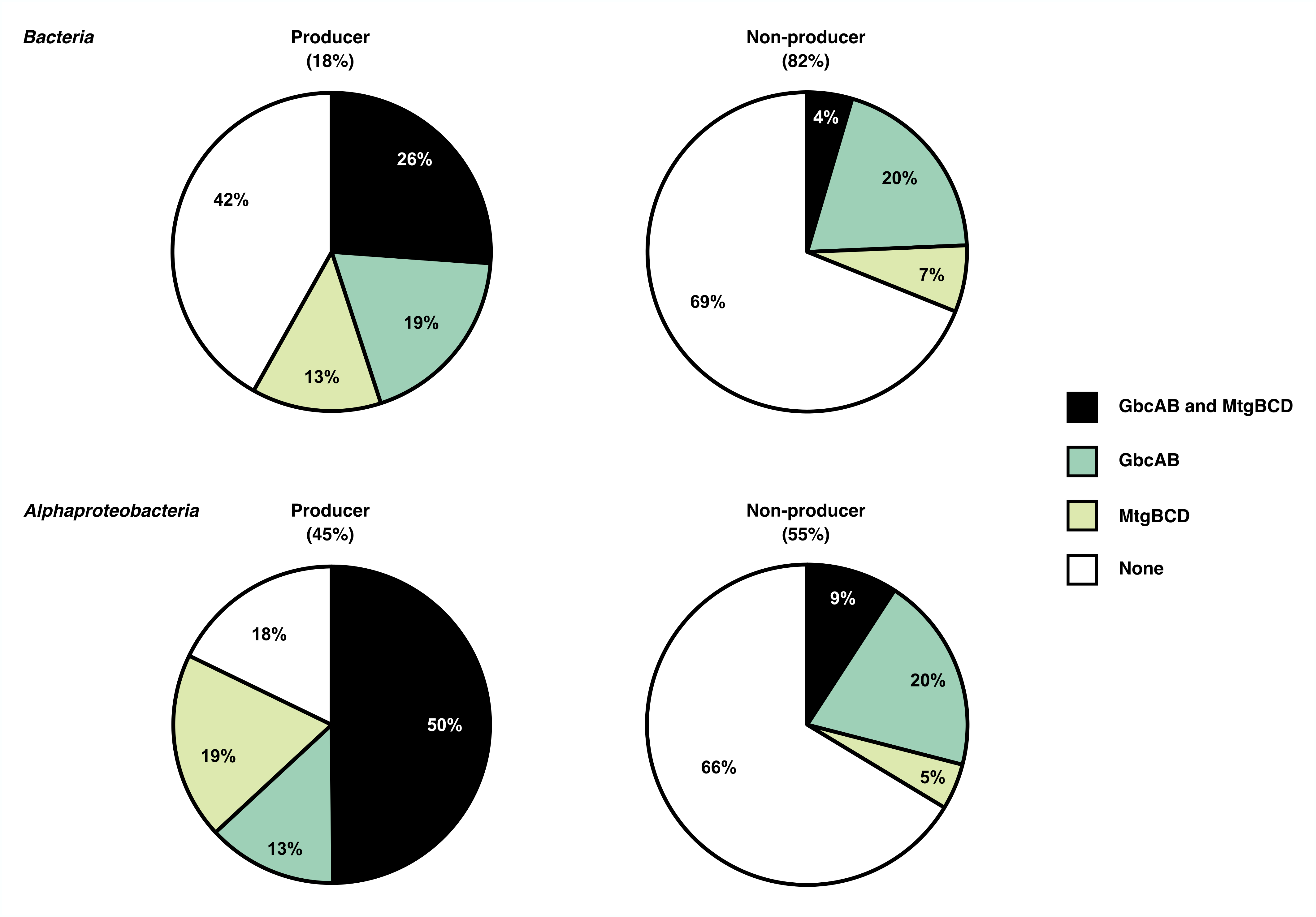

**Supplementary Fig. S3: Distribution of Cbl-dependent (MtgBCD) and -independent (GbcAB) GB demethylation among Cbl-producing and non-producing marine bacteria.** A database containing 15,525 dereplicated, high-quality, species-level genomes of marine bacteria (isolate genomes, metagenome assembled genomes and single-cell amplified genomes) was searched for the presence of protein families (Pfams) associated with Cbl-dependent and -independent GB demethylases. The two functions were considered as present if all Pfams of MtgBCD or GbcAB, respectively, were detected. Following Sañudo-Wilhelmy *et al.* (2014), we further queried 16 marker genes involved in Cbl biosynthesis and considered the pathway as present in a genome when > 75% of the corresponding Pfams were detected^4^.The percentage of bacteria predicted to be able (producer) and unable (non-producer) to biosynthesize Cbl *de novo* is indicated in brackets. The pie charts represent the percentages of producers and non-producers that possess either GbcAB or MtgBCD, both enzymes, or neither. Results are displayed for all marine bacteria included in the database (first row) and for a subset of bacteria of the class *Alphaproteobacteria* (second row)

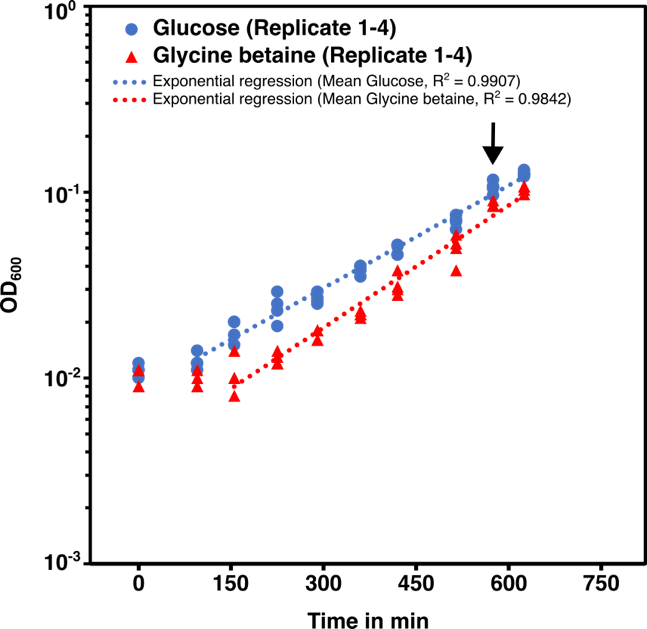
**Supplementary Fig. S4: Sampling point of *Phaeobacter inhibens* cultures for RNA extraction.** Cells sampled from the exponential phase of *P. inhibens* grown in pure cultures with 5.5 mM glucose (blue) or with 5.5 mM GB (red). Cultivation was done in four biological replicates (n = 4). The arrow indicates the timepoint of cell harvest for RNA extraction.

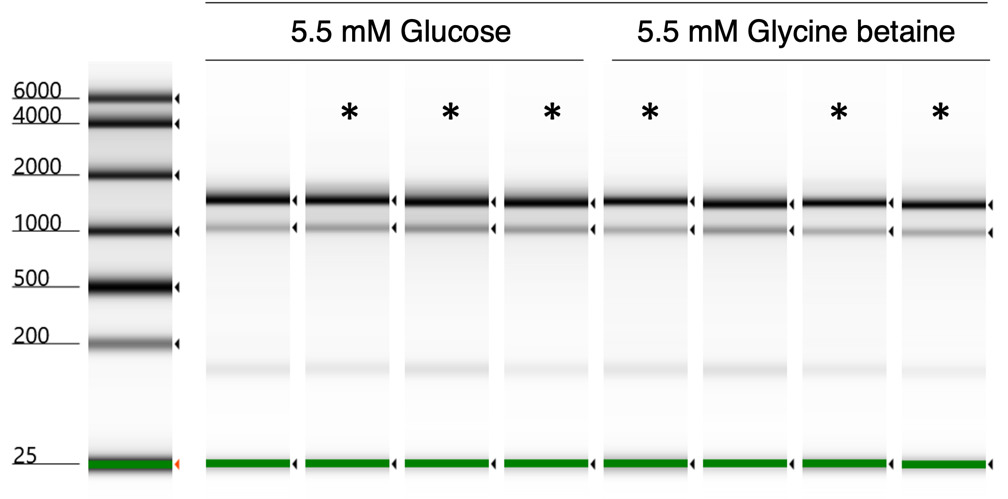

| **Sample ID** | **1** | **2** | **3** | **4** | **5** | **6** | **7** | **8** |
| --- | --- | --- | --- | --- | --- | --- | --- | --- |
| **Condition** | Glucose | Glucose | Glucose | Glucose | Glycine betaine | Glycine betaine | Glycine  betaine | Glycine betaine |
| **Qubit [ng/µl]** | 220.5 | 630 | 575 | 665 | 147.5 | 283 | 127.5 | 166.5 |
| **Tapestation [ng/µl]** | 139 | 429 | 558 | 474 | 87 | 170 | 85.2 | 104 |
| **Tapestation (RIN)** | 8.0 | 8.1 | 8.2 | 8.2 | 8.0 | 8.2 | 8.1 | 8.3 |

**Supplementary Fig. S5: Quality control of RNA extracted from exponential *P. inhibens* cultures for RNA-sequencing.** Tapestation intensities were scaled to samples. Ribosomal subunits are indicated by arrows. RIN-RNA integrity numbers were determined by the Tapestation software. Asterisks indicate samples selected for sequencing

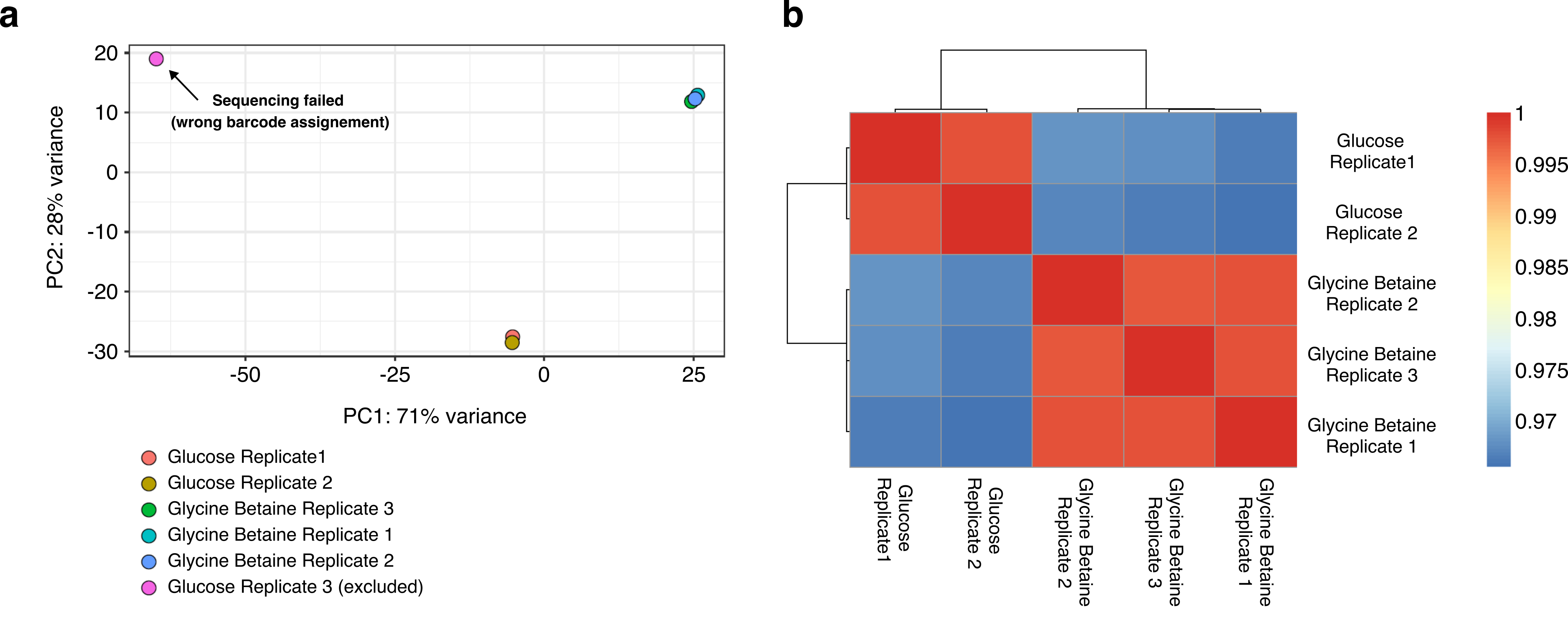

**Supplementary Fig. S6: Characterization of *P. inhibens* gene transcription profiles with glycine betaine and glucose.**

**a,** The PCA plot was generated using the DESeq2::plotPCA function with default options (ntop=500; top 500 variable genes) and DESeq2::rlog transformed read counts (blind=TRUE) as input. **b,** The heat map was generated using sample-to-sample Pearson correlation coefficients (base R; cor function) of the rlog transformed read counts as input for the pheatmap package.

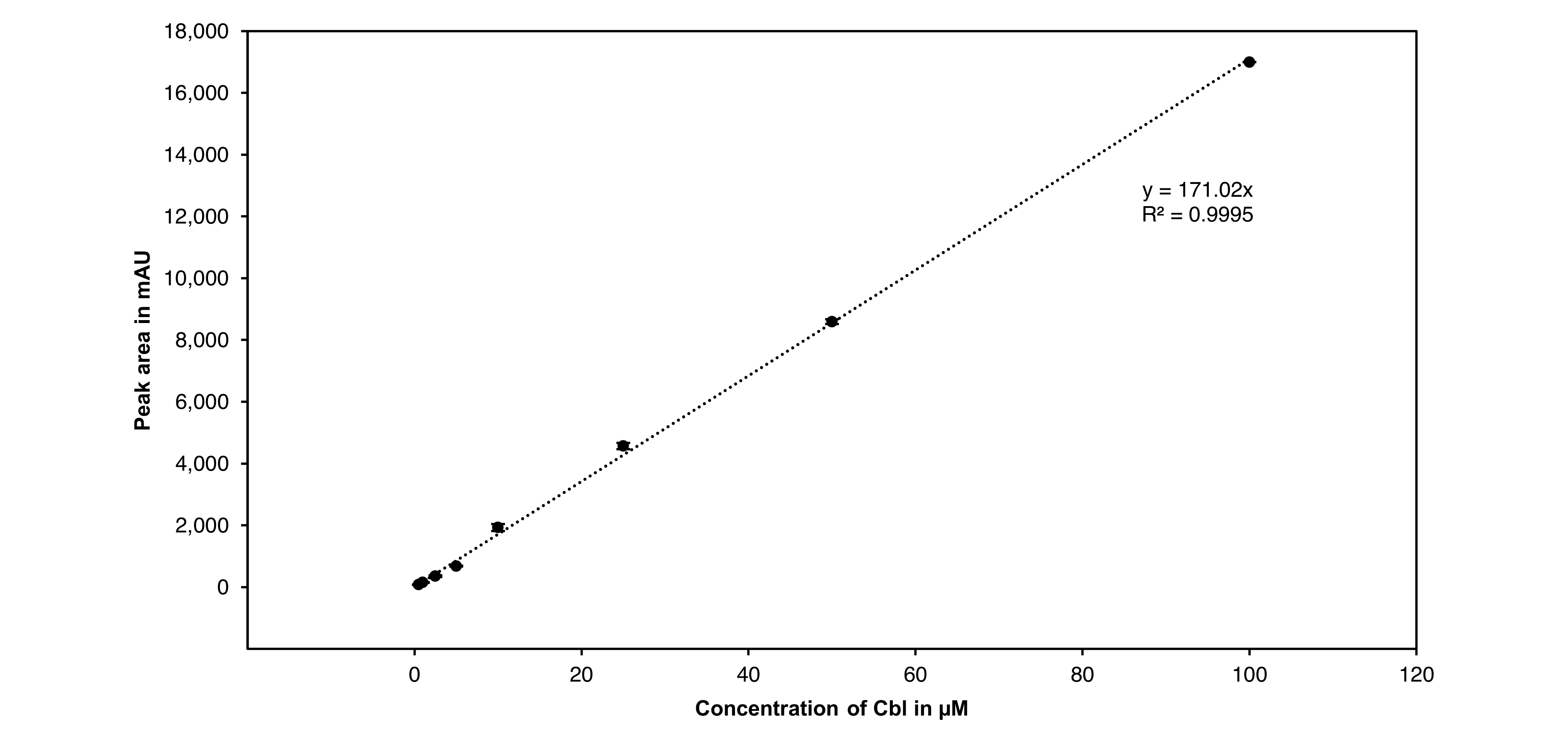

**Supplementary Fig. S7: Calibration curve for Cbl quantification.** Cbl standards were prepared as two independent dilution series and analyzed using HPLC measurement. Area of the Cbl peaks, identified by retention time and absorbance spectrum (Supplementary Fig. S8), are plotted against the concentration of the standards. The line equation and coefficient of determination (R^2^) of the linear regression are displayed in the graph.

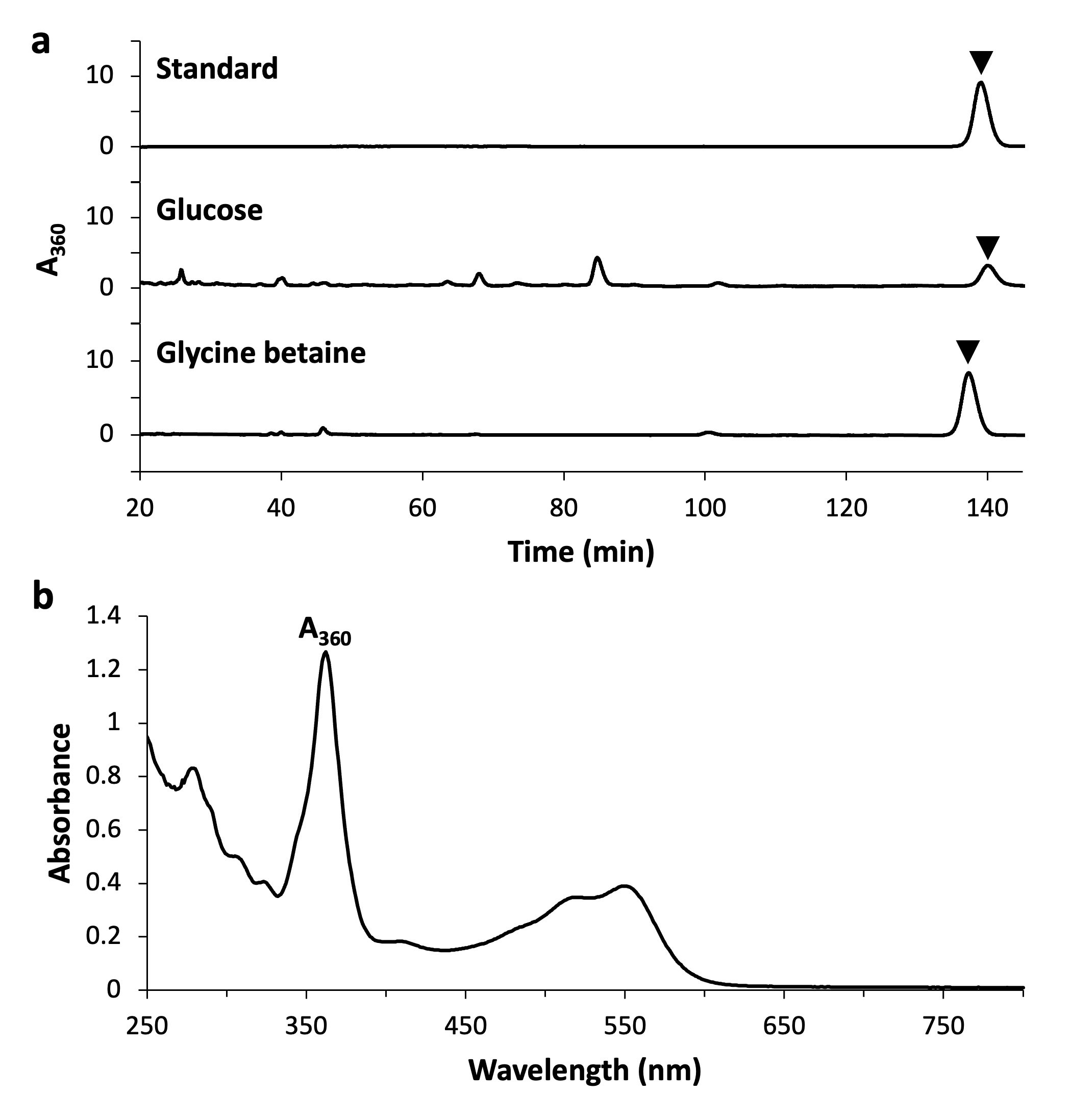

**Supplementary Fig. S8: Analyses of Cbl via HPLC with a UV-Vis detector.** (A) Exemplary chromatograms of a Cbl standard solution and Cbl purified from *P. inhibens* cultivated in ASW_v.2a_ medium with 5.5 mM glucose or 5.5 mM betaine. The purified Cbl extracts were applied undiluted (glucose) or 1:5 diluted (GB). The absorbance at 360 nm (A_360_) was measured. In the purified extracts the Cbl was identified by its retention time in comparison to the standard as well as by its characteristic absorbance spectrum between 200 and 600 nm. Black triangles indicate the Cbl peaks, and the respective peak area was used for quantification. All peaks that are not highlighted did not show the characteristic absorbance spectrum. (B) UV-Vis absorbance spectrum of a Cbl standard in the oxidized state [CoIII] with an absorbance maximum at approximately 360 nm.

**Supplementary Tables**

**Supplementary Table S1: Gene accession numbers and annotations**

| **RefSeq accession** | **Old accession** | **Gene ID** | **RefSeq_Product** | **Submitter_Product** | **Kegg_Product** |
| --- | --- | --- | --- | --- | --- |
| PGA1_RS05110 | PGA1_c10290 | *ompW* | outer membrane beta-barrel protein | outer membrane protein | ompW; outer membrane protein |
| PGA1_RS16700 | PGA1_c33650 | *BCCT* | BCCT family transporter | putative transporter, BCCT family |  |
| PGA1_RS06660 | PGA1_c13370 | *mtgE* | betaine--homocysteine S-methyltransferase | putative homocysteine S-methyltransferase | metH, MTR; 5-methyltetrahydrofolate--homocysteine methyltransferase [EC:2.1.1.13] |
| PGA1_RS02485 | PGA1_c05010 | *gbcA* | aromatic ring-hydroxylating dioxygenase subunit alpha | putative dioxygenase | stc2, hpbB; stachydrine N-demethylase [EC:1.14.13.247] |
| PGA1_RS02490 | PGA1_c05020 | *gbcB* | hybrid-cluster NAD(P)-dependent oxidoreductase | phenylacetic acid degradation NADH oxidoreductase | stc4, hpbC; stachydrine N-demethylase, reductase component |
| PGA1_RS07270 | PGA1_c14630 | *mtgB1* | trimethylamine methyltransferase family protein | trimethylamine methyltransferase v | mttB; trimethylamine---corrinoid protein Co-methyltransferase [EC:2.1.1.250] |
| PGA1_RS08190 | PGA1_c16500 | *mtgB2* | trimethylamine methyltransferase family protein | trimethylamine methyltransferase MttB | mttB; trimethylamine---corrinoid protein Co-methyltransferase [EC:2.1.1.250] |
| PGA1_RS12685 | PGA1_c25550 | *mtgB3* | trimethylamine methyltransferase family protein | trimethylamine methyltransferase MttB | mttB; trimethylamine---corrinoid protein Co-methyltransferase [EC:2.1.1.250] |
| PGA1_RS04260 | PGA1_c08570 | *mtgB4* | trimethylamine methyltransferase family protein | putative trimethylamine methyltransferase | mttB; trimethylamine---corrinoid protein Co-methyltransferase [EC:2.1.1.250] |
| PGA1_RS06650 | PGA1_c13350 | *mtgC* | cobalamin-binding protein | putative dimethylamine corrinoid protein |  |
| PGA1_RS07550 | PGA1_c15200 | *ramA* | ASKHA domain-containing protein | hypothetical protein |  |
| PGA1_RS06645 | PGA1_c13340 | DUF1638 | DUF1638 domain-containing protein | hypothetical protein |  |
| PGA1_RS07955 | PGA1_c16040 | *mtgD* | methyltetrahydrofolate cobalamin methyltransferase | pterin binding domain-containing protein | metH, MTR; 5-methyltetrahydrofolate--homocysteine methyltransferase [EC:2.1.1.13] |
| PGA1_RS05830 | PGA1_c11700 | *metF1* | methylenetetrahydrofolate reductase [NAD(P)H] | 5,10-methylenetetrahydrofolate reductase MetF | metF, MTHFR; methylenetetrahydrofolate reductase (NADH) [EC:1.5.1.54] |

**Table S1 (Continued)**

| **RefSeq accession** | **Old accession** | **Gene ID** | **RefSeq_Product** | **Submitter_Product** | **Kegg_Product** |
| --- | --- | --- | --- | --- | --- |
| PGA1_RS07960 | PGA1_c16050 | *metF2* | methylenetetrahydrofolate reductase | hypothetical protein | metF, MTHFR; methylenetetrahydrofolate reductase (NADH) [EC:1.5.1.54] |
| PGA1_RS14040 | PGA1_c28230 | *dmgdh* | FAD-dependent oxidoreductase | dimethylglycine dehydrogenase DmgdH | DMGDH; dimethylglycine dehydrogenase [EC:1.5.8.4] |
| PGA1_RS09505 | PGA1_c19150 | *soxA* | sarcosine oxidase subunit alpha family protein | sarcosine oxidase subunit alpha | soxA; sarcosine oxidase, subunit alpha [EC:1.5.3.24 1.5.3.1] |
| PGA1_RS09515 | PGA1_c19170 | *soxB* | sarcosine oxidase subunit beta family protein | sarcosine oxidase subunit beta | soxB; sarcosine oxidase, subunit beta [EC:1.5.3.24 1.5.3.1] |
| PGA1_RS09510 | PGA1_c19160 | *soxD* | sarcosine oxidase subunit delta | sarcosine oxidase 2 subunit delta | soxD; sarcosine oxidase, subunit delta [EC:1.5.3.24 1.5.3.1] |
| PGA1_RS09500 | PGA1_c19140 | *soxG* | sarcosine oxidase subunit gamma | sarcosine oxidase-like protein | soxG; sarcosine oxidase, subunit gamma [EC:1.5.3.24 1.5.3.1] |
| PGA1_RS02985 | PGA1_c06010 | *folD* | bifunctional methylenetetrahydrofolate dehydrogenase/methenyltetrahydrofolate cyclohydrolase FolD | bifunctional protein FolD | folD; methylenetetrahydrofolate dehydrogenase (NADP+) / methenyltetrahydrofolate cyclohydrolase [EC:1.5.1.5 3.5.4.9] |
| PGA1_RS02975 | PGA1_c05990 | *fhs* | formate--tetrahydrofolate ligase | formate--tetrahydrofolate ligase Fhs | fhs; formate--tetrahydrofolate ligase [EC:6.3.4.3] |
| PGA1_RS12780 | PGA1_c25740 | *fdwA* | formate dehydrogenase subunit alpha | formate dehydrogenase H | fdoG, fdhF, fdwA; formate dehydrogenase major subunit [EC:1.17.1.9] |
| PGA1_RS12785 | PGA1_c25750 | *fdwB* | NAD(P)H-dependent oxidoreductase subunit E | NADH-quinone oxidoreductase subunits E/F (fused) | fdwB; formate dehydrogenase beta subunit [EC:1.17.1.9] |
| PGA1_RS02370 | PGA1_c04780 | *metK* | methionine adenosyltransferase | S-adenosylmethionine synthetase MetK | metK, MAT; S-adenosylmethionine synthetase [EC:2.5.1.6] |
| PGA1_RS05915 | PGA1_c11870 | *glyA* | serine hydroxymethyltransferase | serine hydroxymethyltransferase GlyA | glyA, SHMT; glycine hydroxymethyltransferase [EC:2.1.2.1] |
| PGA1_RS11830 | PGA1_c23770 | *tdcG* | L-serine ammonia-lyase | L-serine dehydratase TdcG | E4.3.1.17, sdaA, sdaB, tdcG; L-serine dehydratase [EC:4.3.1.17] |
| PGA1_RS10060 | PGA1_c20350 | *hemA* | 5-aminolevulinate synthase | 5-aminolevulinate synthase HemA | E2.3.1.37, ALAS; 5-aminolevulinate synthase [EC:2.3.1.37] |
| PGA1_RS07400 | PGA1_c14900 | *hemB* | porphobilinogen synthase | delta-aminolevulinic acid dehydratase HemB | hemB, ALAD; porphobilinogen synthase [EC:4.2.1.24] |

**Supplementary Table S1 (Continued)**

| **RefSeq accession** | **Old accession** | **Gene ID** | **RefSeq_Product** | **Submitter_Product** | **Kegg_Product** |
| --- | --- | --- | --- | --- | --- |
| PGA1_RS14885 | PGA1_c29950 | *hemC* | hydroxymethylbilane synthase | porphobilinogen deaminase HemC | hemC, HMBS; hydroxymethylbilane synthase [EC:2.5.1.61] |
| PGA1_RS17630 | PGA1_c35500 | *hemD* | uroporphyrinogen-III synthase | uoporphyrinogen-III synthase HemD-like protein | hemD, UROS; uroporphyrinogen-III synthase [EC:4.2.1.75] |
| PGA1_RS04010 | PGA1_c08070 | *cobA1* | uroporphyrinogen-III C-methyltransferase | uroporphyrin-III C-methyltransferase CobA | cobA; uroporphyrin-III C-methyltransferase [EC:2.1.1.107] |
| PGA1_RS04045 | PGA1_c08140 | *cobI/cbiL* | precorrin-2 C(20)-methyltransferase | precorrin-2 C(20)-methyltransferase CobI | cobI-cbiL; precorrin-2/cobalt-factor-2 C20-methyltransferase [EC:2.1.1.130 2.1.1.151] |
| PGA1_RS04055 | PGA1_c08160 | *cobG* | precorrin-3B synthase | putative precorrin-3B synthase | cobG; precorrin-3B synthase [EC:1.14.13.83] |
| PGA1_RS04040 | PGA1_c08130 | *cobJ/cbiH* | precorrin-3B C(17)-methyltransferase | precorrin-3B C(17)-methyltransferase CobJ | cobJ, cbiH; precorrin-3B C17-methyltransferase / cobalt-factor III methyltransferase [EC:2.1.1.131 2.1.1.272] |
| PGA1_RS04020 | PGA1_c08090 | *cobM/cbiF* | precorrin-4 C(11)-methyltransferase | precorrin-4 C(11)-methyltransferase CobM | cobM, cbiF; precorrin-4/cobalt-precorrin-4 C11-methyltransferase [EC:2.1.1.133 2.1.1.271] |
| PGA1_RS04000 | PGA1_c08050 | *cobF* | precorrin-6A synthase (deacetylating) | putative precorrin-6A synthase (deacetylating), cobF | cobF; precorrin-6A synthase [EC:2.1.1.152] |
| PGA1_RS04025 | PGA1_c08100 | *cbiG* | cobalamin biosynthesis protein | hypothetical protein | cbiG; cobalt-precorrin 5A hydrolase [EC:3.7.1.12] |
| PGA1_RS04035 | PGA1_c08120 | *cobK/cbiJ* | cobalt-precorrin-6A reductase | precorrin-6A reductase CobK | cobK-cbiJ; precorrin-6A/cobalt-precorrin-6A reductase [EC:1.3.1.54 1.3.1.106] |
| PGA1_RS04030 | PGA1_c08110 | *cobL/cbiET* | bifunctional cobalt-precorrin-7 (C(5))-methyltransferase/cobalt-precorrin-6B (C(15))-methyltransferase | precorrin-6Y C(5,15)-methyltransferase | cobL-cbiET; precorrin-6B C5,15-methyltransferase / cobalt-precorrin-6B C5,C15-methyltransferase [EC:2.1.1.132 2.1.1.289 2.1.1.196] |
| PGA1_RS04050 | PGA1_c08150 | *cobH/cbiA* | precorrin-8X methylmutase | precorrin-8X methylmutase CobH | cobH-cbiC; precorrin-8X/cobalt-precorrin-8 methylmutase [EC:5.4.99.61 5.4.99.60] |
| PGA1_RS04015 | PGA1_c08080 | *cobB/cbiA* | cobyrinate a,c-diamide synthase | putative cobyrinic acid A,C-diamide synthase CobB | cobB-cbiA; cobyrinic acid a,c-diamide synthase [EC:6.3.5.9 6.3.5.11] |

**Supplementary Table S1 (Continued)**

| **RefSeq accession** | **Old accession** | **Gene ID** | **RefSeq_Product** | **Submitter_Product** | **Kegg_Product** |
| --- | --- | --- | --- | --- | --- |
| PGA1_RS04005 | PGA1_c08060 | *cbiM* | energy-coupling factor ABC transporter permease | hypothetical protein |  |
| PGA1_RS04060 | PGA1_c08170 | *cobN* | cobaltochelatase subunit CobN | aerobic cobaltochelatase subunit CobN | cobN; cobaltochelatase CobN [EC:6.6.1.2] |
| PGA1_RS04860 | PGA1_c09780 | *cobS* | cobaltochelatase subunit CobS | aerobic cobaltochelatase subunit CobS | cobS; cobaltochelatase CobS [EC:6.6.1.2] |
| PGA1_RS04850 | PGA1_c09770 | *cobT* | cobaltochelatase subunit CobT | aerobic cobaltochelatase subunit CobT | cobT; cobaltochelatase CobT [EC:6.6.1.2] |
| PGA1_RS04065 | PGA1_c08180 | *cobW* | cobalamin biosynthesis protein CobW | cobalamin biosynthesis protein CobW | cobW; cobalamin biosynthesis protein CobW |
| PGA1_RS04070 | PGA1_c08190 | DUF1636 | DUF1636 domain-containing protein | conserved hypothetical protein (DUF1636) |  |
| PGA1_RS04075 | PGA1_c08200 | *cobO* | cob(I)yrinic acid a,c-diamide adenosyltransferase | cob(I)yrinic acid a,c-diamide adenosyltransferase CobO | cobA, btuR; cob(I)alamin adenosyltransferase [EC:2.5.1.17] |
| PGA1_RS00620 | PGA1_c01260 | *pduO* | cob(I)yrinic acid a,c-diamide adenosyltransferase | putative cobalamin adenosyltransferase | MMAB, pduO; cob(I)alamin adenosyltransferase [EC:2.5.1.17] |
| PGA1_RS11940 | PGA1_c24000 | *cobQ/cbiP* | cobyric acid synthase | cobyric acid synthase CobQ | cobQ, cbiP; adenosylcobyric acid synthase [EC:6.3.5.10] |
| PGA1_RS00405 | PGA1_c00830 | *cobC* | threonine-phosphate decarboxylase CobD | putative threonine-phosphate decarboxylase CobC | cobC1, cobC; cobalamin biosynthesis protein CobC |
| PGA1_RS00400 | PGA1_c00820 | *cobD/cbiB* | adenosylcobinamide-phosphate synthase CbiB | putative cobalamin biosynthesis protein CobD | cbiB, cobD; adenosylcobinamide-phosphate synthase [EC:6.3.1.10] |
| PGA1_RS13250 | PGA1_c26670 | *cobP* | bifunctional adenosylcobinamide kinase/adenosylcobinamide-phosphate guanylyltransferase | bifunctional adenosylcobalamin biosynthesis protein CobP | cobP, cobU; adenosylcobinamide kinase / adenosylcobinamide-phosphate guanylyltransferase [EC:2.7.1.156 2.7.7.62] |
| PGA1_RS00275 | PGA1_c00550 | *bluB* | 5,6-dimethylbenzimidazole synthase | putative cob(II)yrinic acid a,c-diamide reductase | bluB; 5,6-dimethylbenzimidazole synthase [EC:1.13.11.79] |
| PGA1_RS03760 | PGA1_c07570 | *cobU* | nicotinate-nucleotide--dimethylbenzimidazole phosphoribosyltransferase | nicotinate-nucleotide-dimethylbenzimidazole phosphoribosyltransferase CobT | E2.4.2.21, cobU, cobT; nicotinate-nucleotide--dimethylbenzimidazole phosphoribosyltransferase [EC:2.4.2.21] |
| PGA1_RS13255 | PGA1_c26680 | *cobC/ phpB* | histidine phosphatase family protein | putative phosphoglycerate mutase family protein | cobC, phpB; alpha-ribazole phosphatase [EC:3.1.3.73] |

**Supplementary Table S1 (Continued)**

| **RefSeq accession** | **Old accession** | **Gene ID** | **RefSeq_Product** | **Submitter_Product** | **Kegg_Product** |
| --- | --- | --- | --- | --- | --- |
| PGA1_RS03755 | PGA1_c07560 | *cobV* | adenosylcobinamide-GDP ribazoletransferase | putative cobalamin-5-phosphate synthase | E2.7.8.26, cobS, cobV; adenosylcobinamide-GDP ribazoletransferase [EC:2.7.8.26] |
| PGA1_RS14880 | PGA1_c29940 | *hemE* | uroporphyrinogen decarboxylase | uroporphyrinogen decarboxylase HemE | hemE, UROD; uroporphyrinogen decarboxylase [EC:4.1.1.37] |
| PGA1_RS00180 | PGA1_c00360 | *hemN* | oxygen-independent coproporphyrinogen III oxidase | oxygen-independent coproporphyrinogen III oxidase HemN | hemN, hemZ; oxygen-independent coproporphyrinogen III oxidase [EC:1.3.98.3] |
| PGA1_RS14895 | PGA1_c29970 | *hemF* | oxygen-dependent coproporphyrinogen oxidase | coproporphyrinogen 3 oxidase, aerobic | CPOX, hemF; coproporphyrinogen III oxidase [EC:1.3.3.3] |
| PGA1_RS17640 | PGA1_c35520 | *hemY* | tetratricopeptide repeat protein | HemY-like protein | hemY; HemY protein |
| PGA1_RS17250 | PGA1_c34740 | *hemH* | ferrochelatase | ferrochelatase HemH | hemH, FECH; protoporphyrin/coproporphyrin ferrochelatase [EC:4.98.1.1 4.99.1.9] |
| PGA1_RS02820 | PGA1_c05690 | *ctb* | group III truncated hemoglobin | hypothetical protein | glbN; hemoglobin |
| PGA1_RS07200 | PGA1_c14480 | *speD* | adenosylmethionine decarboxylase | S-adenosylmethionine decarboxylase SpeH | speD, AMD1; S-adenosylmethionine decarboxylase [EC:4.1.1.50] |
| PGA1_RS07195 | PGA1_c14470 | *speE* | polyamine aminopropyltransferase | spermidine synthase SpeE | speE, SRM, SPE3; spermidine synthase [EC:2.5.1.16] |
| PGA1_RS08565 | PGA1_c17250 | *gapA1* | type I glyceraldehyde-3-phosphate dehydrogenase | glyceraldehyde-3-phosphate dehydrogenase 2 | GAPDH, gapA; glyceraldehyde 3-phosphate dehydrogenase (phosphorylating) [EC:1.2.1.12] |
| PGA1_RS02730 | PGA1_c05510 | *gapA2* | glyceraldehyde-3-phosphate dehydrogenase | glyceraldehyde-3-phosphate dehydrogenase 1 | GAPDH, gapA; glyceraldehyde 3-phosphate dehydrogenase (phosphorylating) [EC:1.2.1.12] |
| PGA1_RS19445 | PGA1_78p00290 | *gcvT* | glycine cleavage system aminomethyltransferase GcvT | aminomethyltransferase GcvT | gcvT, AMT; aminomethyltransferase [EC:2.1.2.10] |
| PGA1_RS19450 | PGA1_78p00300 | *gcvH1* | glycine cleavage system protein GcvH | glycine cleavage system protein GcvH | gcvH, GCSH; glycine cleavage system H protein |
| PGA1_RS02760 | PGA1_c05560 | *gcvH2* | glycine cleavage system protein GcvH | glycine cleavage system protein GcvH | gcvH, GCSH; glycine cleavage system H protein |

**Supplementary Table S1 (Continued)**

| **RefSeq accession** | **Old accession** | **Gene ID** | **RefSeq_Product** | **Submitter_Product** | **Kegg_Product** |
| --- | --- | --- | --- | --- | --- |
| PGA1_RS19455 | PGA1_78p00310 | *gcvP1* | aminomethyl-transferring glycine dehydrogenase | glycine dehydrogenase | GLDC, gcvP; glycine dehydrogenase [EC:1.4.4.2] |
| PGA1_RS02765 | PGA1_c05570 | *gcvP2* | aminomethyl-transferring glycine dehydrogenase |  | GLDC, gcvP; glycine dehydrogenase [EC:1.4.4.2] |
| PGA1_RS12185 | PGA1_c24490 | *mce* | methylmalonyl-CoA epimerase | methylmalonyl-CoA epimerase | MCEE, epi; methylmalonyl-CoA/ethylmalonyl-CoA epimerase [EC:5.1.99.1] |
| PGA1_RS10655 | PGA1_c21510 | *mut* | methylmalonyl-CoA mutase | methylmalonyl-CoA mutase BhbA | MUT; methylmalonyl-CoA mutase [EC:5.4.99.2] |
| PGA1_RS09040 | PGA1_c18210 | *late control protein D* | late control protein D | phage late control gene D protein (GPD) | K06905; uncharacterized protein |
| PGA1_RS09050 | PGA1_c18230 | *tail protein* | phage tail protein | phage-related tail protein (gpU) | K06906; uncharacterized protein |
| PGA1_RS09055 | PGA1_c18240 | *tail tape measure p.* | phage tail tape measure protein | putative phage tail tape measure protein |  |
| PGA1_RS09060 | PGA1_c18250 | *tail assembly protein* | phage tail assembly protein | hypothetical protein |  |
| PGA1_RS09065 | PGA1_c18260 | *major tail tube protein* | phage major tail tube protein | phage major tail tube protein | K06908; uncharacterized protein |
| PGA1_RS09070 | PGA1_c18270 | *tail sheath* | phage tail sheath subtilisin-like domain-containing protein | major tail sheath protein | K06907; uncharacterized protein |
| PGA1_RS09095 | PGA1_c18320 | *tail protein* | phage tail protein I | phage tail protein I |  |
| PGA1_RS09100 | PGA1_c18330 | *baseplate assembly p. J* | baseplate J/gp47 family protein | baseplate assembly protein J |  |
| PGA1_RS19860 | PGA1_c18350 | *baseplate assembly p. V* | phage baseplate assembly protein V | putative baseplate assembly protein V |  |
| PGA1_RS09140 | PGA1_c18400 | *major capsid protein* | phage major capsid protein | peptidase U35, phage prohead HK97-like protein |  |

**Supplementary Table S1 (Continued)**

| **RefSeq accession** | **Old accession** | **Gene ID** | **RefSeq_Product** | **Submitter_Product** | **Kegg_Product** |
| --- | --- | --- | --- | --- | --- |
| PGA1_RS09145 | PGA1_c18410 | *portal protein* | phage portal protein | phage portal protein, lambda family |  |
| PGA1_RS09165 | PGA1_c18450 | *spanin-like protein* | bacteriophage spanin2 family protein | hypothetical protein |  |
| PGA1_RS00615 | PGA1_c01250 | *fixA* | electron transfer flavoprotein subunit beta/FixA family protein | electron transfer flavoprotein subunit beta | fixA, etfB; electron transfer flavoprotein beta subunit |
| PGA1_RS00610 | PGA1_c01240 | *fixB* | electron transfer flavoprotein subunit alpha/FixB family protein | electron transfer flavoprotein subunit alpha | fixB, etfA; electron transfer flavoprotein alpha subunit |
| PGA1_RS01625 | PGA1_c03320 | *fixC* | electron transfer flavoprotein-ubiquinone oxidoreductase | electron transfer flavoprotein-ubiquinone oxidoreductase | ETFDH; electron-transferring-flavoprotein dehydrogenase [EC:1.5.5.1] |
| PGA1_RS14970 | PGA1_c30120 | *ccoN* | cytochrome-c oxidase, cbb3-type subunit I | cytochrome c oxidase subunit 1 | ccoN; cytochrome c oxidase cbb3-type subunit I [EC:7.1.1.9] |
| PGA1_RS14965 | PGA1_c30110 | *ccoO* | cytochrome-c oxidase, cbb3-type subunit II | cytochrome c oxidase subunit 2 | ccoO; cytochrome c oxidase cbb3-type subunit II |
| PGA1_RS14960 | PGA1_c30100 | *ccoQ* | CcoQ/FixQ family Cbb3-type cytochrome c oxidase assembly chaperone | cytochrome c oxidase subunit 4 | ccoQ; cytochrome c oxidase cbb3-type subunit IV |
| PGA1_RS14955 | PGA1_c30090 | *ccoP* | cytochrome-c oxidase, cbb3-type subunit III | cytochrome c oxidase subunit 3 | ccoP; cytochrome c oxidase cbb3-type subunit III |

**Supplementary Table S2: Gene accession number, log_2_(fold change) and adjusted *P*-values (padj).**

| **RefSeq accession** | **Old accession** | **Gene ID** | **Glycine betaine vs. Glucose** | | **Co-culture (growth) vs. Glucose** | | **Co-culture (growth) vs.**  **Co-culture (demise)** | | **Co-culture (day 9) vs.**  **Co-culture (day 4)** | |
| --- | --- | --- | --- | --- | --- | --- | --- | --- | --- | --- |
|  |  |  | **log_2_FC*** | **padj**** | **log_2_FC*** | **padj**** | **log_2_FC*** | **padj**** | **log_2_FC*** | **padj**** |
| PGA1_RS05110 | PGA1_c10290 | *ompW* | 2.003 | 0.000 | -0.216 | 0.812 | -0.730 | 0.223 | 6.152 | 0.000 |
| PGA1_RS16700 | PGA1_c33650 | *BCCT* | 3.838 | 0.000 | 3.019 | 0.000 | -1.390 | 0.002 | 0.144 | 0.990 |
| PGA1_RS06660 | PGA1_c13370 | *mtgE* | -0.702 | 0.000 | -1.426 | 0.000 | 0.507 | 0.121 | -0.062 | 0.995 |
| PGA1_RS02485 | PGA1_c05010 | *gbcA* | 8.579 | 0.000 | 5.632 | 0.000 | -4.084 | 0.000 | 0.175 | 0.383 |
| PGA1_RS02490 | PGA1_c05020 | *gbcB* | 6.503 | 0.000 | 2.745 | 0.003 | -0.946 | 0.170 | 2.673 | 0.013 |
| PGA1_RS07270 | PGA1_c14630 | *mtgB1* | 7.018 | 0.000 | 5.763 | 0.000 | -3.885 | 0.000 | 1.336 | 0.023 |
| PGA1_RS08190 | PGA1_c16500 | *mtgB2* | 2.073 | 0.000 | 0.500 | 0.531 | -0.580 | 0.382 | -0.248 | 0.676 |
| PGA1_RS12685 | PGA1_c25550 | *mtgB3* | -0.667 | 0.000 | 0.169 | 0.788 | 2.501 | 0.000 | -0.049 | 0.995 |
| PGA1_RS04260 | PGA1_c08570 | *mtgB4* | 2.881 | 0.000 | 0.516 | 0.482 | -0.094 | 0.888 | -0.052 | 0.995 |
| PGA1_RS06650 | PGA1_c13350 | *mtgC* | 3.635 | 0.000 | 0.966 | 0.010 | -2.307 | 0.000 | 0.149 | 0.990 |
| PGA1_RS07550 | PGA1_c15200 | *ramA* | 3.251 | 0.000 | 0.784 | 0.236 | -1.921 | 0.001 | 1.522 | 0.007 |
| PGA1_RS06645 | PGA1_c13340 | DUF1638 | 2.848 | 0.000 | 0.783 | 0.100 | -1.129 | 0.005 | 0.448 | 0.630 |
| PGA1_RS07955 | PGA1_c16040 | *mtgD* | 2.777 | 0.000 | 0.232 | 0.745 | -2.384 | 0.000 | 1.753 | 0.004 |
| PGA1_RS05830 | PGA1_c11700 | *metF1* | -3.582 | 0.000 | -3.635 | 0.001 | 2.193 | 0.028 | -0.090 | 0.946 |
| PGA1_RS07960 | PGA1_c16050 | *metF2* | 3.134 | 0.000 | 0.483 | 0.408 | -2.523 | 0.000 | 1.053 | 0.110 |
| PGA1_RS14040 | PGA1_c28230 | *dmgdh* | 7.128 | 0.000 | 5.523 | 0.000 | -4.255 | 0.000 | 1.087 | 0.112 |
| PGA1_RS09505 | PGA1_c19150 | *soxA* | 7.324 | 0.000 | 4.577 | 0.000 | -3.588 | 0.000 | 2.495 | 0.000 |
| PGA1_RS09515 | PGA1_c19170 | *soxB* | 7.733 | 0.000 | 5.502 | 0.000 | -4.250 | 0.000 | 1.579 | 0.054 |
| PGA1_RS09510 | PGA1_c19160 | *soxD* | 7.272 | 0.000 | 4.306 | 0.000 | -3.774 | 0.000 | 2.267 | 0.034 |
| PGA1_RS09500 | PGA1_c19140 | *soxG* | 6.970 | 0.000 | 3.993 | 0.000 | -2.245 | 0.002 | 2.634 | 0.000 |
| PGA1_RS02985 | PGA1_c06010 | *folD* | 3.794 | 0.000 | 1.672 | 0.000 | -3.774 | 0.000 | 1.332 | 0.004 |
| PGA1_RS02975 | PGA1_c05990 | *fhs* | 4.773 | 0.000 | 2.804 | 0.000 | -3.931 | 0.000 | 1.315 | 0.029 |
| PGA1_RS12780 | PGA1_c25740 | *fdwA* | 3.639 | 0.000 | 1.430 | 0.003 | -2.479 | 0.000 | 1.349 | 0.008 |
| PGA1_RS12785 | PGA1_c25750 | *fdwB* | 3.762 | 0.000 | 0.619 | 0.277 | -1.764 | 0.000 | 1.269 | 0.034 |

**Supplementary Table S2 (Continued)**

| **RefSeq accession** | **Old accession** | **Gene ID** | **Glycine betaine vs. Glucose** | | **Co-culture (growth) vs. Glucose** | | **Co-culture (growth) vs.**  **Co-culture (demise)** | | **Co-culture (day 9) vs.**  **Co-culture (day 4)** | |
| --- | --- | --- | --- | --- | --- | --- | --- | --- | --- | --- |
|  |  |  | **log_2_FC*** | **padj**** | **log_2_FC*** | **padj**** | **log_2_FC*** | **padj**** | **log_2_FC*** | **padj**** |
| PGA1_RS02370 | PGA1_c04780 | *metK* | -0.278 | 0.006 | -1.569 | 0.000 | -0.129 | 0.719 | 0.133 | 0.995 |
| PGA1_RS05915 | PGA1_c11870 | *glyA* | 2.087 | 0.000 | -0.800 | 0.384 | -0.280 | 0.638 | 0.079 | 0.945 |
| PGA1_RS11830 | PGA1_c23770 | *tdcG* | 4.992 | 0.000 | 0.260 | 0.711 | 0.025 | 0.976 | 7.487 | 0.001 |
| PGA1_RS10060 | PGA1_c20350 | *hemA* | 1.646 | 0.000 | 0.787 | 0.093 | -0.554 | 0.189 | 0.625 | 0.391 |
| PGA1_RS07400 | PGA1_c14900 | *hemB* | 1.208 | 0.000 | -0.329 | 0.617 | -0.555 | 0.284 | 0.095 | 0.995 |
| PGA1_RS14885 | PGA1_c29950 | *hemC* | 0.584 | 0.000 | 0.525 | 0.365 | -1.420 | 0.003 | 0.044 | 0.995 |
| PGA1_RS17630 | PGA1_c35500 | *hemD* | 2.567 | 0.000 | -0.378 | 0.673 | 0.563 | 0.445 | -0.028 | 0.995 |
| PGA1_RS04010 | PGA1_c08070 | *cobA1* | 1.243 | 0.000 | 0.008 | 0.995 | -0.656 | 0.304 | 0.377 | 0.332 |
| PGA1_RS04045 | PGA1_c08140 | *cobI/cbiL* | 1.824 | 0.000 | 1.186 | 0.044 | -2.971 | 0.000 | 0.186 | 0.946 |
| PGA1_RS04055 | PGA1_c08160 | *cobG* | 1.764 | 0.000 | 1.448 | 0.000 | -4.214 | 0.000 | 0.126 | 0.995 |
| PGA1_RS04040 | PGA1_c08130 | *cobJ/cbiH* | 1.618 | 0.000 | 0.756 | 0.240 | -1.783 | 0.002 | 1.232 | 0.158 |
| PGA1_RS04020 | PGA1_c08090 | *cobM/cbiF* | 1.941 | 0.000 | 0.556 | 0.422 | -1.824 | 0.002 | 1.152 | 0.205 |
| PGA1_RS04000 | PGA1_c08050 | *cobF* | 1.941 | 0.000 | 1.319 | 0.007 | -2.769 | 0.000 | 0.221 | 0.918 |
| PGA1_RS04025 | PGA1_c08100 | *cbiG* | 2.058 | 0.000 | 0.916 | 0.194 | -2.149 | 0.001 | 0.137 | 0.946 |
| PGA1_RS04035 | PGA1_c08120 | *cobK/cbiJ* | 0.844 | 0.000 | 0.261 | 0.713 | -0.720 | 0.186 | 0.630 | 0.391 |
| PGA1_RS04030 | PGA1_c08110 | *cobL/cbiET* | 1.362 | 0.000 | 0.764 | 0.274 | -1.712 | 0.005 | 1.094 | 0.179 |
| PGA1_RS04050 | PGA1_c08150 | *cobH/cbiA* | 1.715 | 0.000 | 0.679 | 0.233 | -2.552 | 0.000 | 0.121 | 0.990 |
| PGA1_RS04015 | PGA1_c08080 | *cobB/cbiA* | 1.469 | 0.000 | 0.314 | 0.694 | -1.192 | 0.074 | 2.724 | 0.001 |
| PGA1_RS04005 | PGA1_c08060 | *cbiM* | 3.557 | 0.000 | 0.640 | 0.398 | -6.250 | 0.000 | 3.541 | 0.000 |
| PGA1_RS04060 | PGA1_c08170 | *cobN* | 1.764 | 0.000 | 1.570 | 0.000 | -3.365 | 0.000 | 0.541 | 0.435 |
| PGA1_RS04860 | PGA1_c09780 | *cobS* | 0.723 | 0.000 | 1.125 | 0.000 | -1.791 | 0.000 | -0.008 | 0.996 |
| PGA1_RS04850 | PGA1_c09770 | *cobT* | 0.717 | 0.000 | 0.062 | 0.928 | -0.624 | 0.171 | 0.121 | 0.990 |
| PGA1_RS04065 | PGA1_c08180 | *cobW* | 1.372 | 0.000 | 1.089 | 0.001 | -4.023 | 0.000 | 0.160 | 0.986 |
| PGA1_RS04070 | PGA1_c08190 | DUF1636 | 1.860 | 0.000 | 0.815 | 0.155 | -3.033 | 0.000 | 0.990 | 0.347 |

**Supplementary Table S2 (Continued)**

| **RefSeq accession** | **Old accession** | **Gene ID** | **Glycine betaine vs. Glucose** | | **Co-culture (growth) vs. Glucose** | | **Co-culture (growth) vs.**  **Co-culture (demise)** | | **Co-culture (day 9) vs.**  **Co-culture (day 4)** | |
| --- | --- | --- | --- | --- | --- | --- | --- | --- | --- | --- |
|  |  |  | **log_2_FC*** | **padj**** | **log_2_FC*** | **padj**** | **log_2_FC*** | **padj**** | **log_2_FC*** | **padj**** |
| PGA1_RS04075 | PGA1_c08200 | *cobO* | 0.593 | 0.000 | 0.055 | 0.950 | -0.503 | 0.440 | -0.003 | 0.997 |
| PGA1_RS00620 | PGA1_c01260 | *pduO* | 0.313 | 0.122 | -0.299 | 0.686 | 0.098 | 0.874 | 1.256 | 0.180 |
| PGA1_RS11940 | PGA1_c24000 | *cobQ/cbiP* | 1.944 | 0.000 | 0.862 | 0.070 | -0.454 | 0.313 | -0.096 | 0.995 |
| PGA1_RS00405 | PGA1_c00830 | *cobC* | 0.275 | 0.010 | -1.277 | 0.024 | 2.224 | 0.000 | -0.194 | 0.946 |
| PGA1_RS00400 | PGA1_c00820 | *cobD/cbiB* | 0.441 | 0.000 | -1.329 | 0.046 | 1.989 | 0.001 | -0.003 | 0.998 |
| PGA1_RS13250 | PGA1_c26670 | *cobP* | 0.562 | 0.000 | -0.380 | 0.667 | 0.507 | 0.478 | -0.125 | 0.945 |
| PGA1_RS00275 | PGA1_c00550 | *bluB* | -0.412 | 0.000 | 0.907 | 0.042 | -0.326 | 0.463 | -0.135 | 0.990 |
| PGA1_RS03760 | PGA1_c07570 | *cobU* | 1.316 | 0.000 | 1.566 | 0.000 | -1.285 | 0.000 | 0.161 | 0.986 |
| PGA1_RS13255 | PGA1_c26680 | *cobC/ phpB* | 0.870 | 0.000 | -0.176 | 0.829 | 0.036 | 0.959 | 0.065 | 0.995 |
| PGA1_RS03755 | PGA1_c07560 | *cobV* | 0.824 | 0.000 | 0.182 | 0.834 | -0.229 | 0.734 | -0.034 | 0.995 |
| PGA1_RS14880 | PGA1_c29940 | *hemE* | 1.088 | 0.000 | 0.197 | 0.757 | -1.089 | 0.015 | 0.338 | 0.792 |
| PGA1_RS00180 | PGA1_c00360 | *hemN* | 3.605 | 0.000 | 0.305 | 0.709 | -0.213 | 0.751 | 4.688 | 0.000 |
| PGA1_RS14895 | PGA1_c29970 | *hemF* | 0.713 | 0.000 | -0.366 | 0.611 | -0.603 | 0.294 | 0.110 | 0.990 |
| PGA1_RS17640 | PGA1_c35520 | *hemY* | 2.184 | 0.000 | -0.275 | 0.546 | 1.277 | 0.000 | 0.136 | 0.990 |
| PGA1_RS17250 | PGA1_c34740 | *hemH* | 0.350 | 0.004 | -1.847 | 0.001 | 1.921 | 0.000 | -0.101 | 0.977 |
| PGA1_RS02820 | PGA1_c05690 | *ctb* | 2.313 | 0.000 | 0.702 | 0.275 | -0.952 | 0.073 | 0.250 | 0.789 |
| PGA1_RS07200 | PGA1_c14480 | *speD* | 2.783 | 0.000 | 4.471 | 0.000 | -7.579 | 0.000 | 0.093 | 0.995 |
| PGA1_RS07195 | PGA1_c14470 | *speE* | 3.277 | 0.000 | 4.242 | 0.000 | -7.177 | 0.000 | 0.326 | 0.807 |
| PGA1_RS08565 | PGA1_c17250 | *gapA1* | 7.100 | 0.000 | 6.204 | 0.000 | -4.508 | 0.000 | -2.838 | 0.004 |
| PGA1_RS02730 | PGA1_c05510 | *gapA2* | 1.827 | 0.000 | -0.049 | 0.958 | 0.515 | 0.457 | 0.052 | 0.995 |
| PGA1_RS19445 | PGA1_78p00290 | *gcvT* | -0.683 | 0.000 | -0.416 | 0.553 | 1.383 | 0.015 | -0.178 | 0.943 |
| PGA1_RS19450 | PGA1_78p00300 | *gcvH1* | -0.753 | 0.000 | -0.701 | 0.202 | 1.236 | 0.002 | 0.034 | 0.995 |
| PGA1_RS02760 | PGA1_c05560 | *gcvH2* | -0.739 | 0.000 | -0.818 | 0.381 | 1.998 | 0.048 | -0.029 | 0.995 |
| PGA1_RS19455 | PGA1_78p00310 | *gcvP1* | -0.803 | 0.000 | -1.630 | 0.000 | 1.469 | 0.000 | 0.193 | 0.950 |

**Supplementary Table S2 (Continued)**

| **RefSeq accession** | **Old accession** | **Gene ID** | **Glycine betaine vs. Glucose** | | **Co-culture (growth) vs. Glucose** | | **Co-culture (growth) vs.**  **Co-culture (demise)** | | **Co-culture (day 9) vs.**  **Co-culture (day 4)** | |
| --- | --- | --- | --- | --- | --- | --- | --- | --- | --- | --- |
|  |  |  | **log_2_FC*** | **padj**** | **log_2_FC*** | **padj**** | **log_2_FC*** | **padj**** | **log_2_FC*** | **padj**** |
| PGA1_RS02765 | PGA1_c05570 | *gcvP2* | -1.197 | 0.000 | -0.592 | 0.503 | 0.972 | 0.222 | 0.156 | 0.780 |
| PGA1_RS12185 | PGA1_c24490 | *mce* | 1.566 | 0.000 | -0.565 | 0.364 | -0.684 | 0.190 | 0.421 | 0.576 |
| PGA1_RS10655 | PGA1_c21510 | *mut* | -0.098 | 0.468 | -0.272 | 0.635 | 0.426 | 0.345 | 0.220 | 0.945 |
| PGA1_RS09040 | PGA1_c18210 | *late control protein D* | 1.184 | 0.001 | -0.144 | NA | 0.354 | 0.570 | 0.491 | 0.026 |
| PGA1_RS09050 | PGA1_c18230 | *tail protein* | 1.479 | 0.026 | -0.203 | NA | 0.178 | NA | NA | NA |
| PGA1_RS09055 | PGA1_c18240 | *tail tape measure p.* | 1.832 | 0.000 | -0.325 | NA | 0.536 | 0.446 | 0.474 | 0.006 |
| PGA1_RS09060 | PGA1_c18250 | *tail assembly protein* | 3.210 | 0.000 | -0.182 | NA | 0.350 | NA | NA | NA |
| PGA1_RS09065 | PGA1_c18260 | *major tail tube protein* | 3.864 | 0.000 | 0.055 | NA | 0.066 | NA | NA | NA |
| PGA1_RS09070 | PGA1_c18270 | *tail sheath* | 3.003 | 0.000 | 0.238 | NA | -0.037 | 0.942 | 0.035 | NA |
| PGA1_RS09095 | PGA1_c18320 | *tail protein* | 1.909 | 0.000 | -0.344 | NA | 0.416 | NA | NA | NA |
| PGA1_RS09100 | PGA1_c18330 | *baseplate assembly p. J* | 2.220 | 0.000 | -0.370 | NA | 0.477 | NA | NA | NA |
| PGA1_RS19860 | PGA1_c18350 | *baseplate assembly p. V* | 0.872 | 0.044 | 0.022 | NA | 0.278 | 0.640 | -0.029 | NA |
| PGA1_RS09140 | PGA1_c18400 | *major capsid protein* | 3.512 | 0.000 | -0.071 | NA | 0.306 | 0.618 | 0.031 | NA |
| PGA1_RS09145 | PGA1_c18410 | *portal protein* | 3.363 | 0.000 | -0.013 | NA | 0.093 | 0.883 | -0.014 | NA |
| PGA1_RS09165 | PGA1_c18450 | *spanin-like protein* | 2.389 | 0.000 | NA | NA | 0.483 | NA | NA | NA |

**Supplementary Table S2 (Continued)**

| **RefSeq accession** | **Old accession** | **Gene ID** | **Glycine betaine vs. Glucose** | | **Co-culture (growth) vs. Glucose** | | **Co-culture (growth) vs.**  **Co-culture (demise)** | | **Co-culture (day 9) vs.**  **Co-culture (day 4)** | |
| --- | --- | --- | --- | --- | --- | --- | --- | --- | --- | --- |
|  |  |  | **log_2_FC*** | **padj**** | **log_2_FC*** | **padj**** | **log_2_FC*** | **padj**** | **log_2_FC*** | **padj**** |
| PGA1_RS00615 | PGA1_c01250 | *fixA* | 2.607 | 0.000 | 1.202 | 0.000 | -1.688 | 0.000 | 0.606 | 0.302 |
| PGA1_RS00610 | PGA1_c01240 | *fixB* | 2.418 | 0.000 | 1.176 | 0.002 | -1.535 | 0.000 | 0.734 | 0.146 |
| PGA1_RS01625 | PGA1_c03320 | *fixC* | 2.278 | 0.000 | 2.371 | 0.000 | -1.888 | 0.000 | 0.044 | 0.995 |
| PGA1_RS14970 | PGA1_c30120 | *ccoN* | 1.772 | 0.000 | -0.285 | 0.694 | -0.468 | 0.410 | 2.319 | 0.000 |
| PGA1_RS14965 | PGA1_c30110 | *ccoO* | 1.665 | 0.000 | -0.265 | 0.702 | -1.127 | 0.037 | 1.900 | 0.000 |
| PGA1_RS14960 | PGA1_c30100 | *ccoQ* | 1.467 | 0.000 | -0.110 | 0.899 | -1.106 | 0.085 | 2.656 | 0.001 |
| PGA1_RS14955 | PGA1_c30090 | *ccoP* | 0.857 | 0.000 | -0.062 | 0.938 | -0.603 | 0.290 | 2.017 | 0.000 |

* log_2_(fold change), ** adjusted *P*-value

**Supplementary Table S3: Distribution of GB demethylation and methionine biosynthesis genes in the pangenome of 40 members of the *Rhodobacterales* and in *Sinorhizobium meliloti*.**

|  | ***gbcA*** | ***gbcB*** | **Cbl ribo-switch*** | ***mtgB1*** | ***mtgC*** | ***mtgD*** | ***mtgE*** | ***metH*** | ***metH1 &***  ***metH2*** | ***metE***** |
| --- | --- | --- | --- | --- | --- | --- | --- | --- | --- | --- |
| *Dinoroseobacter shibae* DFL 12 |  |  |  | + | + | + | + |  |  | + |
| *Jannaschia sp.* CCS1 |  |  |  | + | + | + | + |  |  | + |
| *Leisingera aquaemixtae* R2C4 | + | + | + | + | + | + | + |  |  |  |
| *L. methylohalidivorans* DSM 14336 | + | + | + | + | + | + | + |  |  |  |
| *Leisingera sp.* NJS201 | + | + | + | + | + | + | + |  |  | + |
| *Leisingera sp.* NJS204 | + | + | + | + | + | + | + |  |  |  |
| *Octadecabacter antarcticus* 307 |  |  |  | + | + | + | + |  |  | + |
| *Octadecabacter arcticus* 238 |  |  |  | + | + | + | + |  |  | ++ |
| *Octadecabacter sp.* SW4 |  |  |  | + | + | + | + |  |  | + |
| *Octadecabacter temperatus* SB1 |  |  |  | + | + | + | + |  |  |  |
| *Paracoccus aminophilus* DSM 8538 | + | + |  |  |  |  | + | + |  | + |
| *Paracoccus denitrificans* DSM 413 | ++ | ++ |  |  |  |  | + | + |  | + |
| *Paracoccus pantotrophus* DSM 2944 | ++ | ++ |  |  |  |  | + | + |  |  |
| *Paracoccus sp.* Arc7-R13 | + | + |  |  |  |  | + |  | + |  |
| *Paracoccus tegillarcae* BM15 | + | + |  | + | + | ++ | + |  | + | + |
| *Phaeobacter gallaeciensis* DSM 26640 | + | + | + | + | + | + | + |  |  | + |
| *Phaeobacter gallaeciensis* P11 | + | + | + | + | + | + | + |  |  | + |
| *Phaeobacter gallaeciensis* P128 | + | + | + | + | + | + | + |  |  | + |
| *Phaeobacter gallaeciensis* P63 | + | + | + | + | + | + | + |  |  | + |
| *Phaeobacter inhibens* 2.10 | + | + | + | + | + | + | + |  |  | + |
| *Phaeobacter inhibens* DSM 17395 | + | + | + | + | + | + | + |  |  | + |
| *Phaeobacter inhibens* P66 | + | + | + | + | + | + | + |  |  | + |
| *Phaeobacter inhibens* P70 | + | + | + | + | + | + | + |  |  | + |
| *Phaeobacter piscinae* P18 | + | + | + | + | + | + | + |  |  | + |
| *Phaeobacter piscinae* P36 |  |  |  | + | + | + | + |  |  | + |

**Supplementary Table S3 (Continued)**

| **Strain** | ***gbcA*** | ***gbcB*** | **Cbl ribo-switch*** | ***mtgB1*** | ***mtgC*** | ***mtgD*** | ***mtgE*** | ***metH*** | ***metH1 &***  ***metH2*** | ***metE***** |
| --- | --- | --- | --- | --- | --- | --- | --- | --- | --- | --- |
| *Phaeobacter porticola* P97 |  |  |  | + | + | + | + |  |  | + |
| *Phaeobacter sp.* LSS9 | + | + | + | + | + | + | + |  |  | + |
| *Rhodobacter capsulatus* A12 | + | ++ | + | + | + | + | + |  | + |  |
| *Rhodobacter sp.* CZR27 | + | + |  | + | + | + | + |  |  |  |
| *Rhodobacter xanthinilyticus* LPB0142 | + | + |  | + | + | + | + |  |  |  |
| *Cereibacter sphaeroides* 2.4.1 | + | + |  | + | + | + | + |  | + |  |
| *Cereibacter sphaeroides* ATCC17029 |  |  |  | + | + | + | + |  | + |  |
| *Cereibacter sphaeroides* KD131 | + | + |  | + | + | + | + |  | + |  |
| *Roseobacter denitrificans* Och 114 |  |  |  | + | + | + | + |  | + | + |
| *Roseobacter litoralis* Och 149 |  |  |  | + | + | + | + |  | + | + |
| *Roseobacter ponti* DSM 106830 |  |  |  | + | + | + | + |  |  | + |
| *Ruegeria pomeroyi* DSS-3 |  |  |  | + | + | + | + |  |  |  |
| *Ruegeria* *sp.* AD91A |  |  |  | + | + | + | + |  |  | + |
| *Ruegeria* *sp.* THAF33 | + | + | + | + | + | + | + |  |  |  |
| *Ruegeria sp.* TM1040 | + | + | + | + | + | + | + |  | + |  |
| *Sinorhizobium meliloti* SM11*** | + | + | + | + | + | + | + |  |  |  |
| *Sinorhizobium meliloti* 1021*** | + | + | + | + | + | + | + | + |  |  |
| *Cbl riboswitches in the 5’-UTR of the *gbcAB* transcripts as annotated in the RefSeq database (RFAM: RF00174) using INFERNAL:1.1.5  **This protein is only the C-terminal domain of the *Escherichia coli* MetE and likely not functional as indicated by the methionine auxotrophy of *R. pomeroyi* DSS-3 Δ*mtgC mutan*t^5^ that we also observed for the *P. inhibens* DMS 17395 Δ*mtgC mutan*t  ***Homologs were identified by blastp analysis (Sequence identity > 30% and e-value < e-40) using the *P. inhibens* DSM 17395 proteins as query; *Paracoccus aminophilus* DSM 8538 MetH and *Rhodobacter capsulatus* A12 MetH1 & MetH2 served as query for the genes absent in *P. inhibens*.; best hits for *gbcAB* were subunits of the stachydrine demethylase, but we also found hits for *gbcA* (SM_RS04435, SM11_RS02585) and *gbcB* (SM_RS04440, SM11_RS02590) | | | | | | | | | | |

**Supplementary Table S4: Distribution of Cbl biosynthesis genes in the pangenome of 40 members of the *Rhodobacterales.***

| **Strain** | Number of Cbl biosynthesis genes* | | |
| --- | --- | --- | --- |
|  | 28 (complete) | 27 of 28 | 26 of 28 |
| *Dinoroseobacter shibae* DFL 12 |  |  | + |
| *Jannaschia sp.* CCS1 |  | + |  |
| *Leisingera aquaemixtae* R2C4 | + |  |  |
| *L. methylohalidivorans* DSM 14336 | + |  |  |
| *Leisingera sp.* NJS201 | + |  |  |
| *Leisingera sp.* NJS204 | + |  |  |
| *Octadecabacter antarcticus* 307 |  |  | + |
| *Octadecabacter arcticus* 238 |  |  | + |
| *Octadecabacter sp.* SW4 |  |  | + |
| *Octadecabacter temperatus* SB1 |  |  | + |
| *Paracoccus aminophilus* DSM 8538 | + |  |  |
| *Paracoccus denitrificans* DSM 413 | + |  |  |
| *Paracoccus pantotrophus* DSM 2944 | + |  |  |
| *Paracoccus sp.* Arc7-R13 | + |  |  |
| *Paracoccus tegillarcae* BM15 | + |  |  |
| *Phaeobacter gallaeciensis* DSM 26640 | + |  |  |
| *Phaeobacter gallaeciensis* P11 | + |  |  |
| *Phaeobacter gallaeciensis* P128 | + |  |  |
| *Phaeobacter gallaeciensis* P63 | + |  |  |
| *Phaeobacter inhibens* 2.10 | + |  |  |
| *Phaeobacter inhibens* DSM 17395 | + |  |  |
| *Phaeobacter inhibens* P66 | + |  |  |
| *Phaeobacter inhibens* P70 | + |  |  |
| *Phaeobacter piscinae* P18 | + |  |  |
| *Phaeobacter piscinae* P36 | + |  |  |
| *Phaeobacter porticola* P97 | + |  |  |
| *Phaeobacter sp.* LSS9 | + |  |  |
| *Rhodobacter capsulatus* A12 |  |  | + |
| *Rhodobacter sp.* CZR27 |  | + |  |
| *Rhodobacter xanthinilyticus* LPB0142 | + |  |  |
| *Cereibacter sphaeroides* 2.4.1 |  | + |  |
| *Cereibacter sphaeroides* ATCC17029 |  | + |  |
| *Cereibacter sphaeroides* KD131 |  | + |  |
| *Roseobacter denitrificans* Och 114 | + |  |  |
| *Roseobacter litoralis* Och 149 | + |  |  |
| *Roseobacter ponti* DSM 106830 |  | + |  |
| *Ruegeria pomeroyi* DSS-3 | + |  |  |
| *Ruegeria* *sp.* AD91A |  | + |  |
| *Ruegeria* *sp.* THAF33 | + |  |  |
| *Ruegeria sp.* TM1040 | + |  |  |

*A set of 28 Cbl biosynthesis genes was searched: Supplementary Table S5 (See Supplementary Data S4 for the gene clusters summary of the pangenome)

**Supplementary Table S5: Set of 28 Cbl biosynthesis genes and respective KO annotations that are present in *P. inhibens* DSM17395 and were used to evaluate presence-absence variations within the pangenome of 40 *Rhodobacterales*.**

| **Gene ID** | **KEGG annotation (KO)** |
| --- | --- |
| *hemA* | K00643 |
| *hemB* | K01698 |
| *hemC* | K01749 |
| *hemD* | K01719 |
| *cobA* | K02303 |
| *cobI-cbiL* | K03394 |
| *cobG* | K02229 |
| *cobJ-cbiH* | K13541 |
| *cobM-cbiF* | K05936 |
| *cobF* | K02228 |
| *cobK-cbiJ* | K05895 |
| *cobL-cbiET* | K00595 |
| *cobH-cbiC* | K06042 |
| *cobB-cbiA* | K02224 |
| *cobS* | K09882 |
| *cobT* | K09883 |
| *cobN* | K02230 |
| *cobW* | K02234 |
| *cobA-btuR* | K00798 |
| *pduO* | K00798 |
| *cobQ-cbiP* | K02232 |
| *cobD-cbiB* | K02227 |
| *pduX* | K16651 |
| *cobC* | K02225 |
| *cobP-cobU* | K02231 |
| *cobP-cobV* | K02233 |
| *cobC* | K02226 |
| *bluB* | K04719 |
| *cobU-cobT* | K00768 |

**Supplementary Table S6: Per sample sequencing depth and *P. inhibens* feature counts.**

| **Sequencing sample ID** | **Glucose Replicate 1** | **Glucose Replicate 2** | **Glucose Replicate 3**  **(excluded)** | **Glycine betaine Replicate 1** | **Glycine betaine Replicate 2** | **Glycine**  **betaine Replicate 3** |
| --- | --- | --- | --- | --- | --- | --- |
| Comment |  |  | Wrong barcode assignement |  |  |  |
| Barcode id | BC_13 | BC_14 | BC_18 | BC_16 | BC_17 | BC_15 |
| Barcode | GCCCTCCGT | GTACCGAGT | GTGGCTGCT | AGTTAGAGT | CGATCCAGT | AAGTGCCGT |
| Paired-end reads (after demultiplexing) | 9,894,442 | 7,476,438 | 5,219 | 10,066,284 | 10,705,676 | 7,546,578 |
| Paired-end reads (after cutadapt) | 9,893,675 | 7,475,903 | 5,216 | 10,065,705 | 10,704,979 | 7,546,218 |
| *P. inhibens* feature counts (total) | 9,622,448 | 7,243,662 | 4,150 | 9,835,796 | 10,437,077 | 7,359,083 |
| *P. inhibens* feature counts (non-rRNA) | 9,471,743 | 7,129,556 | 4,049 | 9,723,245 | 10,319,624 | 7,277,772 |

**Supplementary Table S7: Bacterial and algal strains (wild types and mutants) used in this study.**

| **Strain name** | **Source** |
| --- | --- |
| *Escherichia coli* TOP10 | Thermo Scientific |
| *Phaeobacter inhibens* DSM17395*;* wild type | German Collection of Microorganism and Cell Cultures (DSMZ) |
| *Phaeobacter inhibens* DSM17395 Δ*mtgC*::*kan^R^*; knockout mutant | This study |
| *Phaeobacter inhibens* DSM17395 Δ*gbcAB*::*kan^R^*; knockout mutant | This study |
| *Phaeobacter inhibens* DSM17395 Δ*mtgB1*::*gm^R^*; knockout mutant | This study |
| *Phaeobacter inhibens* DSM17395 Δ*mtgB1*::*gm^R^*Δ*gbcAB*::*kan^R^*; double knockout mutant | This study |
| *Emiliania huxleyi* CCMP3266 | Bigelow National Center for Marine Algae and Microbiota (NCMA) |

*gm^R^*, gentamycin resistance; *kan^R^*, kanamycin resistance

**Supplementary Table S8: Plasmids used in this study.**

| **Plasmids** | **Source** |
| --- | --- |
| pDEL(kan); *E. coli* vector (ColE1 ori_Ec_, *kan^R^*) | Haufschild *et al.*, 2025^6^ |
| pDEL(kan)_*mtgC*-updown; pDEL(kan) derivative harbouring the homology arms flanking the gene *mtgC* | This study |
| pDEL(kan)_*gbcAB*-updown; pDEL(kan) derivative harbouring the homology arms flanking the genes *gbcAB* | This study |
| pDN5-Gm-TOPO2; *E. coli* vector (ColE1 ori_Ec_, *gm^R^*) | This study |
| pDN5-*mtgB1*Gm-TOPO2; pDEL(kan) derivative harbouring the homology arms flanking the gene *mtgB1* | This study |

*kan^R^*, kanamycin resistance; *gm^R^*, gentamycin resistance **Supplementary Table S9: Oligonucleotides used in this study.** Restriction sites used for cloning are underlined.

| **Primer name** | **Sequence (5'-3')** | **Source** |
| --- | --- | --- |
| *mtgC* upstream forward primer, *Xho*I site for cloning | TGA**CTCGAG**AGCTAACACTGGAGCGGATCGTTGAGGTGC | This study |
| *mtgC* upstream reverse primer, *Nhe*I site for cloning | TCT**GCTAGC**GATGTCATCTGCGTCTTCCGACATGGGTGTATTCCTTGACTG | This study |
| *mtgC* downstream forward primer, *Xba*I site for cloning | GGA**TCTAGA**AAGCACAACCAGATGAGCGCCTGATTTGGCAC | This study |
| *mtgC* downstream reverse primer, *Hind*III site for cloning | TCT**AAGCTT**GTTTTTGACATGAGGATTCATACCCGCAACATCCTGACAGGAATTG | This study |
| upstream check primer for *mtgC* | AATCATTGGTGGCTGCTGTGGCACCATGC | This study |
| downstream check primer for *mtgC* | GAAACCTCCATCGCCGTGGCAAGC | This study |
| *gbcAB* upstream forward primer, *Xho*I site for cloning | ACT**CTCGAG**AGTGCCAGAACGGACACCCATGCCCAGCGTTTAAC | This study |
| *gbcAB* upstream reverse primer, *Kpn*I site for cloning | TCT**GGTACC**CAAGATCTCGCTGTCGTGGAGCATAAGCACCCCTCTTTG | This study |
| *gbcAB* downstream forward primer, *Eco*RI site for cloning | TCT**GAATTC**GGCAGTGTTTCGGTTGAGATCTAAATGAATATGGGAGGCTCACATTTG | This study |
| *gbcAB* downstream reverse primer, *Hind*III site for cloning | TCT**AAGCTT**TCATAGCTGTATACAAATGTTTGGGTGGCGGAGACGAAGGGATTC | This study |
| upstream check primer for *gbcAB* | GCAATCAGCAGTGTTGCAATCAAAAAATGCAGCCCAC | This study |
| downstream check primer for *gbcAB* | TTACCACCACCTGCAAGTCACGAATAGCGCTCAG | This study |
| Kanamycin cassette verification forward primer | CAGGGCGGGGCGTAACCGATTCGCAGCGCATCGCCTTCTA | This study |
| Kanamycin cassette verification reverse primer | GATTTTTTTCTCCATGCGAAACGATCCTCATCCTGTCTCTTG | This study |

**Supplementary Table S9 (Continued)**

| **Primer name** | **Sequence (5'-3')** | **Source** |
| --- | --- | --- |
| *mtgB1* upstream forward primer with homology to pCII-TOPO | GTAACGGCCGCCAGTGTGCTGGAATTCGCCCTTGCGCAACAGGCCATGGTCCAC | This study |
| *mtgB1* upstream reverse primer with homology to gentamycin cassette | GCATTACAGTTTACGAACCGAACAGGCTTATGTCAAGGTTTTTCCTCACCACAGGCATTTCG | This study |
| *mtgB1* downstream forward primer with homology to gentamycin cassette | GGTGGGCTGCCCTTCCTGGTTGGCTTGGTTTCCAGCGGCGGTTCTTATCTGAACAG | This study |
| *mtgB1* downstream reverse primer with homology to pCII-TOPO | CCGCCAGTGTGATGGATATCTGCAGAATTCGCCCTTCCCGCACGACCACCGCCTTC | This study |
| Gentamycin cassette forward primer | GTTGACATAAGCCTGTTCG | Sperfeld & Narváez-Barragán, *et al*., 2024^7^ |
| Gentamycin cassette reverse primer | GTTAGGTGGCGGTACTTGG | Sperfeld & Narváez-Barragán, *et al*., 2024^7^ |
| *mtgB1* KO verification forward primer | GGACGCCATCGCGCTGAAC | This study |
| *mtgB1* KO verification reverse primer | GGCTGGCTTCTCATGATCTCGC | This study |
| Gentamycin cassette verification forward primer | GTGCAAGCAGATTACGGTGACG | Sperfeld & Narváez-Barragán, *et al*., 2024^7^ |
| Gentamycin cassette verification reverse primer | GAGCCTACATGTGCGAATGATGC | Sperfeld & Narváez-Barragán, *et al*., 2024^7^ |

**Supplementary Table S10: Compatible solutes used in this study.**

| **Osmolyte** | **Manufacturer** | **CAS number** |
| --- | --- | --- |
| Betaine monohydrate | Sigma-Aldrich (Steinheim, Germany) | 590-47-6 |
| L-carnitine hydrochloride | Tokyo Chemical Industry (Tokyo, Japan) | 6645-46-1 |
| DMSP chloride | Tokyo Chemical Industry (Tokyo, Japan) | 4337-33-1 |
| Trigonelline hydrochloride | Sigma-Aldrich (Steinheim, Germany) | 6138-41-6 |

**Supplementary Table S11:** Overview of the 40 members of the *Rhodobacterales* included in the pangenomic analysis and their genome accession numbers.

| **Isolate** | **Habitat** | **Isolation source** | **NCBI RefSeq assembly accession number** |
| --- | --- | --- | --- |
| *Dinoroseobacter shibae* DFL 12 | marine | dinoflagellate, *Prorocentrum lima* | GCF_000018145.1 |
| *Jannaschia sp.* CCS1 | marine | coastal seawater | GCF_000013565.1 |
| *Leisingera aquaemixtae* R2C4 | marine | marine biofilm | GCF_006740765.1 |
| *L. methylohalidivorans* DSM 14336 | marine | seawater collected from a tide pool | GCF_000511355.1 |
| *Leisingera sp.* NJS201 | marine | marine Biofilm | GCF_004535885.1 |
| *Leisingera sp.* NJS204 | marine | marine Biofilm | GCF_004123675.1 |
| *Octadecabacter antarcticus* 307 | marine | ice | GCF_000155675.2 |
| *Octadecabacter arcticus* 238 | marine | ice | GCF_000155735.2 |
| *Octadecabacter sp.* SW4 | marine | seawater | GCF_008065155.1 |
| *Octadecabacter temperatus* SB1 | marine | seawater | GCF_001187845.1 |
| *Paracoccus aminophilus* DSM 8538 | terrestrial | soil | GCF_000444995.1 |
| *Paracoccus denitrificans* DSM 413 | terrestrial | soil | GCF_004063735.1 |
| *Paracoccus pantotrophus* DSM 2944 | terrestrial | denitrifying, sulfide-oxidizing effluent-treatment plant | GCF_008824185.1 |
| *Paracoccus sp.* Arc7-R13 | marine | arctic ocean sediment, 267 m depth | GCF_004010775.1 |
| *Paracoccus tegillarcae* BM15 | marine | clam, *Tegillarca granosa* | GCF_002847305.1 |
| *Phaeobacter gallaeciensis* DSM 26640 | marine | clam larvae aquaculture, *Pecten maximus* | GCF_000511385.1 |
| *Phaeobacter gallaeciensis* P11 | marine | clam larvae aquaculture, *Pecten maximus* | GCF_002381305.1 |
| *Phaeobacter gallaeciensis* P128 | marine | zooplankton from North Pacific | GCF_002381365.1 |
| *Phaeobacter gallaeciensis* P63 | marine | algae aquaculture, diatom | GCF_002393525.1 |
| *Phaeobacter inhibens* 2.10 | marine | algae, *Ulva lactuca* | GCF_000154745.2 |
| ***Phaeobacter inhibens* DSM 17395*** | marine | clam larvae aquaculture, *Pecten maximus* | GCF_000154765.2 |
| *Phaeobacter inhibens* P66 | marine | algae aquaculture, phytoplankton mixture | GCF_002892065.1 |
| *Phaeobacter inhibens* P70 | marine | algae aquaculture, phytoplankton mixture | GCF_002892125.1 |
| *Phaeobacter piscinae* P18 | marine | fish larvae aquaculture, *Scophthalmus maximus* | GCF_002891925.1 |
| *Phaeobacter piscinae* P36 | marine | fish aquaculture, *Scophthalmus maximus* | GCF_002412025.1 |
| *Phaeobacter porticola* P97 | marine | barnacle | GCF_001888185.1 |

**Supplementary Table S11 (Continued)**

| **Isolate** | **Habitat** | **Isolation source** | **NCBI Ref Seq assembly**  **accession number** |
| --- | --- | --- | --- |
| *Phaeobacter sp.* LSS9 | marine | red seaweed *Delisea pulchra* | GCF_003443595.1 |
| *Rhodobacter capsulatus* A12 | terrestrial | soil | GCF_014622665.1 |
| *Rhodobacter sp.* CZR27 | terrestrial | soil | GCF_002407205.1 |
| *Rhodobacter xanthinilyticus* LPB0142 | marine | seawater | GCF_001856665.1 |
| *Cereibacter sphaeroides* 2.4.1** | unknown | unknown | GCF_003324715.1 |
| *Cereibacter sphaeroides* ATCC17029** | unknown | unknown | GCF_000015985.1 |
| *Cereibacter sphaeroides* KD131** | marine | sea mud | GCF_000021005.1 |
| *Roseobacter denitrificans* Och 114 | marine | seaweed *Enteromorpha linza* (coastal marine sediments) | GCF_002983865.1 |
| *Roseobacter litoralis* Och 149 | marine | seaweed | GCF_000154785.2 |
| *Roseobacter ponti* DSM 106830 | marine | seawater | GCF_012932215.1 |
| *Ruegeria pomeroyi* DSS-3 | marine | seawater | GCF_000011965.2 |
| *Ruegeria* *sp.* AD91A | marine | red seaweed *Delisea pulchra* | GCF_003443535.1 |
| *Ruegeria* *sp.* THAF33 | marine | surface of a polyethylene microplastic particle | GCF_009363615.1 |
| *Ruegeria sp.* TM1040 | marine | marine dinoflagellate culture (*Pfiesteria piscicida*) | GCF_000014065.1 |

*Laboratory strain of this study, ***Rhodobacter sphaeroides*

**Supplementary Table S12: Pfams, COG and KO annotations of GB demethylases and Cbl biosynthesis genes.**

| **Gene name** | **Pfams** | **COG** | **KO** |
| --- | --- | --- | --- |
| *mtgE* | PF02574.18 | COG0646 | K00548 |
| *mtgB* | PF06253.13 | COG5598 | K14083 |
| *mtgC* | PF02310.21 | COG5012 | NA |
| *mtgD'* | PF00809.24 | COG1410 | NA |
| *gbcA* | PF00848.21 | COG4638 | K24003 |
| *gbcB* | PF00175.23 | COG0633\|COG1018 | K24004 |
| *cobA1* | PF00590 | COG0007 | K02303 |
| *cbiM* | PF01891.18 | COG0310 | K02007 |
| *cobV* | PF02654.17 | COG0368 | K02233 |
| *cobJ/cbiH* | PF01890.18 | COG1010 | K13541\|K05934 |
| *cobD* | PF03186.15 | COG1270 | K02227 |
| *cobN* | PF02514.18 | COG1429 | K02230 |
| *cobQ/cbiP* | PF07685.16 | COG1492 | K02232 |
| *cobB/cbiA* | PF07685.16 | COG1797 | K02224 |
| *cbiD* | PF01888.19 | COG1903 | K02188 |
| *cbiG* | PF01890.18 | COG2073 | K02189 |
| *cobH/cbiA* | PF02570.17 | COG2082 | K06042 |
| *cobP/cobU* | PF02283.18 | COG2087 | K02231 |
| *cobK/cbiJ* | PF02571.16 | COG2099 | K05895 |
| *pduO* | PF01923.20 | COG2109\|COG2096 | K00798 |
| *cobL/cbiET* | PF00590.22 | COG2241 | K00595 |
| *cobI/cbiL* | PF00590.22 | COG2243 | K03394 |

**Supplementary Data S1:** *P. inhibens* feature table (genes) with RNA-sequencing results from exconentally growing cultures with glycine betaine or glucose. The dataset includes bacterial gene accession numbers, functional annotations, transcript abundances (TPM normalized read counts), results of DESeq2 differential gene expression analysis and lists of the top 100 upregulated genes ranked either by TPM or log_2_(fold change) (Fig. 1, Supplementary Table S2).

**Supplementary Data S2:** *P. inhibens* feature table (genes) with results of DESeq2 differential gene expression analysis using previously published data for co-cultures on day 4 and day 9 of the growth phase. The dataset also includes bacterial gene accession numbers, functional annotations and a list of upregulated genes (Fig. 2, Supplementary Table S2).

**Supplementary Data S3:** *P. inhibens* feature table (genes) with results of DESeq2 differential gene expression analysis using previously published data for co-cultures during growth and demise phase. The dataset also includes bacterial gene accession numbers and functional annotations (Fig. 1, Supplementary Table S2).

**Supplementary Data S4:** Gene clusters summary file of the pangenome of 40 *Rhodobacterales*. The dataset includes gene cluster and genome identifier, functional annotations as well as amino acid sequences. (Supplementary Table S3 and S4)

**Supplementary Data S5:** Prevalence of GB metabolism genes across 15,525 marine bacterial genomes. The analyzed genomes are a subset of the Ocean Microbiomics Database (OMDB), including only high-quality genomes that are representatives of species-level operational taxonomic units (mOTUs). The dataset includes OMDB genome identifiers, presence-absence values for GB demethylases and Cbl biosynthesis genes, and GTDB taxonomic assignments. (Fig. 6 and Supplementary Fig. S3)
